# Mechanisms underpinning osteosarcoma genome complexity and evolution

**DOI:** 10.1101/2023.12.29.573403

**Authors:** Jose Espejo Valle-Inclan, Solange De Noon, Katherine Trevers, Hillary Elrick, Mélanie Tanguy, Thomas Butters, Francesc Muyas, Fernanda Amary, Roberto Tirabosco, Adam Giess, Alona Sosinky, Greg Elgar, Adrienne M. Flanagan, Isidro Cortés-Ciriano

**Author notes:** These authors contributed equally to this work (Jose Espejo Valle-Inclan, Solange De Noon).

## Abstract

Osteosarcoma is the most common primary cancer of bone with a peak incidence in children and young adults. Despite progress, the genomic aberrations underpinning osteosarcoma evolution remain poorly understood. Using multi-region whole-genome sequencing, we find that chromothripsis is an ongoing mutational process, occurring subclonally in 74% of tumours. Chromothripsis drives the acquisition of oncogenic mutations and generates highly unstable derivative chromosomes, the evolution of which drives clonal diversification and intra-tumour heterogeneity. In addition, we report a novel mechanism, loss-translocation-amplification (LTA) chromothripsis, which mediates rapid malignant transformation and punctuated evolution in about half of paediatric and adult high-grade osteosarcomas. Specifically, a single double-strand break triggers concomitant *TP53* inactivation and segmental amplifications, often amplifying oncogenes to high copy numbers in extrachromosomal circular DNA elements through breakage-fusion-bridge cycles involving multiple chromosomes. LTA chromothripsis is detected at low frequency in soft-tissue sarcomas, but not in epithelial cancers, including those driven by *TP53* mutation. Finally, we identify genome-wide loss of heterozygosity as a strong prognostic indicator for high-grade osteosarcoma.

## Introduction

Whole-genome sequencing (WGS) studies of tumours has revealed that most human cancers are riddled by complex forms of structural variation(*1*, *2*). A major mutational process driving cancer genome complexity is chromothripsis, which refers to the acquisition of hundreds to thousands of clustered rearrangements in one or a few chromosomes(*3–5*) resulting from chromosomal shattering in micronuclei(*6*) or through fragmentation of dicentric chromosomes during anaphase bridge resolution(*7–9*). Chromothripsis is pervasive across diverse cancer types(*10*, *11*), and often mediates malignant transformation(*10*, *12–14*) and drug resistance(*12*, *15*). A remarkable example of a cancer characterised by chromothripsis is osteosarcoma, the most common primary malignancy of bone for which therapeutic options and survival rates have not improved for over four decades(*16*). Although the remarkable karyotypic complexity of osteosarcoma has long been established(*17–21*), the mechanisms underpinning osteosarcoma genome complexity and the impact of chromothripsis during tumour evolution remain elusive.

Chromothripsis can be identified by the presence of rearrangement profiles characterised by interleaved structural variants (SVs) with variable levels of loss of heterozygosity (LOH), which reflect the random rejoining of DNA fragments following chromosome fragmentation(*3*, *10*, *22*). However, in osteosarcoma, chromothripsis events remarkably often involve >10 chromosomes and co-localize with other genomic aberrations, such as whole-genome doubling (WGD) and segmental amplifications(*10*, *19*), the acquisition of which cannot be fully explained by rearrangement mechanisms already described(*10*).

Here, through the analysis of multi-region long and short-read WGS data, we show that chromothripsis is an ongoing mutational process throughout osteosarcoma evolution and drives remarkable karyotypic heterogeneity. In fact, we detect subclonal chromothripsis events in 74% of tumours. In addition, we report a novel mechanism, loss-translocation-amplification (LTA) chromothripsis, which mediates *TP53* loss, oncogene amplification and WGD in about half of both paediatric and adult high-grade osteosarcomas, leading to malignant transformation and punctuated evolution over the course of a few cell divisions. LTA chromothripsis leads to the formation of highly unstable derivative chromosomes, the evolution of which drives the formation of the most complex chromothripsis events detected in human cancers, often involving >15 chromosomes. We find that LTA chromothripsis also occurs in other sarcoma subtypes but not across >2,600 tumours spanning 31 cancer types, including those driven by *TP53* inactivation, such as ovarian and oesophageal adenocarcinoma. Our findings suggest that LTA is a mechanism specific to the evolution of high-grade osteosarcoma, and to a lesser extent, other sarcomas.

## Results

### Clinical samples and whole-genome sequencing data

We processed WGS data uniformly for 356 primary and metastatic osteosarcoma samples from 197 patients, including published data for 128 tumour samples(*19*, *21*, *23*, *24*) (**Fig. 1A, fig. S1 and Methods**). New data include high-depth multi-region WGS for 228 tumour regions from 85 patients (2-8 regions per tumour; median 4 regions) generated as part of the Genomics England 100,000 Genomes Project (G100k). Additionally, we sequenced 13 samples from uncommon histological variants of primary bone sarcomas (**Methods and Tables S1 and S2**). The final data set included 246 high-grade conventional (HGOS), 43 parosteal, 15 periosteal, 9 low-grade central, and 30 soft-tissue extraskeletal osteosarcomas, as well as 13 osteoclast-rich primary high-grade bone sarcomas (**fig. S1, Tables S1 and S2, and Methods**). The average patient age was 30 years (range 4 to 89 years; 40% of patients under 18 years old), and 40% of samples were from tumours treated with the MAP chemotherapy protocol (methotrexate, doxorubicin, and cisplatin) or variations thereof (**Table S1 and Methods**).

**Fig. 1.**
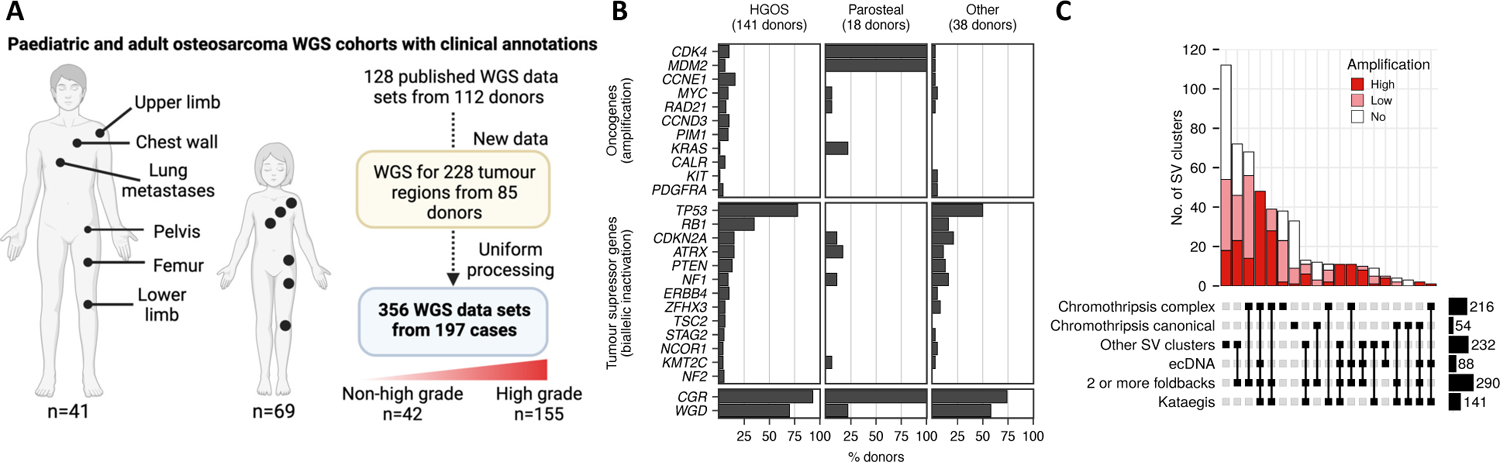
Overview of the study. **(A)** Overview of the data sets and types of osteosarcoma samples analysed in this study. **(B)** Frequency of oncogene amplification, biallelic inactivation of tumour suppressor genes, CGR: complex genome rearrangements (CGR), and whole-genome doubling (WGD) in high-grade osteosarcomas (HGOS), parosteal osteosarcomas and other osteosarcoma subtypes. **(C)** Upset plot showing the overlap of different types of complex rearrangement patterns in the SV clusters detected by ShatterSeek across all osteosarcomas in the cohort. The category “Other SV clusters” encompass complex clusters of SVs without copy-number oscillations. The SV clusters in more than one chromosome were considered a single cluster if linked by translocations. ecDNA: extrachromosomal circular DNA; WGS: whole-genome sequencing.

### Clonal evolution in osteosarcoma is driven by complex genomic rearrangements

Driver gene analysis identified genes known to be involved in osteosarcoma evolution, including *TP53* (altered in 136/197 tumours, 69%, primarily by rearrangements in intron 1), *RB1* (58/197, 29%), and *CDKN2A* (32/197, 16%) (**Fig. 1B, fig. S1, Table S3 and Methods**). We estimated an average burden of 1.85 SNVs per Mb (**fig. S1**), which is consistent with the low burden of point mutations previously reported for osteosarcoma(*14*, *17*, *19*). However, kataegis events were detected in 74% of tumours (145/197), indicating punctuated bursts of mutation accumulation (**Fig. 1C, fig. S1 and Methods**). We detected genomic features associated with activation of the alternative lengthening of telomeres (ALT) pathway in 53% (104/197) tumours, suggesting a predominant role of this pathway in driving replicative immortality in osteosarcoma(*25*, *26*) (**fig. S1, Table S1 and Methods**). Among HGOS, 15% did not harbour alterations in *TP53* pathway genes (*i.e.*, *TP53* and *MDM2*). These cases did not show an enrichment of mutations in other driver alterations compared with *TP53*-mutant HGOS. On average, 55% of SNVs were subclonal (median: 53.4%), indicating high levels of intra-tumour heterogeneity. However, most driver mutations were clonal (**fig. S2**), and we did not detect positive selection in subclonal mutations (**fig. S3 and Methods**), indicating neutral subclonal evolution at the level of point mutations. *De novo* mutational signature analysis of single-base substitutions (SBS) revealed that all signatures detected showed a comparable contribution to clonal and subclonal mutations, except for those associated with platinum therapy (SBS31 and SBS35)(*20*)(**fig. S2 and Table S4**). Platinum-associated signatures revealed clonal sweeps driven by single chemoresistant clones in relapse and metastatic samples, and the survival and outgrowth of multiple subclones in primary tumours upon treatment (**fig. S4**).

In contrast to SNVs, the SV burden was high, with a median of 414 SVs per tumour (**fig. S1**). The majority of SVs mapped to rearrangement clusters showing an equal distribution of SV types, a hallmark of chromothripsis(*3–5*), templated insertions and microhomology tracts at the breakpoints (**fig. S5**). However, only 11% (54/502) of SV clusters were classified as canonical chromothripsis(*10*) (**Methods**). The remainder co-localized with other events, such as extrachromosomal circular DNA (ecDNA) and segmental amplifications (**Fig. 1C, fig. S1. Table S5 and Methods**), which masked the oscillating copy-number pattern characteristic of canonical chromothripsis. Thus, we collectively refer to such complex SV clusters as complex genomic rearrangements (CGRs). HGOS tumours showed high rates of WGD (99/141 tumours; 70%) and CGRs (131/141 tumours; 93%). High-level oncogene amplification often occurred in the context of CGRs (**Fig. 1C**). In parosteal osteosarcomas, co-amplification of *MDM2* and *CDK4* mapped to ecDNA events in all cases (**fig. S1**). In HGOS, 57% (81/141) of tumours harboured at least one high-level oncogene amplification, which were significantly enriched in chromosome arms with CGRs and associated with foldback inversions(*27*) (**Fig. 1C and fig. S1**), indicating that oncogene amplification in HGOS is driven by breakage-fusion-bridge (BFB) cycles.

### Evolutionary trajectories of rearranged chromosomes

To characterise the contribution of chromothripsis to clonal and genome evolution in osteosarcoma, we constructed tumour phylogenies using multi-region WGS data from 46 tumours and WGS data from multiple tumour samples from 10 individuals (**Fig. 2A and Methods**). We classified SVs as clonal, subclonal and private based on whether they were detected in all, a subset or just one of the tumour regions analysed, respectively. We found a high frequency of subclonal CGRs in high, intermediate (periosteal) and low-grade osteosarcoma subtypes, with 62% (130/210) of tumour regions and 74% (40/54) of tumours showing subclonal chromothripsis, often leading to highly divergent karyotypes (**Fig. 2B, figs. S6 to S9, and Table S6**). Thus, these results indicate that chromothripsis is a common subclonal mutational process.

**Fig. 2.**
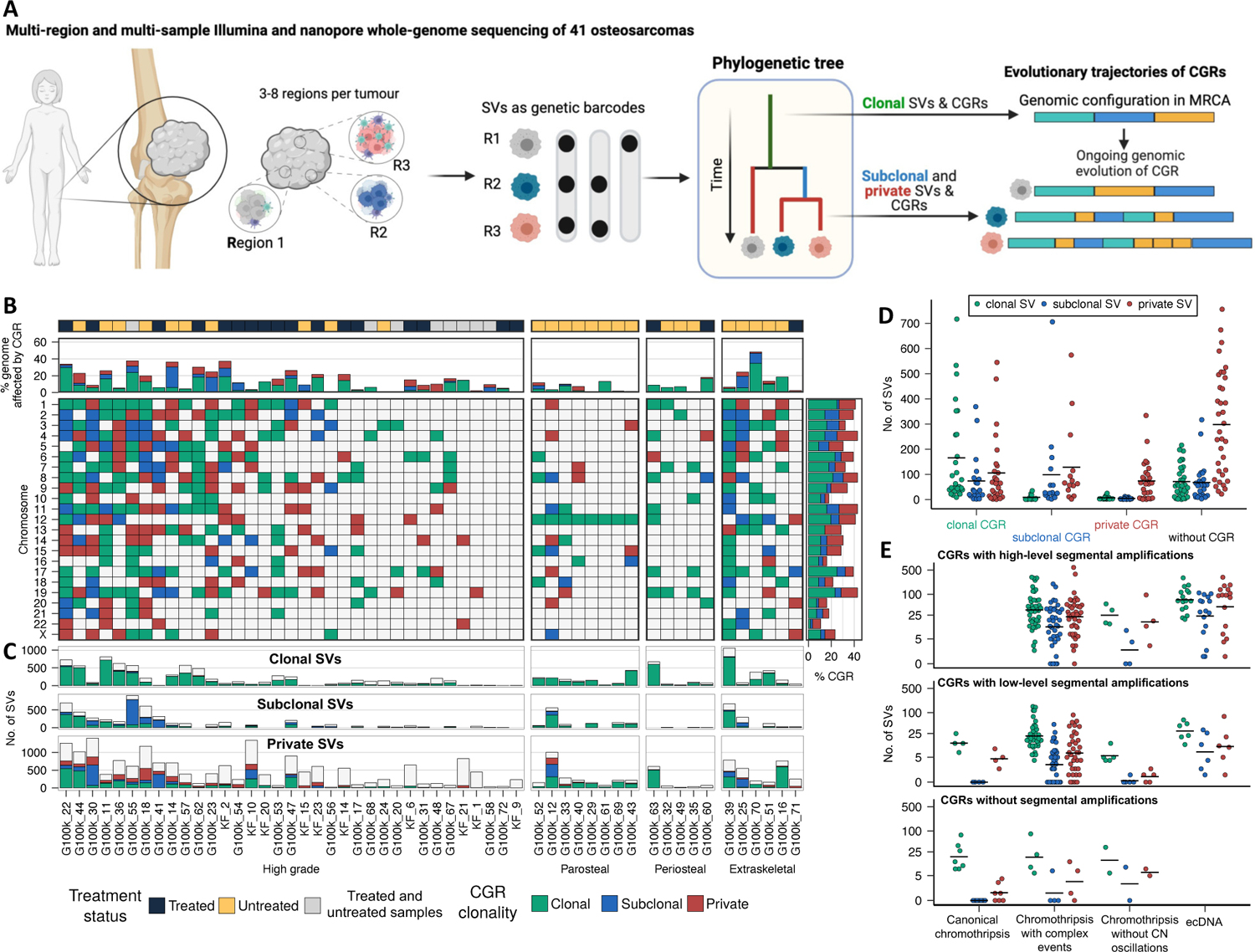
Intra-tumor heterogeneity and evolutionary trajectories of chromothriptic chromosomes. **(A)** Schematic representation of the experimental design based on multi-region whole-genome sequencing used to characterize the patterns of intra-tumour heterogeneity and clonal evolution in osteosarcomas. **(B)** Distribution of clonal (green), subclonal (blue), and private (red) CGRs across autosomes and chromosome X. Each column represents an osteosarcoma with WGS data for at least 3 tumour regions. **(C)** Number of clonal, subclonal and private SVs in each tumour mapping to genomic regions overlapping clonal (green), subclonal (blue), or private CGRs (red). The number of SVs not overlapping regions affected by CGRs is shown in white. **(D)** Number of clonal (green), subclonal (blue) and private (red) SVs mapping to regions affected by clonal, subclonal and private CGRs, as well as outside regions affected by CGRs. Each dot represents a tumour with multi-region WGS data. **(E)** Number of clonal (green), subclonal (blue) and private (red) SVs mapping to regions with canonical chromothripsis, chromothripsis with other complex events, chromothripsis events without copy-number oscillations, and ecDNA stratified based on the presence of high-level, low-level or no segmental amplifications in the SV cluster. CGR: complex genome rearrangement; CN: copy-number; Mixed: individuals with samples collected before and after chemotherapy treatment.

Previous studies suggest that chromothripsis can precede or follow other mutational processes(*7*, *8*), and that rearranged chromosomes are subject to ongoing genomic instability(*9*, *28*). However, studying the evolutionary trajectories of rearranged chromosomes has been challenging due to the limited sensitivity for subclonal SV detection using bulk WGS data and the lack of multi-region or single-cell WGS data. Here, to investigate the evolutionary trajectories of chromosomes affected by chromothripsis, we mapped the rate of subclonal SVs across the genome using the multi-region WGS data. This analysis revealed that genomic regions with clonal CGRs are significantly enriched for subclonal SVs (*P* < 0.001; Fisher’s exact test), indicating that CGRs prime the cancer genome for ongoing chromosomal instability (**Fig. 2C and figs. S10 to S12**). Canonical chromothripsis events do not acquire subclonal or private SVs (**Fig. 2D**). However, we detected hundreds of subclonal and private SVs in genomic regions affected by clonal chromothripsis with segmental amplifications, even when the clonal chromothripsis event only affected a small region of the genome (**Fig. 2C-D and fig. S10**). High-level amplifications showed marked heterogeneity caused by subclonal rearrangements, including translocations to multiple chromosomes, as observed in other ecDNA-driven cancers(*29*, *30*). Together, these results show that derivative chromosomes resulting from chromothripsis events are unstable and evolve through complex rearrangement processes, thereby driving intra-tumour heterogeneity and clonal diversification.

### Loss-Translocation-Amplification (LTA) chromothripsis triggers punctuated tumour evolution

Given the high burden of SVs in HGOS (**fig. S1**), we next sought to identify genomic regions recurrently affected by breakpoints (**Methods**). Consistent with previous studies(*17*, *31*, *32*), the *TP53* locus was significantly rearranged in both adult and paediatric HGOS (**Fig. 3A and fig. S13A**). Investigation of the rearrangement profiles in chromosome 17 revealed frequent co-occurrence of terminal loss of 17p, including the *TP53* locus, and segmental amplification of 17p downstream of *TP53* together with other genomic regions across one or multiple chromosomes, with or without WGD (**Fig. 3A and figs S11 to S17**). Analysis of tumour phylogenies reconstructed using SNVs and SVs detected in multi-region WGS data revealed that a single double-strand break in chromosome 17p (commonly in intron 1 of *TP53*, **fig. S1C**) generates an unprotected terminal end that translocates to another chromosome or its sister chromatid to form a dicentric chromosome, the resolution of which triggers a series of BFB cycles during mitosis that generate segmental amplifications (**Fig. 3B, fig. S18 to S20, and Methods**). As a result, genomic regions and oncogenes from multiple chromosomes can be amplified as part of this process (**Fig. 3E and figs. S11 to S17**). We detected this rearrangement pattern, which we term loss-translocation-amplification (LTA) chromothripsis, in 48% (68/141) of HGOS, including paediatric and adult HGOS in all cohorts analysed (**Fig. 3A-B, figs S1 and S13 to S17**), which we validated using long-read nanopore WGS of selected cases (**fig. S18**). Specifically, we detected LTA chromothripsis in 48% (48/99) of HGOS with WGD, and in 48% of HGOS cases without WGD (20/42). In 66% (45/68) of HGOS with LTA chromothripsis (32% of all HGOS, 45/141), the amplified regions encompassed oncogenes, often in the form of ecDNA, indicating that LTA chromothripsis is a frequent mechanism of oncogene amplification in HGOS (**Fig. 3E and figs. S1, S11 to S17**). In the other 23 cases, the complexity of the derivative chromosomes was comparable, but the segmental amplifications caused by LTA did not encompass recurrently amplified oncogenes (**figs. S19 to S20**). LTA chromothripsis occurred in cells with one *TP53* allele already inactivated by somatic point mutations (24%, 16/68), somatic rearrangements (71%, 48/68), or germline pathogenic variants (6%, 4/68; 4/6 tumours from Li-Fraumeni individuals) (**fig. S1**). Notably, the SV burden and the number of chromosomes involved in chromothripsis are significantly higher in cases with LTA chromothripsis (**Fig. 3C-D**), indicating that the most complex chromothripsis events in osteosarcomas are caused by LTA chromothripsis.

**Fig. 3.**
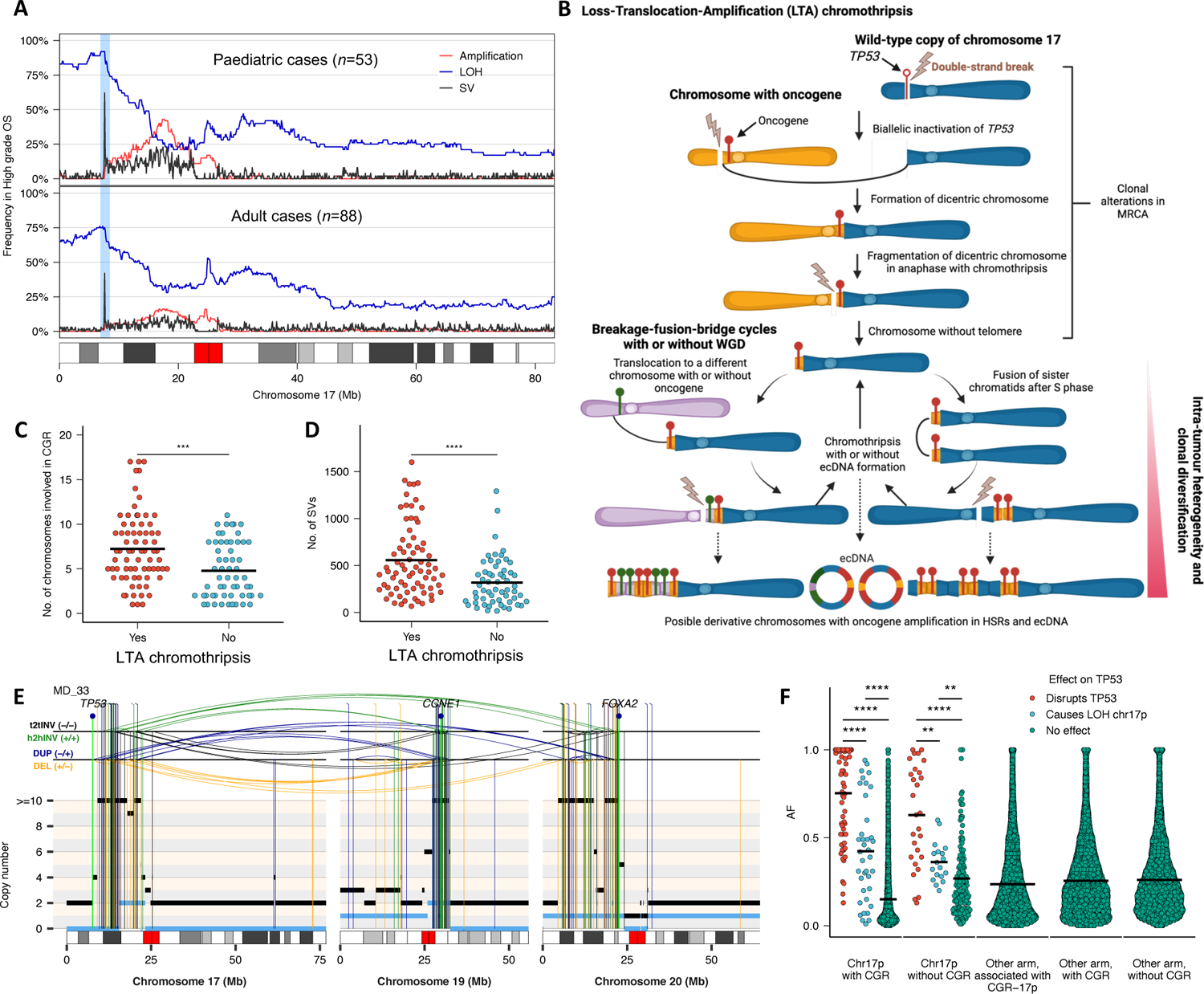
Mechanism of Loss-Translocation-Amplification (LTA) chromothripsis in high-grade osteosarcoma. **(A)** Analysis of the fraction of paediatric (individuals younger than 18 years old) and adult osteosarcomas showing amplification (red), LOH (blue) and SVs (black) across chromosome 17. The frequency of amplification, LOH and breakpoints were computed using non-overlapping windows of 100 kilobase pairs. Only one SV per window and patient was considered for this analysis. **(B)** Schematic representation of the events that lead to the loss of *TP53* and oncogene amplification observed in high-grade osteosarcomas. **(C)** Number of chromosomes involved in CGRs in high-grade osteosarcomas stratified based on the presence a CGR on chromosome 17p triggered by a breakpoint in the *TP53* locus. **(D)** Number of SVs detected across the genome in high-grade osteosarcomas stratified based on the presence a CGR on chromosome 17p triggered by a breakpoint in the *TP53* locus. **(E)** Representative rearrangement profiles for an osteosarcoma in which inactivation of *TP53* by a double-strand break leads to the amplification of multiple oncogenes via LTA chromothripsis. **(F)** Allele fraction (AF) of rearrangements mapping to the CGR detected in 17p in tumours with or without LTA, in other chromosome arms linked to 17p via LTA, chromosome arms with CGR not involved in LTA chromothripsis, and in chromosome arms without CGRs. The total and minor allele copy-number data in **F** are represented in black and blue, respectively. DEL, deletion-like rearrangement; DUP, duplication-like rearrangement; h2hINV, head-to-head inversion; t2tINV, tail-to-tail inversion. Significance in **C**, **D** and **F** was assessed using the two-sided Wilcoxon’s rank test (\*\**P* < 0.001; \*\*\**P* < 0.0001; \*\*\*\**P* < 0.00001).

The patterns of clonal dynamics in HGOS with LTA chromothripsis inferred from the tumour phylogenies are consistent with a model of punctuated evolution. Specifically, macroscopically distant tumour regions mapping centimetres apart from each other (see **figs. S21 to S23** for an example) only share a small set of point mutations and the rearrangements that initiated LTA chromothripsis, which show high allele fraction values (**Fig. 3F**). The lack of shared mutations across regions that harbour tens to hundreds of private SNVs and SVs indicates that clonal diversification occurred rapidly, probably within a few cell divisions. Thus, the genomic instability cascade triggered by LTA chromothripsis following a single double-strand break in the *TP53* locus prompts malignant transformation within a few cell divisions.

We next investigated the frequency of LTA chromothripsis in other sarcomas and common cancers using WGS data from ∼2,800 tumours from the TCGA sarcoma cohort (TCGA-SARC) and the Pan-Cancer Analysis of Whole Genomes (PCAWG) project. We did not detect evidence of LTA in epithelial cancers, including cancers with frequent *TP53* mutation, such as ovarian and oesophageal cancer (**figs. S24 to S27**). By contrast, we detected LTA chromothripsis at low frequency in soft-tissue sarcomas (**figs. S28 to S29**). Together, this indicates that LTA chromothripsis is rare in cancers other than osteosarcoma.

### Whole-genome doubling is a late clonal event that triggers genomic instability

Motivated by the high frequency of WGD in osteosarcoma (**Fig. 1B and fig. S1**), we next sought to determine the timing and relative order of genomic aberrations acquired throughout tumour evolution. The timing of genomic aberrations relative to WGD can be estimated by counting the fraction of somatic SNVs acquired before WGD, which should be present in at least two chromosomal copies (**Methods**). Due to low SNV burdens (**fig. S1**), only 44% (55/125) of tumours with WGD had sufficient clock-like mutations for WGD timing analysis (**Methods**). WGD, which we determined computationally and experimentally (Table **S7 and Methods**), was a late event in 96% (53/55) of tumours, with WGD occurring on average at 80% of mutation time (**Fig. 5A**). No significant differences in WGD timing were observed between HGOS and other osteosarcoma subtypes (**Fig. 5B**). However, both the timing of WGD, and the latency between WGD and the expansion of the MRCA were significantly different between paediatric and adult HGOS cases (**Fig. 5C-D and Table S8**). We estimated an average latency period of 0.6 years (95% CI: 0.4-0.8) between WGD and the expansion of the MRCA in patients under 40 years old, as compared to 1.7 years (95% CI: 1.1-2.4) in those over 40 (**Fig. 5A and Methods**).

These results, which are in agreement with the findings from PCAWG, represent a relatively shorter period between WGD and MRCA emergence compared to other cancer types(*33*). In tumours with LTA chromothripsis, we detected LOH at the terminal region of 17p, including the *TP53* locus, suggesting that LTA likely precedes WGD, as otherwise two independent LTA events would need to occur for the two terminal copies of 17p to be lost (**figs. S13 to S17**). Thus, in these tumours, WGD occurs during the short interval between the initial steps of LTA chromothripsis and the expansion of the MRCA.

To study the relationship between oncogene amplification and WGD, we next defined a scoring system based on rearrangement and copy number information to determine whether CGRs mediating oncogene amplification occur before or after WGD(*21*) (**fig. S30, Table S9 and Methods**). Overall, we found that in 98.8% (815/825) of CGRs, the majority of breakpoints per chromosome arm occurred after WGD. In 11% of these events, at least 5 breakpoints occurred prior to WGD, suggesting an initial pre-WGD event. At the rearrangement level, only 0.4% (7/1863) of foldbacks associated with an oncogene amplification were predicted to occur prior to WGD (**Fig. 4E-H**). These results are consistent with *in vitro* studies showing that WGD causes chromatin remodelling and genomic instability(*34*, *35*), and the higher rates of chromothripsis observed in WGD tumours(*3*). By contrast, SVs associated with the loss of tumour suppressors, such as *RB1*, consistently occurred before WGD. Together, these results indicate that WGD is a late clonal event that drives chromosomal instability and oncogene amplification.

**Fig. 4.**
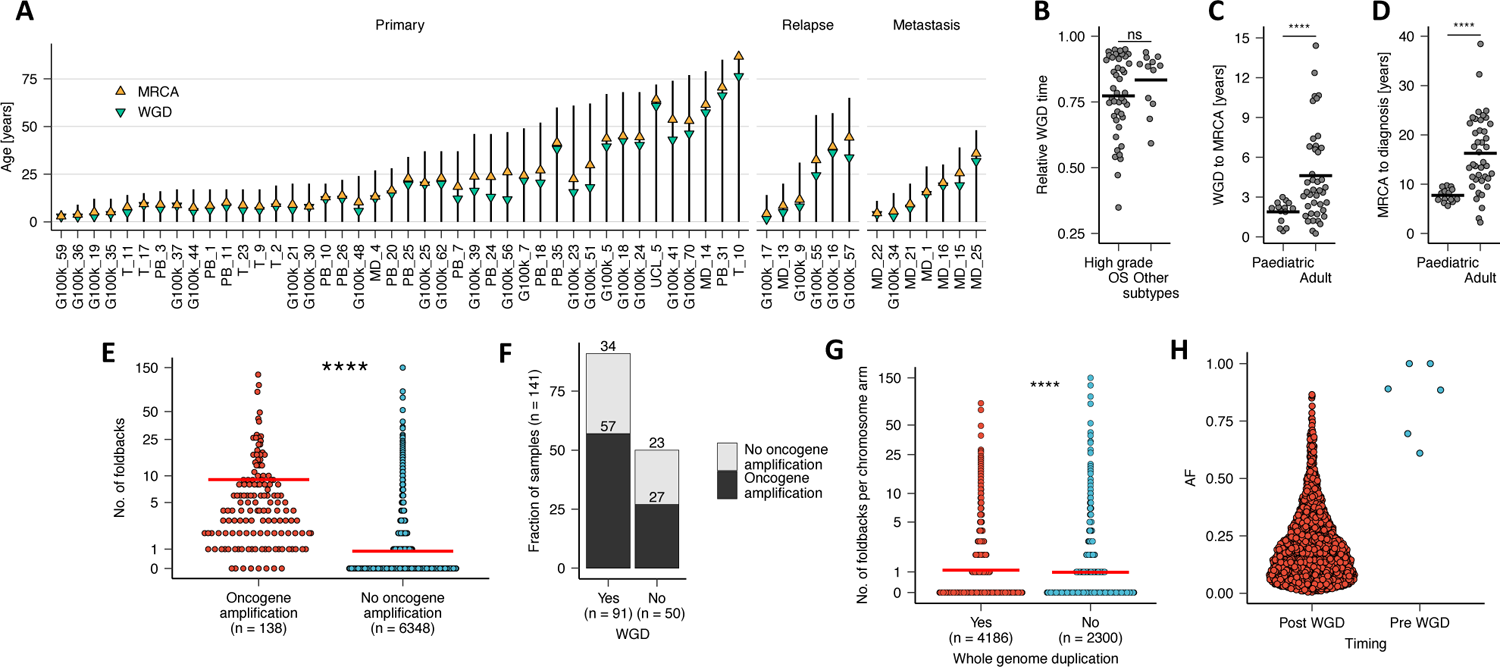
Timing of whole-genome doubling in high-grade osteosarcoma and relationship with oncogene amplification. **(A)** Timing estimates for the emergence of the most recent common ancestor (MRCA) and whole-genome doubling (WGD) during the evolution of paediatric and adult high-grade of osteosarcomas. **(B)** Relative timing estimates for WGD events in high-grade and other subtypes of osteosarcoma (parosteal, extraskeletal, periosteal and low-grade central). **(C)** Number of years between WGD and the emergence of the MRCA in paediatric and adult osteosarcomas. **(D)** Number of years between the emergence of the MRCA and diagnosis in paediatric and adult osteosarcomas. **(E)** Number of foldback inversions per chromosome arm stratified based on the presence of an oncogene amplification event in the same chromosome arm. **(F)** Fraction of HGOS tumours showing oncogene amplification stratified based on whole-genome doubling status. **(G)** Number of foldback inversions per chromosomes arm in HGOS tumours stratified based on whole-genome doubling status. **(H)** Allele fraction (AF) for foldback inversions stratified based on whether they were estimated to occur before or after WGD. Significance in **B**–**E** and **G** was assessed using the two-sided Wilcoxon’s rank test (\*\*\*\**P* < 0.00001); ns: not significant.

### Chromosomal losses precede whole-genome doubling and predict prognosis

Identification of robust biomarkers for accurate prognostication to inform clinical decision making remains an urgent clinical need for osteosarcoma(*16*). Previous studies based on SNP arrays identified an association between high genome-wide levels of LOH and prognosis(*36*). However, this and other biomarkers have not been validated in large cohorts for risk stratification(*16*). To evaluate the prognostic power of genomic aberrations, we conducted unbiased survival analysis for HGOS using diverse genomic and clinical covariates (**Methods**). Genome-wide LOH was the only genomic event significantly associated with progressive disease in both univariate and multivariate analysis correcting for known covariates affecting survival, such as metastasis at diagnosis (HR 54.2; 95% CI: 7.5-391; *P*=0.002; **Fig. 5A-B and Methods**).

**Fig. 5.**
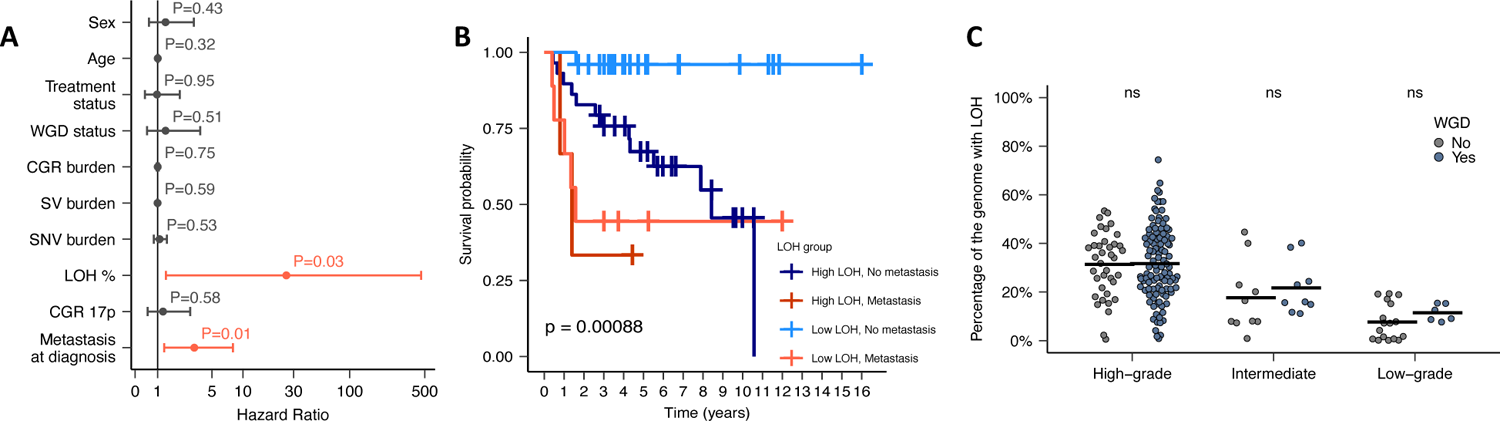
Genome-wide loss of heterozygosity (LOH) predicts overall survival for high-grade osteosarcoma. **(A)** Hazard ratios with 95% confidence intervals and *P* values computed using Cox proportional-hazards regression. **(B)** Kaplan–Meier plot showing the overall survival for high-grade osteosarcoma patients stratified according to the degree of genome-wide LOH and the presence of metastasis at diagnosis. **(C)** Percentage of the genome showing loss of heterozygosity (LOH) in high, intermediate and low-grade osteosarcomas from the G100k and TARGET cohorts stratified based on the presence of WGD. Significance in **C** was assessed using the two-sided Wilcoxon’s rank test; ns: not significant.

Chromosomal losses following *TP53* inactivation have been reported to precede WGD in mouse models of pancreatic ductal adenocarcinoma(*37*). Using the multi-region WGS data, we estimated that 60% (95% CI: 52.4 to 67.6%) of genomic regions with LOH are common to all regions of a tumour, suggesting that LOH is an early event that occurs before WGD. HGOS tumours had significantly higher levels of LOH than low-grade cases (average of 32% vs 14%, respectively; *P*<0.001, Wilcoxon’s rank-sum test). In contrast to other cancer types(*38*), we detected high levels of genome-wide LOH independent of WGD status (**Fig. 5C**). Consistent with the role of WGD in buffering the acquisition of biallelic mutations in essential genes as chromosomal losses accumulate(*38*), we found a depletion of inactivating mutations in essential genes in LOH regions suggestive of negative selection, although our analysis did not reach statistical significance, likely due to limited statistical power given the low mutational burden in osteosarcoma (**fig. S31**).

## Discussion

We report a comprehensive genomic analysis of the mechanisms underpinning cancer genome complexity and evolution in paediatric and adult osteosarcomas. We describe a novel rearrangement process, LTA chromothripsis, which provides a mechanistic basis for the generation of the most complex derivative chromosomes frequently observed in high-grade osteosarcoma of bone, which are characterised by chromothripsis patterns interleaved with segmental amplifications across multiple chromosomes(*19*). We find that such complex events are the result of a karyotypic evolution process in which the resolution of an initial dicentric chromosome generated by a single double-strand break in the *TP53* locus triggers chromothripsis, oncogene amplification in the form of ecDNA and segmental amplifications across multiple chromosomes. LTA chromothripsis is different from translocation-bridge amplification(*39*) and classical BFB cycles(*40*) in that LTA causes concomitant biallelic inactivation of *TP53*, leading to tolerance to WGD and genomic instability, and a multi-generational BFB cycle process that can engage additional chromosomes, thus leading to the amplification of oncogenes across multiple chromosomes. LTA chromothripsis also explains mechanistically the generation of segmental amplifications commonly detected in osteosarcomas(*19*). Through phylogenetic analysis, we show that LTA drives rapid karyotype evolution and clonal diversification, providing evidence for the role of chromothripsis in punctuated evolution and promoting rapid tumour growth. Somatic inactivation of one copy of *TP53* before LTA chromothripsis is required for malignant transformation in sporadic HGOS tumours. However, in Li-Fraumeni individuals, LTA is the only somatic event required to cause biallelic inactivation of *TP53* and oncogene amplification within a few cell divisions. Finally, we show that the degree of genome-wide LOH has strong prognostic power for HGOS, which may help improve the management of osteosarcoma patients, especially given the increasing availability of genome sequencing technologies in clinical settings.

## Data availability

WGS data from the participants enrolled in the 100,000 Genomes Project can be accessed via Genomics England Limited following the procedure described at: https://www.genomicsengland.co.uk/about-gecip/joining-research-community/. In brief, applicants from registered institutions can apply to join one of the Genomics England Research Network, and then register a project. Access to the Genomics England Research Environment is then granted after completing online training. Raw WGS data from the Gabriella Miller Kids First Pediatric Research Program: An Integrated Clinical and Genomic Analysis of Treatment Failure in Pediatric Osteosarcoma (project number 1 X01 HL 132378-01) are available at dbGAP under study accession code phs001714.v1.p1. The WGS data from the Therapeutically Applicable Research to Generate Effective Treatments (TARGET) program is available at dbGAP under study accession code phs000218. The raw WGS data from the Behjati et al. cohort(*19*) are available at the European Genome-Phenome Archive (EGA) under accession numbers EGAD00001000107, EGAS00001000196 and EGAD00001000147. The raw WGS data from the MDACC cohort(*21*) are available at the EGA under accession number EGAS00001003247. PCAWG analysis results, including the somatic copy number and rearrangement calls used in this study, are available at https://dcc.icgc.org/releases/PCAWG. The code used in this study is available at: https://github.com/cortes-ciriano-lab/osteosarcoma_evolution. The raw WGS data generated by The Cancer Genome Atlas (TCGA) can be accessed through controlled data access application via dbGAP under study accession code phs000178.

## Acknowledgements

J.E.V.-I., H.E., F.M. and I.C.-C. thank EMBL for funding. Funding was also provided by the Tom Prince Trust and the Sarcoma Foundation of America (SFA 20-05). Provision of patients’ samples from the RNOH was made possible through the Royal National Orthopaedic Hospital Pathology Department and the Research and Development Department, The Rosetrees Trust, Skeletal Cancer Trust, Sarcoma UK, The Bone Cancer Research Trust and The Pathological Society of Great Britain and Ireland, over the last two decades. SDN and TB are PhD students funded jointly by the Pathological Society of Great Britain and Ireland and the Jean Shanks Foundation. The project was also supported by the National Institute for Health Research, UCLH Biomedical Research Centre, and the UCL Experimental Cancer Centre. We thank the patients and their families for their participation in this study. Some figures were generated using BioRender.com. This research was made possible through access to data in the National Genomic Research Library, which is managed by Genomics England Limited (a wholly owned company of the Department of Health and Social Care). The National Genomic Research Library holds data provided by patients and collected by the NHS as part of their care and data collected as part of their participation in research. The National Genomic Research Library is funded by the National Institute for Health Research and NHS England. The Wellcome Trust, Cancer Research UK and the Medical Research Council have also funded research infrastructure. The authors thank Dr. Emma McCargow and Dr. Hélène Louis dit Picard for technical and administrative support.

## Author contributions

Conceptualization: AMF, ICC. Methodology: JEVI, SDN, KT, HE, MT, FM, AG, GE, AMF, ICC. Experimental work: KT, MT, TB, GE, AMF. Pathology review: SDN, FA, RT, TB, AMF. Investigation: JEVI, SDN, AMF, ICC. Visualisation: JEVI, SDN, AMF, ICC. Technical support: KT, MT, AG, AS, GE, AMF, ICC. Funding acquisition: AMF, ICC. Project administration: AMF, ICC. Supervision: AMF, ICC. Writing – original draft: JEVI, SDN, AMF, ICC. Writing – review & editing: JEVI, SDN, AMF, ICC.

## Methods

### Data sets

– **G100k RNOH cohort**. Paediatric and adult fresh frozen tumour samples were obtained from patients consented and enrolled in both the Genomics England 100,000 Genomes Project (G100k) as well as the Royal National Orthopaedic Hospital (RNOH) Biobank satellite of the UCL/UCLH Biobank for Health and Disease (REC reference 20/YH/0088). Tumours diagnosed as osteosarcoma by specialist pathologists in accordance with the WHO Tumour classification(*41*) were included in the study. Surplus tumour tissue from resection and or biopsy samples were collected and frozen as part of routine clinical practice. WGS data from matched blood samples were used as germline controls in genomic analyses. Prior to sequencing, tumour samples were reviewed, and eligible samples were selected based on a tumour content of >40% and <20% necrotic material. Multiple samples from individual tumours were included whenever suitable. DNA was extracted from matched tumour and blood samples using established protocols and in accordance with the 100,000 Genomes Project guidelines. DNA was sent for centralised library preparation and sequencing at the Illumina Laboratory Services (ILS) in Cambridge, UK(*42*). Sequencing was performed as part of the 100K Genomes Project. Samples were sequenced to an average depth of 117x (median 117x) and 43x (median 38x) for tumour and germline DNA, respectively. After processing of WGS data, 5 samples were discarded due to contamination, and 5 samples were discarded due to low purity values. The final GEL-100k cohort consisted of 215 tumour samples from 72 donors. Dual consented genomic data (somatic and germline) was shared to RNOH and EMBL-EBI. Data linked to local clinical data (no NHS Digital or NHS England data) were shared by Genomics England with RNOH and EMBL-EBI. No analysis was undertaken in the Genomics England Research Environment or National Genomic Research Library.
– **PCAWG and Behjati et al. cohorts.** We included 33 paediatric and adult osteosarcoma cases analysed by the TCGA/ICGC Pan-Cancer Analysis of Whole Genomes Project (PCAWG). In addition, we included 3 cases from the original study by Behjati et al. (*19*) not included in PCAWG. The average depth of sequencing for tumour samples was 44x (median 42x) and 36x (median 34x) for normal samples.
– **TARGET cohort.** We included 23 paediatric and adult osteosarcoma and matched germline WGS data sets from the TARGET osteosarcoma study (TARGET OS; dbGaP study phs000468.v21.p8). Patients were recruited through The Children’s Oncology Group (U.S.) and The Hospital for Sick Children (Toronto, Canada). TARGET OS is part of the NCI’s Therapeutically Applicable Research to Generate Effective Treatments (TARGET) program. The average depth of sequencing for tumour samples was 35x (median 44x) and 33x (median 47x) for normal samples.
– **Kids First cohort**. We included 39 osteosarcoma WGS data sets from 23 patients with matched germline WGS data from the Gabriella Miller Kids First Pediatric Research Program (dbGaP study phs001714.v1.p1). All samples were sequenced on the Illumina Hiseq X Ten platform. The average depth of sequencing for tumour samples was 49x (median 50x) and 25x (median 22x) for normal samples.
– **MDACC cohort.** We included WGS data from 30 cases from the MD Anderson Cancer Center (MDACC) generated by Wu et al.(*21*). The average depth of sequencing for tumour samples was 74x (median 75x) and 37x (median 37x) for normal samples.
– **UCL cohort.** To enrich our data set with uncommon subtypes of osteosarcoma, we performed whole-genome sequencing on the following cases from the RNOH Biobank, satellite of the UCL/UCLH Biobank for Health and Disease (REC reference 20/YH/0088): malignant giant cell tumour of bone (n=5), pagetic bone disease associated osteosarcoma (n=2), osteoclast rich HGOS (n=2), HGOS with unusual histological features (n=2), and *MDM2*-negative low grade central osteosarcoma (n=2). Between 350 and 500ng of tumour DNA were used to prepare DNA sequencing libraries using the TruSeq DNA PCR Free 350bp kit (Illumina). The samples were then sequenced on the Illumina NovaSeq6000 sequencing machine using S4 flow cells to generate 2 x 151 paired-end reads. Quality control for all steps was performed using a DNA Screentape on the Agilent TapeStation system. The average depth of sequencing for tumour samples was 77x (median 77x) and 34x (median 34x) for normal samples.

### Uniform processing of short-read whole-genome sequencing data

Raw sequencing reads from all cohorts included (356 samples from 197 patients) were mapped to the hg38 build of the human reference genome using BWA-MEM(*43*) version 0.7.17-r1188. Aligned reads in BAM format were processed following the Genome Analysis Toolkit (GATK, version 4.1.8.0) Best Practices workflow to remove duplicates and recalibrate base quality scores(*44*). Somatic and germline SNVs, multi-nucleotide variants (MNVs) and indels were called and filtered with SAGE (v2.8). Somatic SVs were called with GRIDSS2(*45*) (v2.12.0, https://github.com/PapenfussLab/gridss), annotated with RepeatMasker (v4.1.2-p1, http://www.repeatmasker.org) and kraken2(*46*)(v2.1.2), filtered with GRIPSS (v1.9) and clustered and visualised with LINX (v1.15)(*47*). B-allele frequency (BAF) of heterozygous SNP sites was computed using AMBER (v3.5) and read depth ratios were calculated with COBALT (v1.11). B-allele frequency information, read-depth ratios, SV breakpoints and allele frequencies of SNVs were integrated to estimate the purity, ploidy and copy number profile of each tumour using PURPLE (v2.54)(*47*). Quality control (QC) was performed using AMBER and PURPLE and samples were discarded when: (1) FAIL_CONTAMINATION: measured tumour contamination in homozygous sites from the normal sample was >10%; (2) FAIL_NO_TUMOR: no evidence of tumour was found in the sample – in these cases, samples were kept if at least one of the following criteria were met (i) the tumour had one or more HOTSPOT SV or point mutation, (ii) the number of somatic SNVs was > 1000, (iii) the number of somatic SVs was > 1000 (excluding single breakends), or (iv) the tumour sample had 3000 BAF points in germline diploid regions with a tumour ratio < 0.8 OR > 1.2 (i.e., there is evidence of some level of aneuploidy). SAGE, GRIPSS, AMBER, COBALT, PURPLE and LINX are developed by the Hartwig Medical Foundation (HMF) and code, tools and extensive documentation are freely available on github: https://github.com/hartwigmedical/hmftools. Missense point mutations predicted to be deleterious by MetaLR and MetaSVM, as implemented in Annovar(*48*) (version 2018Apr16), were considered pathogenic. Analysis of homologous recombination deficiency was performed using CHORD(*49*). To determine the Alternative Lengthening of Telomeres (ALT) status of tumours, we used the software TelFusDetector (v1; https://github.com/cortes-ciriano-lab/TelFusDetector) with default parameter values. In brief, TelFusDetector identifies ALT-associated telomere fusions in WGS data, which are then used to predict ALT status using Random Forest classification(*50*). We detected kataegis foci using the R package Katdetectr(*51*) (v1.2), requiring a minimum number of 10 mutations and a maximum intermutation distance of 1000 bps.

### Driver mutation analysis

Somatic mutations (point mutations, indels, SVs and copy number aberrations) mapping to cancer genes, as listed in the HMF driver panel(*52*) and reported as pathogenic by the PURPLE and LINX algorithms, were curated manually to assemble a list of high confidence mutations. In addition, hotspot mutations in *H3F3A,* a well-established driver of giant cell tumours of bone(*53*), were manually included, and in cases where single hits in highly recurrent osteosarcoma tumour suppressor genes (*TP53, RB1, PTEN, CDKN2A, ATRX, NF1*) were detected, the gene locus was visually inspected to verify the absence of second hits missed by our WGS data analysis pipeline due to segmental or arm level LOH.

SNVs and indels classified as pathogenic by PURPLE were annotated using ANNOVAR (*48*) (version 2018Apr16) and prioritised as follows:

– Tier 1: Clinically reportable and/or actionable driver mutations based on a list provided by the Cambridge Genomic Medicine Service (correspondence), and hotspot mutations with validated functional impact.
– Tier 2: All missense and nonsense mutations with a driver likelihood predicted by PURPLE > 0.95 (Tier 2a), and nonsense mutations with a predicted driver likelihood >0.5 and <=0.95 (Tier 2b). Missense mutations predicted as pathogenic by either MetaSVM or MetaLR, or by both VEST3 and CADDphred (Tier 2c).
– Tier 3: All remaining potential driver mutations reported by PURPLE with a driver likelihood less than 0.5 and reported as non-pathogenic by other pathogenicity scores. All non-hotspot mutations in *IDH1* and *IDH2,* as well as non-5’-UTR *TERT* mutations were automatically assigned to Tier 3 and excluded from further analysis.

The final set of driver mutations was generated as follows:

– All amplifications and partial amplifications of oncogenes listed in the COSMIC Cancer Gene Census or the HMF driver catalogue of at least 3 times the tumour ploidy.
– Biallelic mutations in tumour suppressor genes in the HMF driver catalogue panel(*52*).
– Heterozygous gain-of-function mutations in *TP53* with experimental evidence of promoting tumourigenesis in a heterozygous state(*54*).
– All Tier 1 and Tier 2a SNVs.
– Tier 2b and 2c SNVs in COSMIC Cancer Gene Census listed as driver genes or genes previously recognised as drivers in osteosarcoma(*19*).

### Driver-CGR associations

Driver alterations in tumour suppressor genes caused by structural variants (deletions, disruptions, or both) were determined to be associated with a CGR if: (1) a CGR-associated breakpoint mapped to the gene body resulting in the deletion or disruption of the gene, or (2) the genomic region encompassed by a CGR partially or completely overlapped the gene body. In the case of oncogenes, the breakpoints of the SVs causing oncogene amplification generally map outside of the gene body. Therefore, we considered oncogene amplifications to be CGR-associated if a CGR occurred in the same chromosome arm with an oncogene amplification. Oncogenes were deemed to be associated with foldback inversions if one or more foldback inversions mapped to the same chromosome arm harbouring the oncogene.

### Mutational signature analysis

The R package MutationalPatterns (v3.2.0)(*55*) was used for single base substitution (SBS) signature analysis. Samples with less than 500 SNVs (n=7) were excluded for this analysis. All mutational signature analysis (*de novo* extraction and refitting) was performed separately on samples from tumours which had been treated with chemotherapy (‘treated subgroup’) and those which had not been exposed to chemotherapy prior to sampling (‘untreated subgroup’). The 96-trinucleotide mutational spectra of somatic SNVs were computed using the *mut_matrix* function from the R package MutationalPatterns. For each subgroup, we performed hierarchical clustering of mutational spectra using 1 - cosine similarity as the distance metric(*56*). Samples assigned to clusters diverging above a distance of 2 were identified through manual inspection and were considered outlier samples. The remaining samples were assigned to the “common” cohort. The optimal number of signatures or rank (r) was determined by the *vb_factorize* function from the R package ccfindR. *De novo* signature extraction was performed on the mutational spectra of the common cohort using a rank of *r*, *r*+1 and *r*-1. The optimal rank values (r=8 and r=7 for the treated and untreated groups, respectively) were selected based on the goodness of fit of the extracted signatures to the observed mutational spectra and on the biological plausibility of the mutational signatures identified. Signatures identified *de novo* with a cosine similarity of >0.8 to a COSMIC v3 SBS signature were assigned the name of the matching COSMIC v3 SBS signature(*57*, *58*). Common signatures in human cancers (SBS1, SBS5, SBS8, SBS17b, SBS18, SBS37, and SBS40) were identified *de novo* in both treated and untreated cohorts, as well as a signature representing a combination of SBS2 and SBS13 previously described(*56*). Our analysis also detected a signature, which we termed “DN1”, with cosine similarity values to SBS39, SBS3 and SBS40 below our cut-off value for matching *de novo* and COSMIC signatures. Notably, signature DN1 did not decompose into more signatures by increasing rank. We detected SBS35 in the treated cohort, consistent with the relationship between this signature and chemotherapy treatment with platinum drugs. *De novo* signatures were then fitted to both common and outlier samples in each subgroup using the *strict refit* function, and samples with a cosine similarity between the original and the reconstructed spectra less than 0.95 were marked as outliers. Refits were accepted for the remaining samples. One signature with a cosine similarity of 0.8 to SBS37 was highly enriched in the Kids First cohort compared to the remaining samples. Given the demonstrated cohort bias and absence of this signature in published signature analyses of osteosarcoma(*24*, *34*, *35*), we considered this signature to represent a technical rather than a biological difference in this cohort. Signature refitting was then repeated without this signature, resulting in a minimal decrease in fitting quality. Therefore, we excluded this signature from further analysis. For each group (treated and untreated samples), we performed a second *de novo* signature extraction on outlier samples. Outlier signatures that could not be merged with any common signature (cosine similarity <0.75) were refit alongside the common signatures only in the samples in which they were detected if they led to an improvement in the goodness of fit of the reconstructed mutational spectra. Final outlier signatures were SBS31 in the treated cohort (detected in G100k_47), and SBS7a (detected in extraskeletal tumour G100k_16) and SBS30 (PB_21, case with a pathogenic germline variant *NTLH1*) in the untreated cohort. For the cases with multi-region WGS data, we assigned SNVs to three groups based on their clonality: clonal (detected in all tumour samples), subclonal (detected in a subset of tumour samples), and unique (present in just one tumour sample). We then performed signature refitting on each set of SNVs based on the treatment status of each sample.

### Analysis of positive selection

Selection analysis was performed with the R package dNdScv(*59*) using the trinucleotide substitution model with 192 rates and covariates for the reference genome hg38. No limits were set for the maximum number of coding mutations per sample or gene. dNdS values were obtained for missense, nonsense, and splicing mutations. The LOH status of each mutation was set when the minor allele copy number < 0.5 and the total copy number ≥ 0.5. For samples with WGD, mutations were classified as pre-WGD if the mutation copy number was ≥1.75 and post-WGD if the mutation copy number was ≤1.25. All other mutations were classified as “unknown”. Cancer genes for the selection analysis were obtained from the COSMIC Cancer Gene Census. Essential genes were selected as previously described(*60*).

### Analysis of loss of heterozygosity

Regions of the genome with LOH were identified as those with a minor allele copy number <0.5 and a major allele copy number ≥0.5. Genome-wide LOH was determined as the total size of all genomic segments with LOH divided by the genome size. Arm level LOH events were defined as chromosome arms with >90% of the arm length in a state of LOH. The p arm of acrocentric chromosomes (13, 14, 15, 21, 22) and chromosome Y were excluded from this analysis. In the case of tumours with multi-region WGS data, we classified LOH events into clonal and subclonal depending on whether LOH was observed in all or a subset of the tumour regions, respectively.

### Detection of complex genomic rearrangements

Complex genomic rearrangements (CGR) were detected using ShatterSeek (version 0.7). ShatterSeek calls were filtered using recommended cut-off values(*3*) and the *P* values for all statistical tests were corrected using the False Discovery Rate (FDR) method. In brief, we made a CGR call if the following criteria were satisfied: (i) at least 6 interleaved intrachromosomal SVs, (ii) the fragment joins test was not significant (FDR-corrected *P* > 0.05), and (iii) either the chromosomal enrichment or the exponential distribution of breakpoints test were significant (FDR-corrected *P* < 0.01). Additionally, we classified CGRs as “chromothripsis” when the copy number for at least 7 contiguous genomic segments oscillates between 2 total copy number states. Chromothripsis events were further classified as “canonical chromothripsis”, when at least 60% of the adjacent copy number segments showed copy number oscillations, and as “chromothripsis with other complex events” otherwise. Foldback inversions were detected from the SV-clustering output of LINX(*47*). Inversions smaller than 50kb were also classified as foldback(*24*). To detect ecDNA events, we used the AmpliconSuite pipeline (v0.1032.3)(*61*), a wrapper for AmpliconArchitect (v1.3) and AmpliconClassifier (v0.4.7). Visualization of CGRs was performed using ReConPlot(*62*).

To determine whether individual CGRs were clonal or subclonal, we reasoned that those CGRs occurring before the clonal expansion of the most recent common ancestor (MRCA) should be detected in all tumour regions analysed, whereas subclonal CGRs occurring after the expansion of the MRCA should be detected in just a subset. Therefore, we first detected CGRs in each chromosome and tumour region separately. Next, we intersected their coordinates, defined as the region encompassed between the most downstream and most upstream breakpoints involved in each CGR. We considered CGRs detected in the same chromosome across two or more different tumour regions to be the same event if the regions encompassed by each CGR overlapped by at least 25%, in which case the coordinates of each CGR were updated. Specifically, the CGR span in each chromosome and tumour region was defined as the genomic coordinate of the most upstream and downstream breakpoint coordinates across the tumour regions showing the CGR.

### Timing of WGD and the expansion of the MRCA

Timing analysis of WGD events was performed as previously described(*63*). Briefly, the multiplicity (i.e., number of chromosomal copies harbouring a given mutation), cancer cell fraction and copy number of clock-like SNVs (C>T mutations in NpCpG contexts) were used to determine subclonal versus clonal mutations using 1,000 bootstrap resamples. Major allele copy number information was then used to stratify clonal mutations based on their occurrence before or after WGD (pre-WGD and post-WGD, respectively). Tumours with at least 50 clock-like SNVs, tumour purity >0.3 and with patient age data available were selected for WGD timing analysis. We estimated the relative timing of WGD as the average fraction of pre-WGD clonal clock-like mutations in non-amplified regions (i.e., genomic regions with major and minor allele copy number values of 2 and 2, and 2 and 0, respectively). The relative timing for the MRCA was determined as the average proportion of clonal clock-like mutations across resamples.

To estimate the absolute timing of WGD and the expansion of the MRCA, we used the age at diagnosis to compute the mutational rate per sample as the number of clonal clock-like mutations per year adjusted for the effective genome size. Pan-cancer analysis of matched primary and relapse samples demonstrated that there is an acceleration in the mutation rate during the last stages of clonal evolution(*33*). To account for mutation acceleration, we applied an acceleration rate of 5x to the base mutational rate as previously reported(*33*). Absolute timing of WGD (using years as unit of time) was calculated using the number of clock-like mutations estimated to occur before WGD. Specifically, we multiplied the total number of clonal clock-like mutations by the estimated relative timing for the WGD events, and adjusted the mutation rate using an acceleration rate of 5x(*33*).

### Timing of SVs relative to WGD

To determine the timing of SVs relative to WGD, we used the copy number values on each side of the breakpoints, the jump in copy number between the genomic segments connected by each rearrangement, as well as the allelic fraction of rearrangement breakpoints(*8*). Specifically, we defined copy number jump (CN jump) as the change in copy number between the two genomic regions bridged by each SV, and copy number junction (CN junction) as the number of DNA copies (also known as multiplicity) in which each breakpoint was detected. In addition, we computed the purity-adjusted allele frequency (AF) for each breakpoint. We defined a timing score based on the following principles.

● We consider that a given breakpoint only occurs once in each tumour. That is, that the probability for the same exact SV to occur more than once in a tumour is zero.
● The reordering and reassembly of DNA fragments in a chromothripsis event does not cause copy number amplification.
● Genomic regions with a minor allele copy number of 0 and a total copy number of 2 or more are considered to be the result of a loss followed by whole-genome doubling, as we consider the probability of two independent losses in the two copies of the same parental chromosome after WGD to be negligible(*38*).

For this analysis, we focused on 158 regions from 92 tumours showing WGD. Due to the lack of reliable copy number jump and junction information, we did not perform timing analysis for breakpoints in haploid or diploid regions (total copy number of 1, or major and minor allele copy number values of 1), and breakpoints mapping to copy number segments less than 100bp in length.

AF, CN jump and CN junction information for each breakpoint were scored as follows:

● AF score (AF_score). We computed AF values for all breakpoints in each chromosome and assigned AF scores as follows: AF_score=1: breakpoints with an AF value above the 50th percentile; AF_score=2: breakpoints with an AF value between the 25th and the 50th percentiles; and AF_score=3: breakpoints with an AF value below the 25th percentile.
● Copy number jump scores (CN_jump_score). We assigned a CN_jump_score of 1 to breakpoints with CN jump values of 0, and a CN_jump_score of 0 otherwise.
● Copy number junction scores (CN_junction_score). CN_junction_scores were assigned on the basis that breakpoints occurring prior to WGD would be present in all copies of the affected allele. Thus, we assigned a CN_junction_score of 1 to breakpoints with a multiplicity equal to the major allele copy number (i.e., the number of copies of the most amplified allele). We assigned a CN_junction_score of 2 to breakpoints with a multiplicity at least two thirds the major allele copy number, a CN_junction_score of 3 to breakpoints with a multiplicity of at least one third but less than two thirds of the major allele copy number, and a CN_junction_score of 4 to breakpoints with a multiplicity less than a third of the major allele copy number.

The final timing score was calculated as the sum of the AF, copy number jump, and copy number junction scores. Thus, the timing score has a minimum and a maximum value of 3 and 9, respectively. Manual inspection of canonical chromothripsis events with clear evidence of occurring before or after WGD was used to set thresholds for likelihood of breakpoints occurring before and after WGD (see examples in **fig. S30**). Breakpoints with a total score ≤4 were classified as pre-WGD, and as post-WGD otherwise.

Next, we assessed the timing of CGR-associated breakpoints on a per chromosome arm basis. First, we considered a chromosome arm to be rearranged by a CGR before WGD when at least 50% of the breakpoints mapping to the CGR had a timing score of 3 or 4. Because chromosome arms are sometimes rearranged by multiple mechanisms throughout tumour evolution (e.g., LTA followed by WGD and chromothripsis), we considered chromosome arms to have a pre-WGD component when 5 or more CGR-associated breakpoints had timing scores ≤4.

### Phylogenetic analysis using short-read whole-genome sequencing data

We constructed a binary matrix for each tumour with rows indexed by somatic SNVs or SVs, and columns by tumour regions such that the *i,j* entry in each matrix was set to one if mutation *i* was present in region *j*, and to zero otherwise. To reconstruct phylogenetic trees, we used the UPGMA (unweighted pair group method with arithmetic mean) method (*treeUPGMA*) as implemented in the R package phangorn(*64*). Branch lengths were computed by counting the number of mutations (SNVs or SVs) shared by only the descendants of a given branch. Visualisation of phylogenetic trees was performed using custom scripts, which are available at https://github.com/cortes-ciriano-lab/osteosarcoma_evolution. A complete analysis of tumour phylogenetics and intra-tumour heterogeneity for tumours with multi-region WGS data and for individuals with WGS data from multiple tumours can be found in figs. S32 to S65.

### Survival analysis

We performed survival analysis using the Cox proportional hazards model as implemented in the R package survminer (version 0.4.9)(*65*). Significance was assessed by the likelihood ratio test using a cut-off value for statistical significance of 0.05. The proportional hazards assumption was tested using the *cox.zph* function from the R package survival(*66*). The LOH threshold used to stratify high-grade osteosarcomas into high LOH and low LOH groups was determined through a ROC curve analysis of survival status and percentage of genomewide LOH using the R package pROC(*67*).

### Estimation of tumour DNA ploidy

We performed DNA ploidy analysis by flow cytometry for a random subset of osteosarcoma cases. The most representative formalin-fixed, paraffin-embedded (FFPE) block was selected for each case. For each case, 50 μm scrolls were cut and deparaffinized by xylene, then rehydrated with ethanol and water. Samples were then dissociated into single nuclear suspensions using Proteinase from *Bacillus licheniformis* (Sigma-Aldrich P5380). Nuclear suspensions were filtered through 40 μm filters and resuspended in FACS buffer with 2 μM EDTA. Nuclear density (nuclei/mL) was estimated for each sample by haemocytometry, and samples were standardised to approximately 2,000,000 nuclei/mL. Samples were stained with DAPI (Thermo Scientific 62247) at a concentration of 1:1000 and incubated at room temperature in the dark for 1h. Stained samples were run through the Fortessa A flow cytometer (BD Fortessa X20), using a wavelength of V450/50. Events were gated using forward scatter and side scatter to exclude doublets and debris. Prior to the tumour samples being processed, an external control sample (normal adult kidney) was run to obtain the approximate mean DAPI fluorescence intensity of a diploid population. To calculate tumour ploidy, the mean DAPI fluorescence intensity (MFI) of the background diploid cell population was taken as the MFI representing the ploidy of a diploid cell (MFI-2n). The MFI of the largest, discrete tumour subclone was taken as the tumour MFI(MFI-t). Ploidy was then calculated as 2(MFI-t/MFI-2n).

### DNA extraction and processing for nanopore sequencing

Tumour genomic DNA was extracted from fresh-frozen tissue with the Nanobind tissue kit (Circulomics NB-900-701-01, now PacBio SKU 102-302-100) and Germline DNA from blood with the Nanobind CBB kit (Circulomics NB-900-001-01, now PacBio SKU 102-301-900). After extraction, the genomic DNA (gDNA) was homogenised by 3-10 passes of needle shearing (26G) and 1h of incubation at 50°C. All samples were quantified on a Qubit fluorometer (Invitrogen, Q33226) with the Qubit BR dsDNA assay (Thermo Fisher Scientific, Q32853) and manual volume checks were performed. The DNA size distributions were assessed at each relevant step by capillary pulse-field electrophoresis with the FemtoPulse system (Agilent, M5330AA and FP-1002-0275). 4-10 μg of gDNA were either fragmented to a size of 10-50kb with the Megaruptor 3 (Diagenode, E07010001 and E07010003) or fragmented to a size of 15-25kb with a gTUBE (Covaris, 520079) centrifuged at 1,500rcf. All samples were depleted of short DNA fragments (less than 10kb long) with the short read eliminator (SRE) kit (Circulomics, SS-100-101-01; now PacBio 102-208-300) or the SRE XS kit (Circulomics SS-100-121-01; now PacBio SKU 102-208-200) when sample availability or concentration was limiting.

### Library preparation and nanopore whole-genome sequencing

Sequencing libraries were generated from 600ng up to 1.8μg of gDNA with 1 library prepared for germline samples and 2 libraries prepared for tumour samples with the SQK-LSK110 or SQK-LSK114 kits (Oxford Nanopore Technologies, ONT), respectively, according to manufacturer’s recommendation with minor modifications. Briefly, the samples were end-repaired by adding 2 μl NEBNext FFPE DNA Repair Mix (NEB, M6630) and 3 μl NEBNext Ultra II End Prep Enzyme Mix (NEB, E7546), incubated for 10 min at room temperature followed by 10 min at 65 °C, then cleaned up with 1 × AMPure XP beads (Beckman Coulter, A63880) and eluted in 60 μl of EB. The end-repaired DNA was ligated with 5 μl Adapter Mix (ONT, SQK-LSK110) using 8 μl NEBNext Quick T4 DNA ligase (NEB, E6056) at 21°C for up to 1h. The adapter-ligated DNA was cleaned up by adding a 0.4 × volume of AMPure XP beads. Sequencing libraries were quantified and using the average peak size of the samples determined after sample preparation on the Femto Pulse. 20fmol of the obtained sequencing libraries were loaded onto R9.4 flow cell (ONT), or 10fmol of libraries were loaded onto R10 flow cells and sequenced on a PromethION48 (ONT). The libraries were stored overnight in the fridge. After 24h, sequencing runs were paused and a DNAse treatment or nuclease flush (ONT, WSH-003) was performed and 20fmol of the libraries were re-loaded on the flow cells.

Full details of the protocol can be found at https://research-help.genomicsengland.co.uk/display/GERE/Genomic+Data+from+ONT?preview=/38046759/38047942/v1_protocol_ONT_LSK109.pdf.

### Long-read nanopore sequencing data analysis

We aligned long-read nanopore WGS data to the hg38 build of the human reference genome using minimap2 (version 2.24)(*68*) with parameters “-ax map-ont--MD”. Copy number variation was estimated using Mosdepth (version 0.3.2)(*69*) with parameters “-n--fast-mode--by 1000” to obtain a coverage profile, followed by Spectre (version 0.1) (https://github.com/fritzsedlazeck/Spectre) with a minimum copy number length of 10,000bp to detect regions of deletion and duplication. We detected structural variants with SAVANA (version 1.0.4)(*70*) using default parameter values.

## Supplementary Tables

**Table S1.** Description of the tumour samples, main genomic aberrations detected in each tumour and associated clinical data.

**Table S2.** Summary of demographic and clinical data for the cases included in this study.

**Table S3.** Description of mutations in common cancer driver genes.

**Table S4.** Contribution of SBS mutational signatures to the mutational patterns detected in each tumour sample.

**Table S5.** Coordinates, characteristics, and classification of the CGRs detected in tumours across all cohorts analysed.

**Table S6.** Coordinates, characteristics, classification and clonality of the CGRs detected in tumours with multi-region and multi-sample WGS data.

**Table S7.** Comparison of tumour ploidy results by flow cytometry and WGS estimated with PURPLE.

**Table S8.** Analysis of the timing of WGD and the expansion of the MRCA.

**Table S9.** Analysis of the timing of SVs and CGRs per chromosome arm relative to WGD.

---

**Fig. S1.**
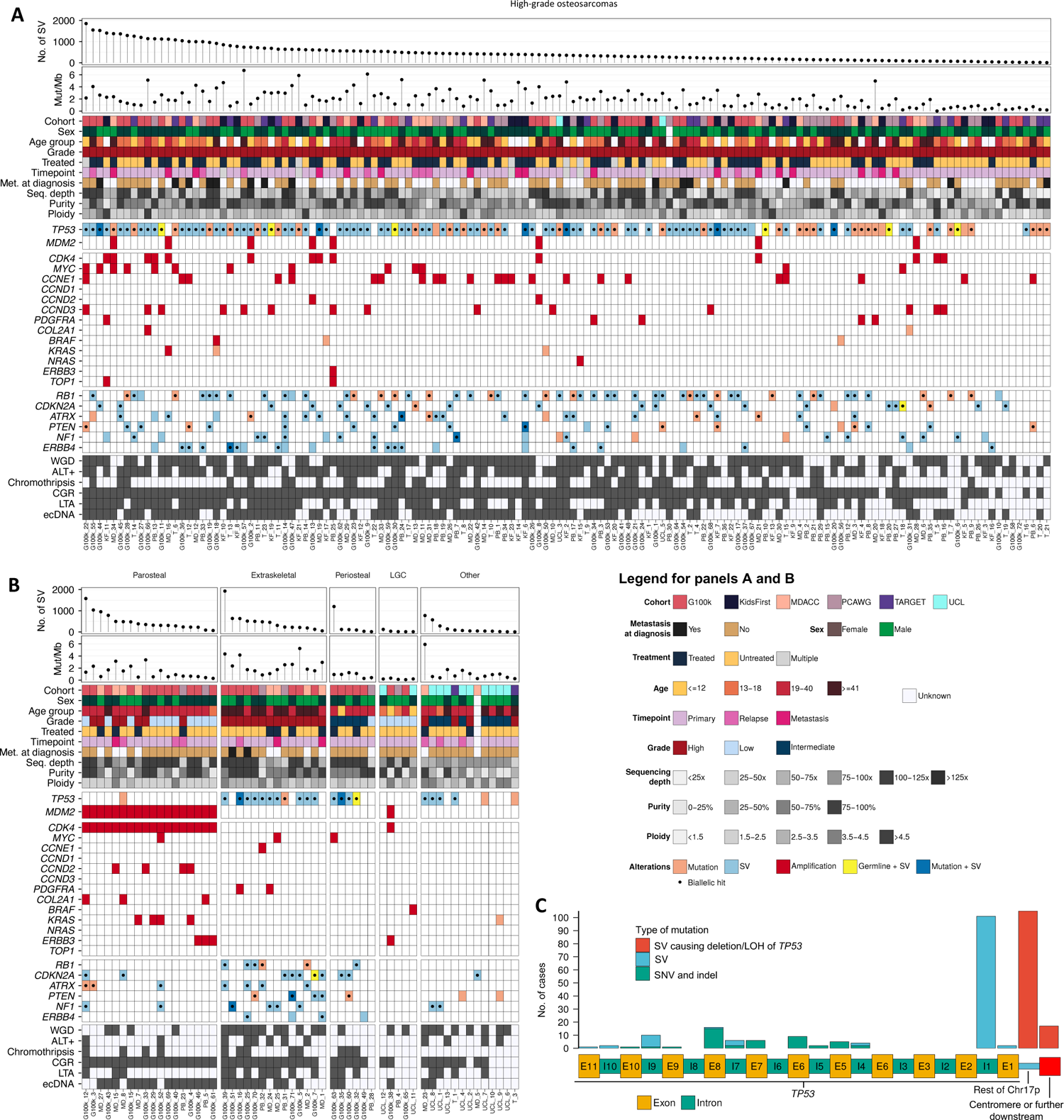
Genomic and clinical landscape of the osteosarcoma samples analysed in this study. Clinical information, histopathologic features, mutations and amplifications in driver genes, presence of WGD, chromothripsis, CGRs, and ecDNA in high-grade (**A**) and other subtypes (**B**) of osteosarcomas. (**C**) Distribution of somatic mutations causing the inactivation of *TP53*. ALT+: positive for Alternative Lengthening of Telomeres(ALT) pathway. LTA: loss-translocation-amplification. For cases with multi-region WGS data available, only the tumour sample with the highest tumour purity is shown. Patient cohorts: G100k (Genomics England 100,000 Genomes Project; new data generated for this study), KF (Kids First), MD (MD Anderson Cancer Center, Wu et al., *Nature Communications* 2020), PB (PCAWG cases from Behjati et al. *Nature Communications*, 2017), T (NCI’s Therapeutically Applicable Research to Generate Effective Treatments (TARGET) program, UCL (University College London; new data generated for this study).

**Fig. S2.**
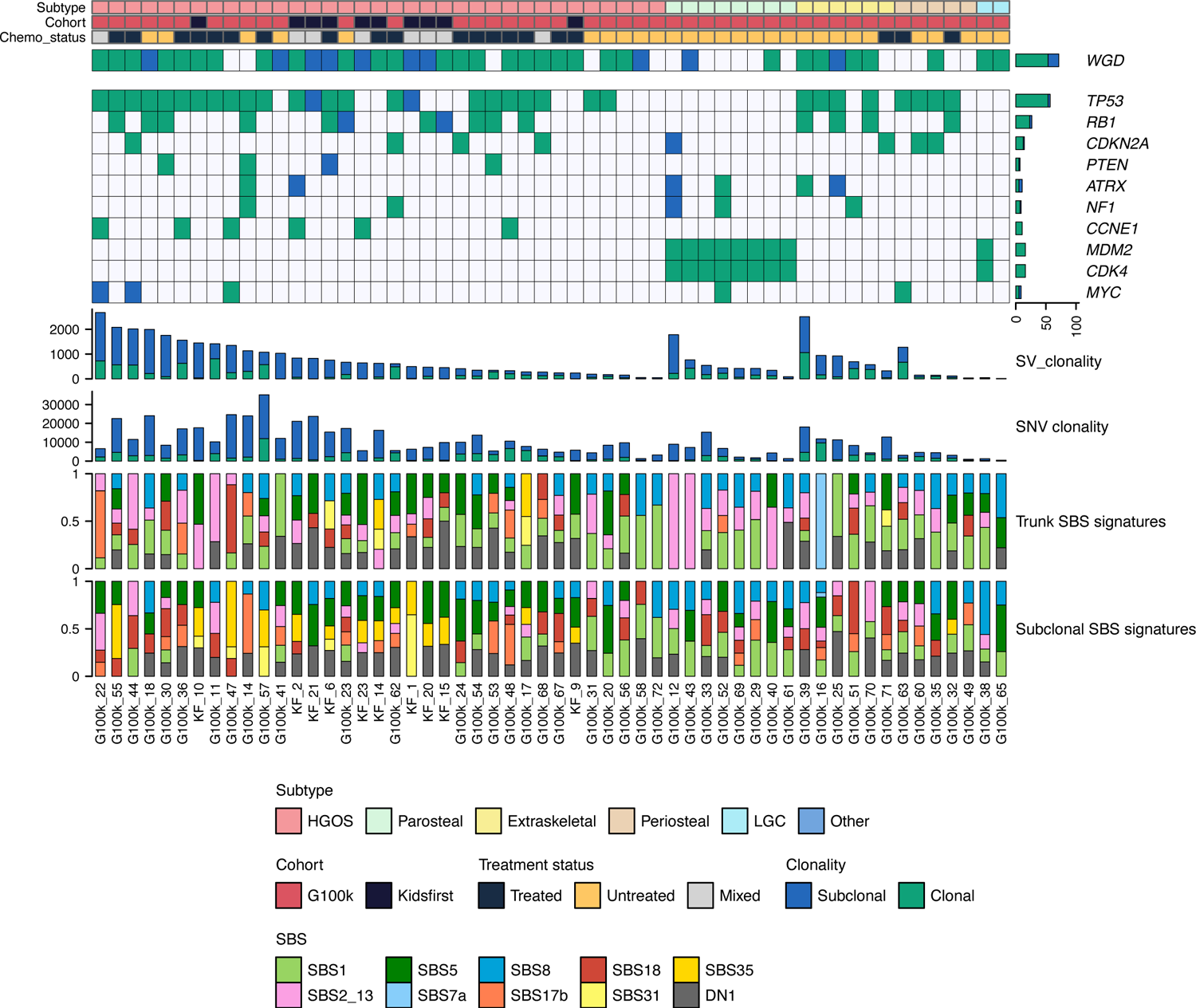
Intra-tumour heterogeneity of genomic alterations and single-base substitution (SBS) mutational signatures estimated using multi-region WGS data. Clonal and subclonal alterations are shown in green and blue, respectively. Patient cohorts: G100k (Genomics England 100,000 Genomes Project; new data generated for this study), KF (Kids First).

**Fig. S3.**
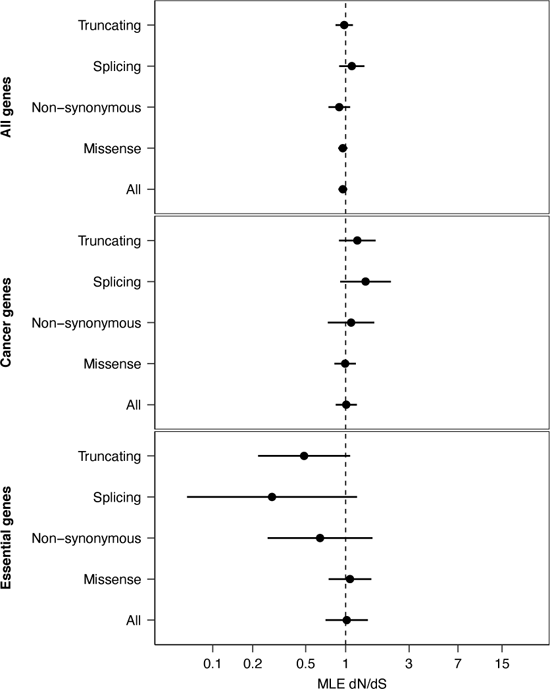
Analysis of selection for subclonal coding mutations. dN/dS values for subclonal coding mutations in high-grade osteosarcomas with multi-region data available calculated for all genes, essential genes, and cancer-associated genes. dN/dS values equal to, greater and lower than 1 are consistent with neutral evolution, positive selection, and negative selection, respectively. Point dN/dS estimates with 95% confidence intervals are shown. MLE dN/dS: maximum-likelihood dN/dS estimates.

**Fig. S4.**
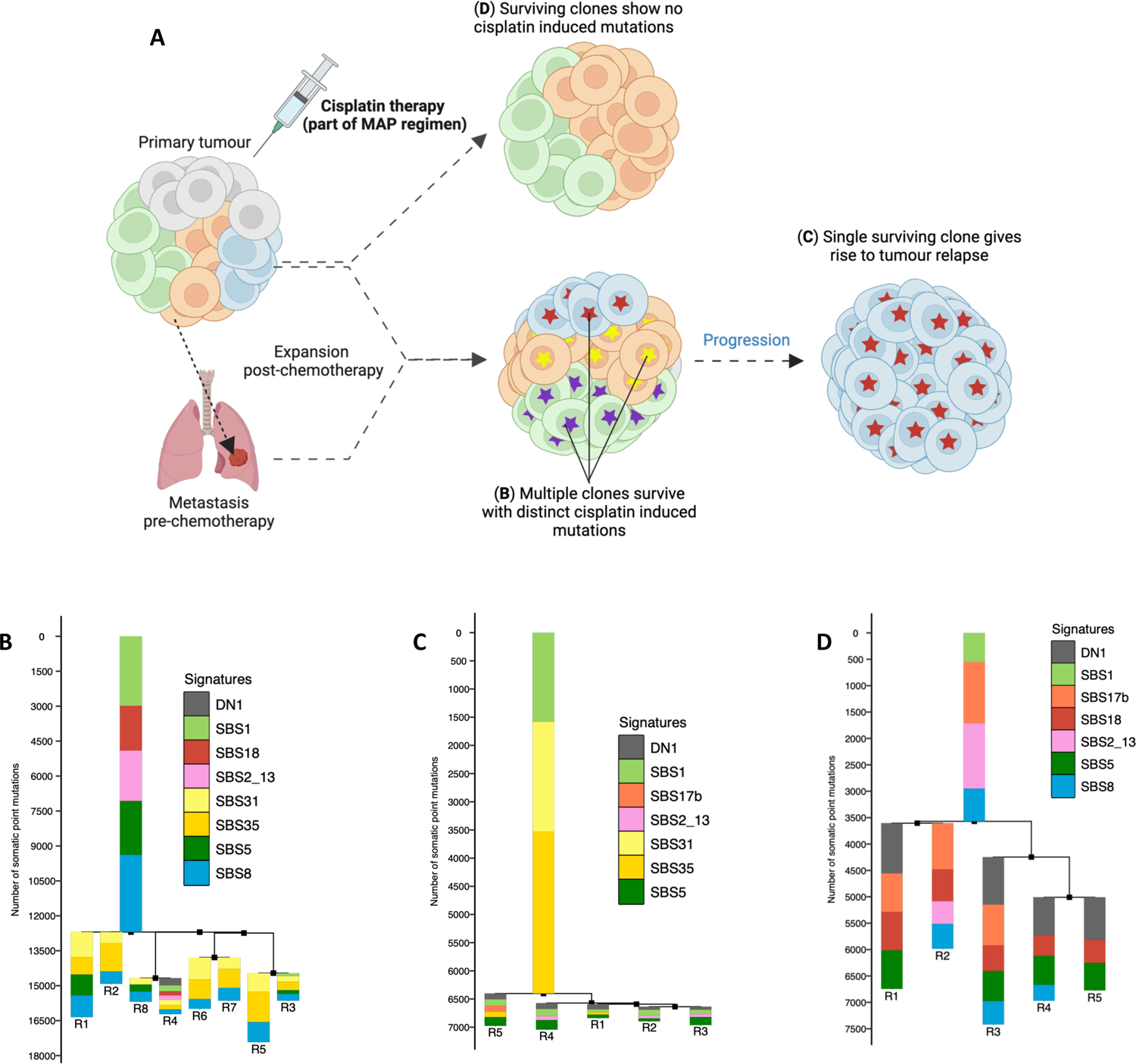
Impact of chemotherapy treatment on the clonal dynamics in high-grade osteosarcoma. **(A)** Schematic representation of the three types of clonal dynamics observed in high-grade osteosarcomas in response to chemotherapy treatment. **(B)** Phylogenetic tree established using somatic single-nucleotide variants detected in multi-region WGS data from case G100k_57. It can be observed that multiple clones receive (as indicated by the accumulation of chemotherapy-induced mutations; SBS31 and SBS35) and survive chemotherapy treatment. **(C)** Phylogenetic tree established using somatic single-nucleotide variants detected in multi-region WGS data from case G100k_36. In this case, a clonal sweep occurs in which a single clone received and survives chemotherapy treatment, and subsequently expanded**. (D)** Phylogenetic tree established using somatic single-nucleotide variants detected in multi-region WGS data from case G100k_17. Finally, in some osteosarcomas, multiple clones survive treatment and show no evidence of accumulation of mutations induced by chemotherapy treatment, as indicated by the absence of mutations attributed to single-base substitution mutational signatures 31 and 35.

**Fig. S5.**
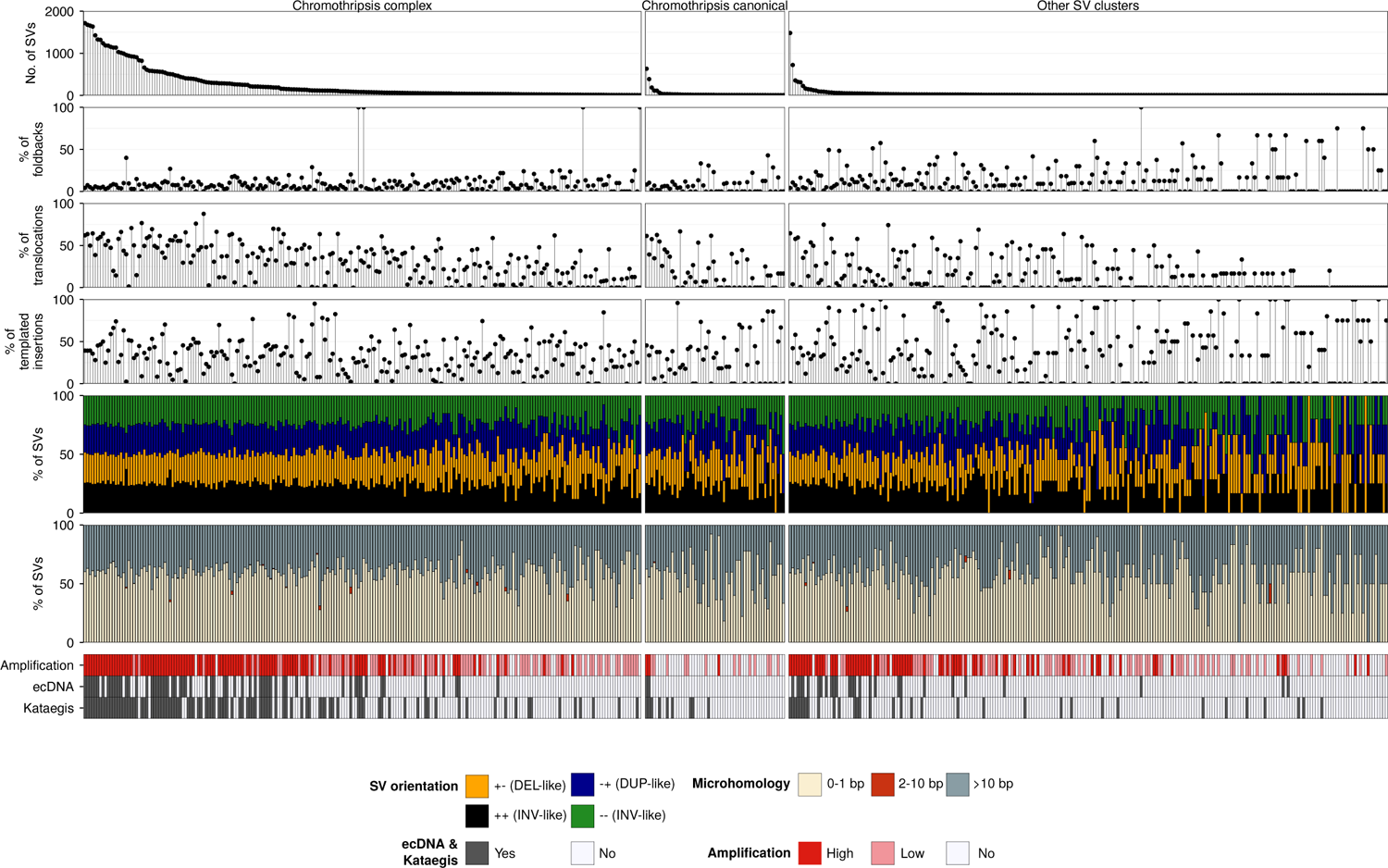
Analysis of rearrangements mapping to complex clusters of SVs. Each column summarizes the properties of the rearrangements mapping to complex SV clusters. DEL-like, deletion-like rearrangement; DUP-like, duplication-like rearrangement; h2hINV-like, head-to-head inversion; t2tINV-like, tail-to-tail inversion.

**Fig. S6.**
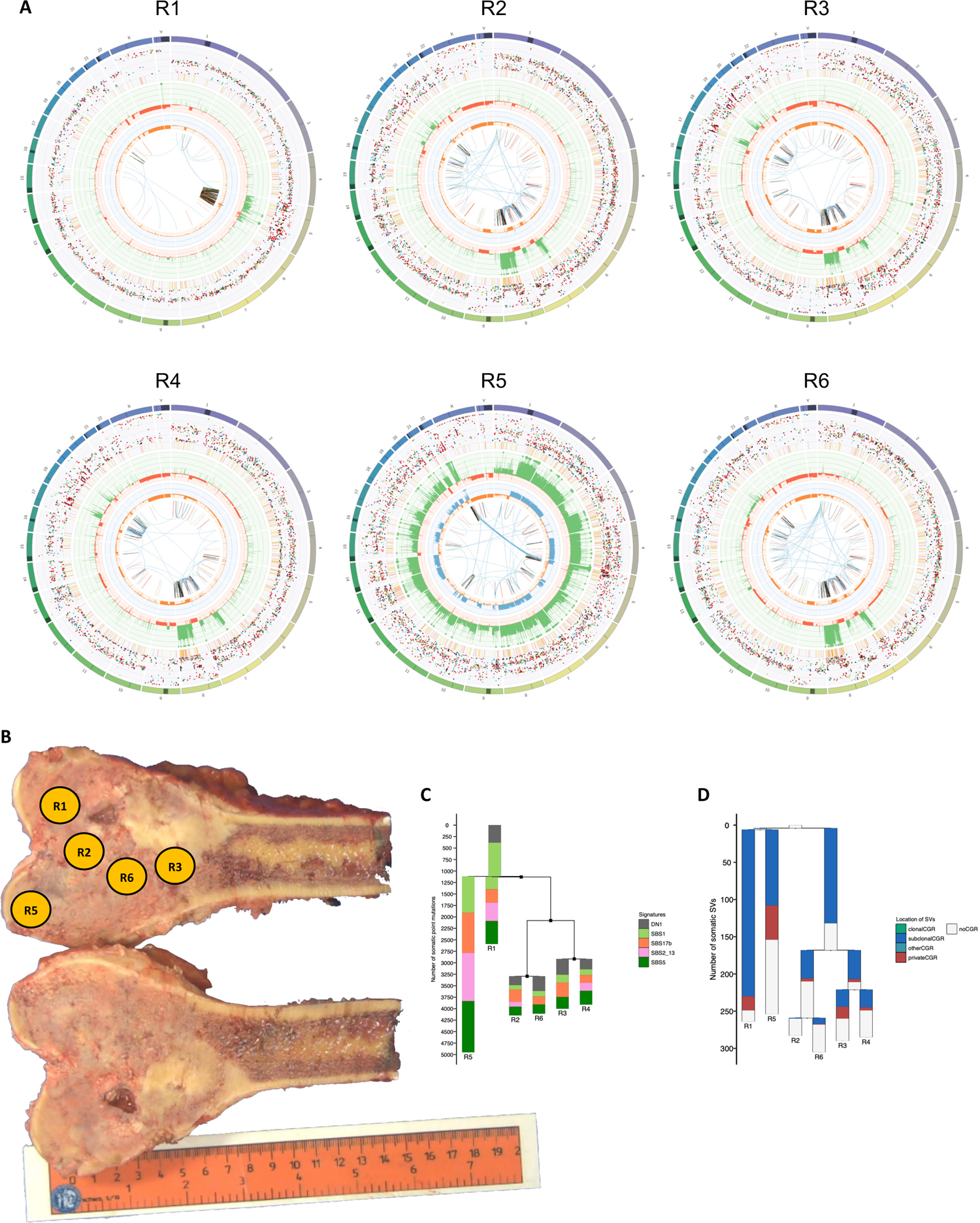
Chromothripsis drives clonal diversification and karyotype heterogeneity in osteosarcoma. **(A)** Circos plots representing the landscape of somatic aberrations detected in WGS data from six tumour regions (R) from case G100k_41. **(B)** Macroscopic image of the primary tumour from case G100k_41. Sampling regions are labelled. Region 4 corresponds to a biopsy sample, whose location within the primary tumour was not recorded. The side of the ruler is marked in centimeters. Phylogenetic trees constructed using somatic SNVs (**C**) or SVs (**D**) detected in multi-region WGS data from case G100k_30. The colours of the bars in the SNV tree represent the contribution of different SBS mutational signatures to the SNVs detected. The colours in the SV tree indicate whether the SVs in the tree map to clonal, subclonal or private CGRs (green, blue, and red), respectively, or outside CGR regions (grey).

**Fig. S7.**
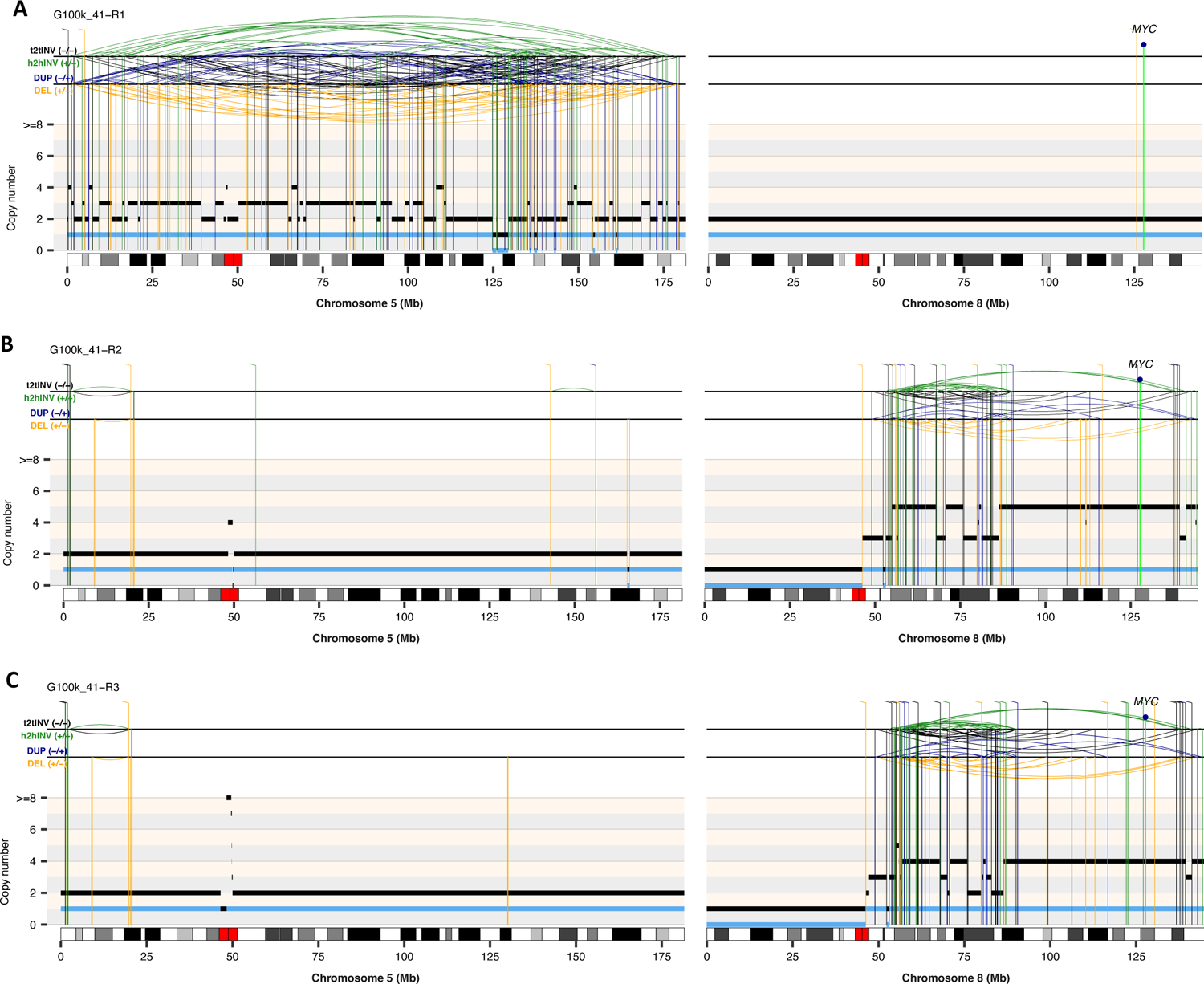
Rearrangement profiles of chromosomes 5 and 8 computed using somatic copy-number and SV data from regions 1. (**A**), 2 (**B**), and 3 (**C**) for case G100k_41. The total and minor copy-number data in **A-D** are represented in black and blue, respectively. DEL, deletion-like rearrangement; DUP, duplication-like rearrangement; h2hINV, head-to-head inversion; t2tINV, tail-to-tail inversion.

**Fig. S8.**
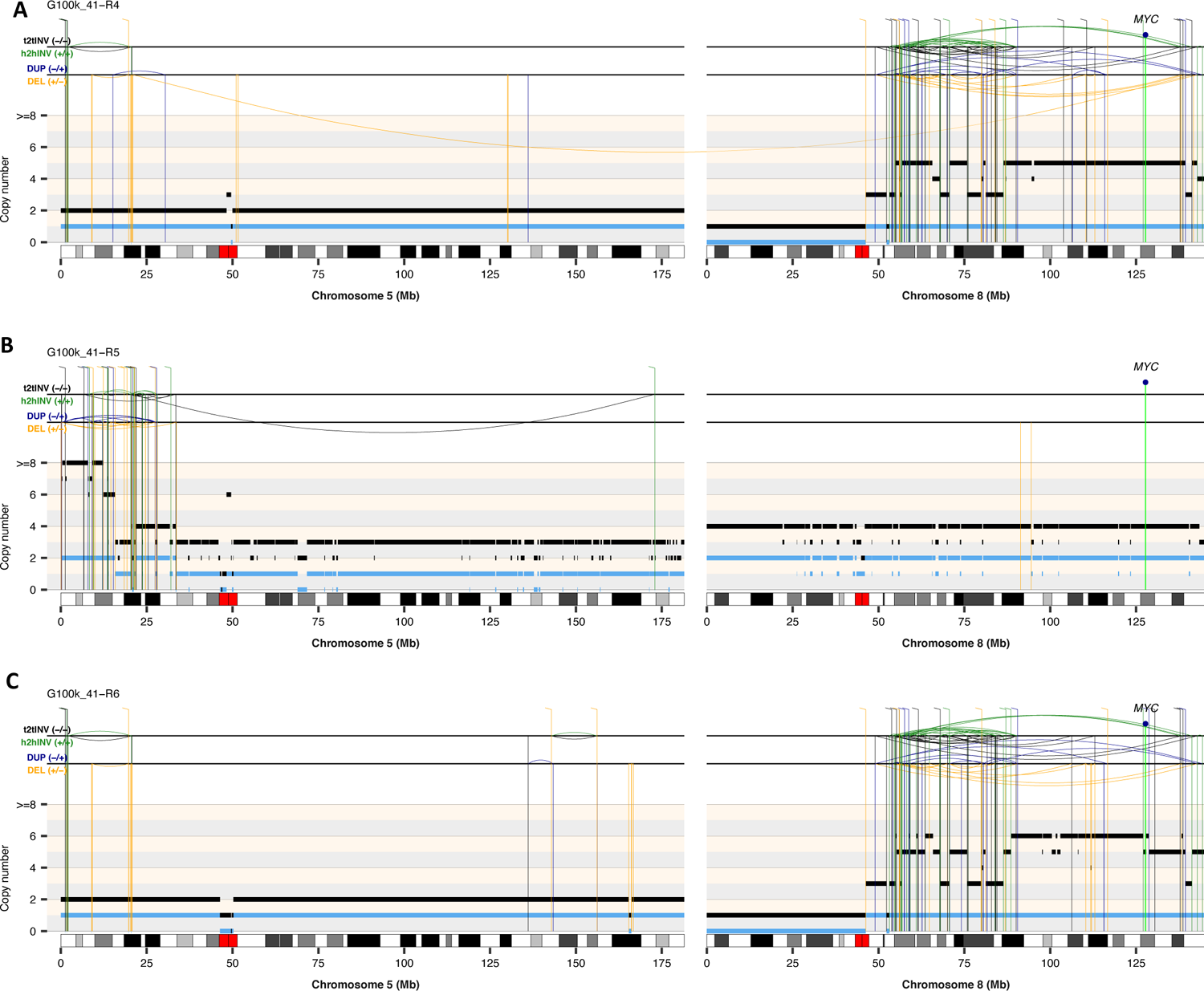
Rearrangement profiles of chromosomes 5 and 8 computed using somatic copy-number and SV data from regions 4 (**A**), 5 (**B**), and 6 (**C**) for case G100k_41. The total and minor allele copy-number data in **A-D** are represented in black and blue, respectively. DEL, deletion-like rearrangement; DUP, duplication-like rearrangement; h2hINV, head-to-head inversion; t2tINV, tail-to-tail inversion.

**Fig. S9.**
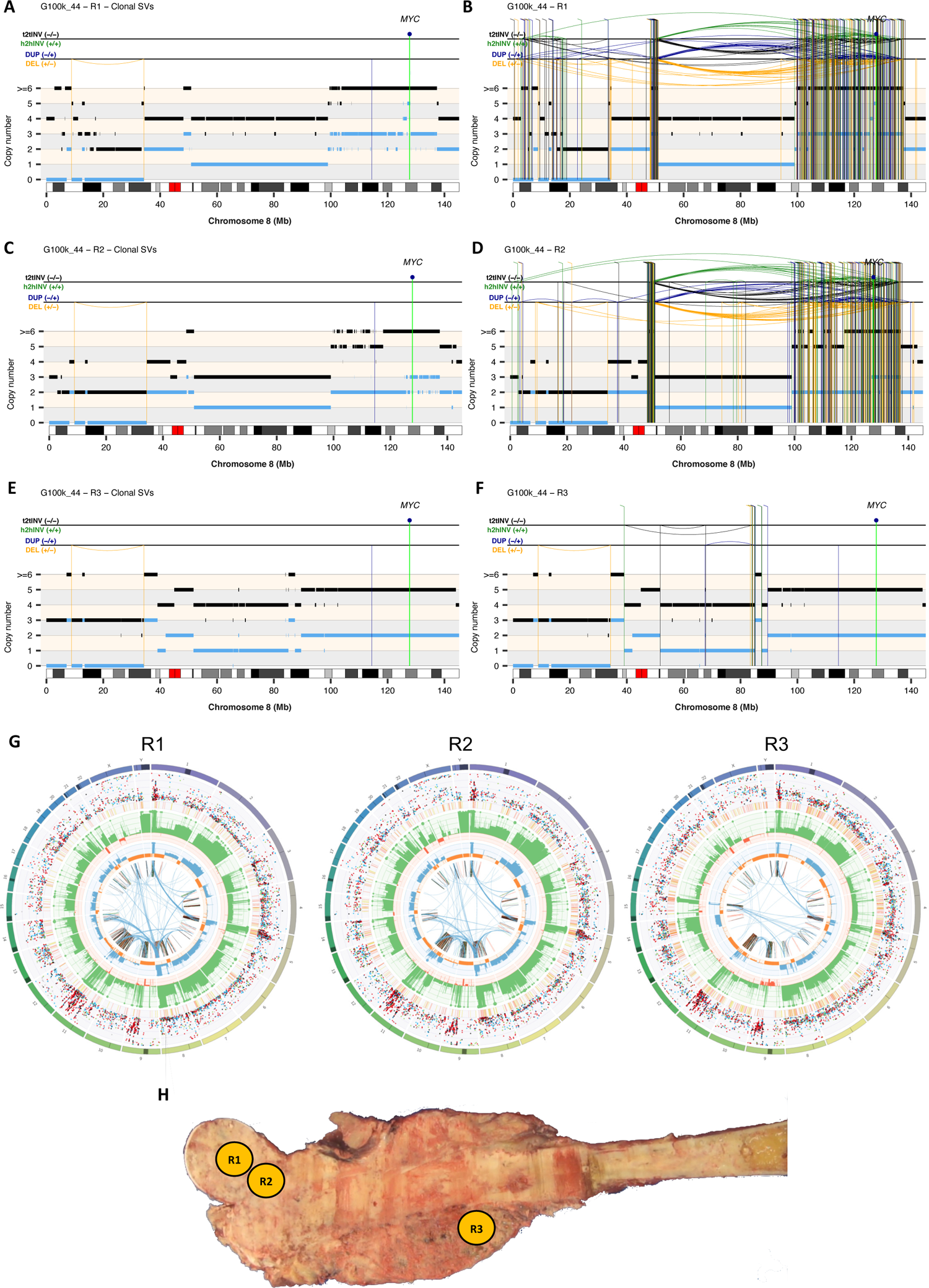
Evolutionary trajectory of chromosomes affected by CGRs in osteosarcoma G100k_44. Rearrangement profiles of chromosome 8 computed using somatic SV and copy-number information for tumour regions R1, R2 and R3 from case G100k_44. In **A**, **C** and **E**, only somatic SVs detected in the three regions are shown. All SVs detected in each region, including clonal SVs, are shown in **B**, **D** and **F.** The total and minor copy-number data in **A-F** are represented in black and blue, respectively. DEL, deletion-like rearrangement; DUP, duplication-like rearrangement; h2hINV, head-to-head inversion; t2tINV, tail-to-tail inversion. **(G)** Circos plots representing the landscape of somatic aberrations detected in WGS data from regions R1, R2 and R3. **(H)** Macroscopic picture of the tumour resection showing the regions profiled using WGS in this study. Sampling regions 1-3 are labelled.

**Fig. S10.**
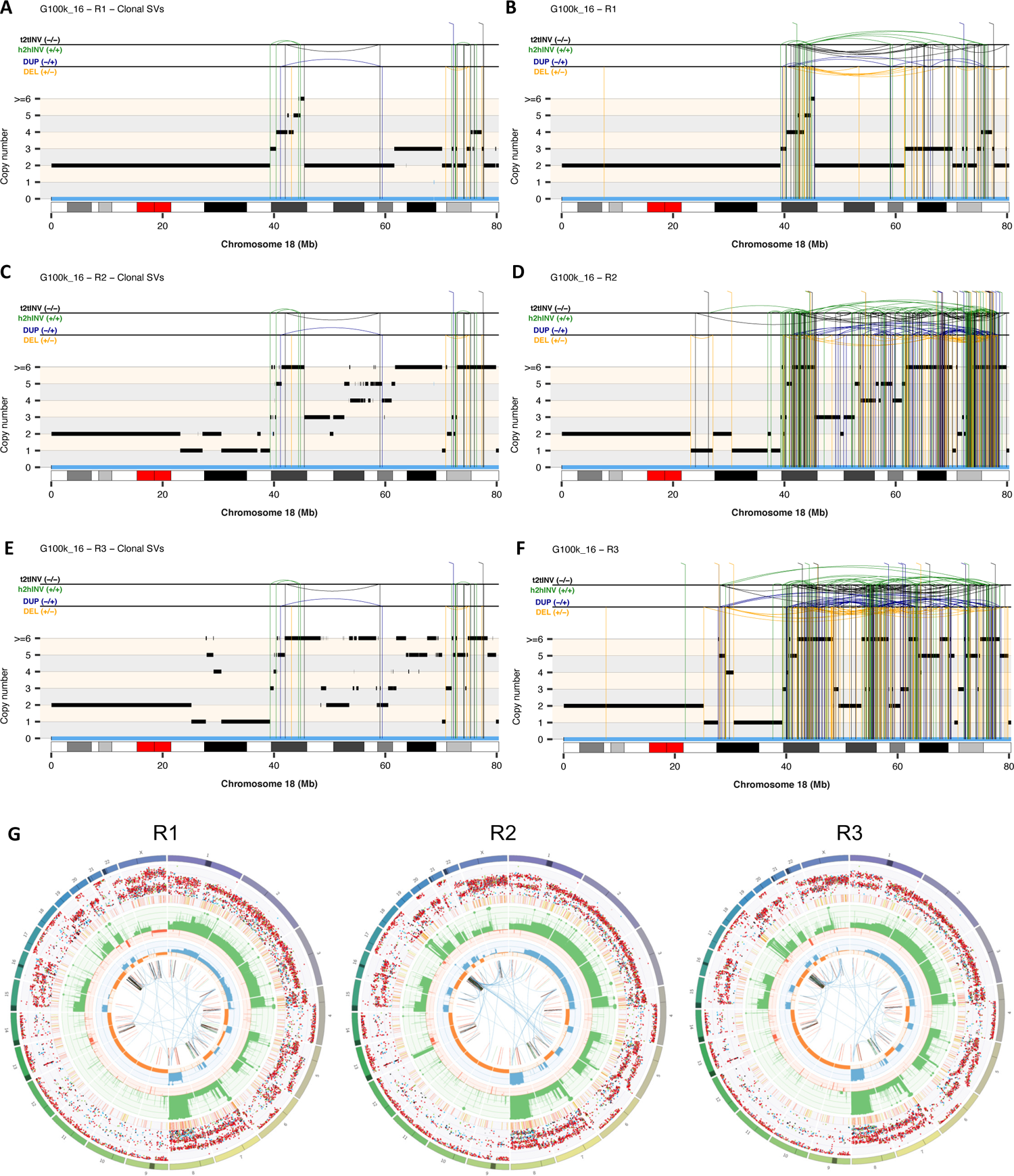
Evolutionary trajectory of chromosomes affected by CGRs in osteosarcoma G100k_16. Rearrangement profiles of chromosome 18 computed using somatic SV and copy-number information for tumour regions R1, R2 and R3 from case G100k_16. In **A**, **C** and **E**, only somatic SVs detected in the three regions are shown. All SVs detected in each region, including clonal SVs, are shown in **B**, **D** and **F.** The total and minor allele copy-number data in **A-F** are represented in black and blue, respectively. DEL, deletion-like rearrangement; DUP, duplication-like rearrangement; h2hINV, head-to-head inversion; t2tINV, tail-to-tail inversion. **(G)** Circos plots representing the landscape of somatic aberrations detected in WGS data from regions R1, R2 and R3.

**Fig. S11.**
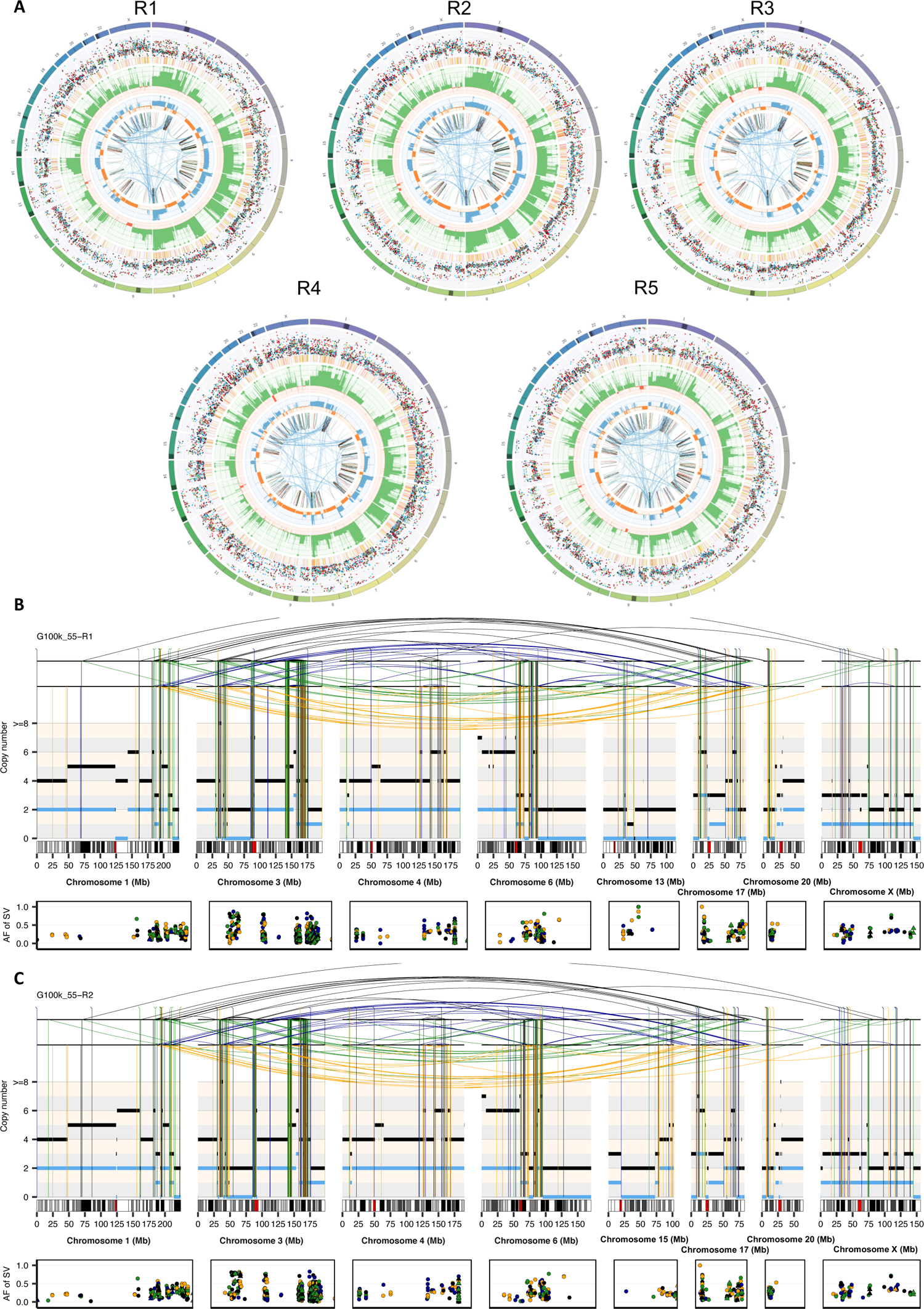
**(A)** Circos plots representing the landscape of somatic aberrations detected in multi-region WGS data for case G100k_55. Rearrangement profiles of chromosomes 1, 3, 4, 6, 15, 17, 20 and X using somatic copy-number and SV data from regions 1 (**B**) and 2 (**C**) for case G100k_55. The total and minor copy-number data in **B-C** are represented in black and blue, respectively. DEL, deletion-like rearrangement; DUP, duplication-like rearrangement; h2hINV, head-to-head inversion; t2tINV, tail-to-tail inversion.

**Fig. S12.**
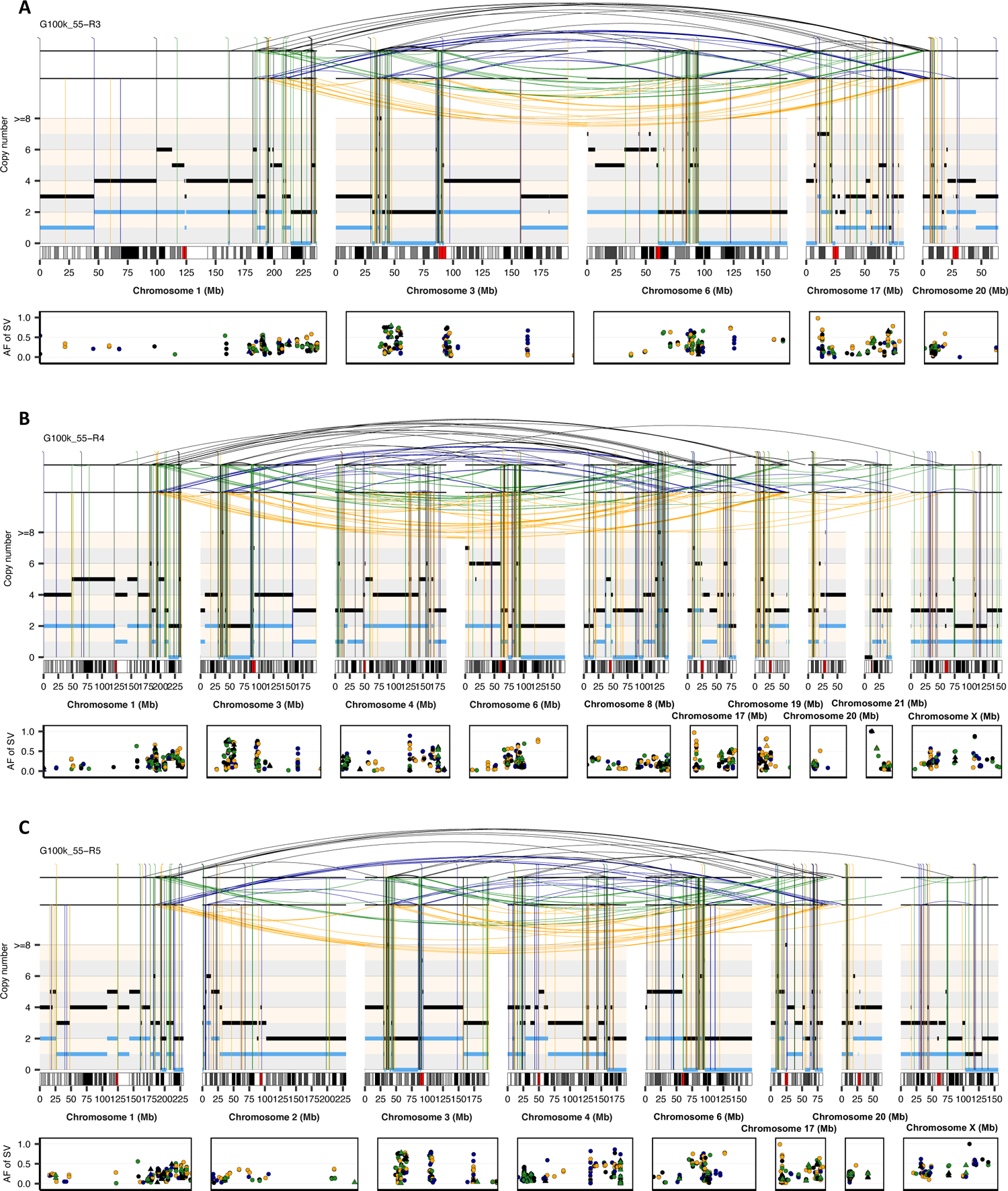
Rearrangement profiles of chromosomes 1, 3, 4, 6, 15, 17, 20 and X using somatic copy-number and SV data from regions 3 (**A**), 4 (**B**) and 5 (**C**) for case G100k_55. The total and minor allele copy-number data in **B-C** are represented in black and blue, respectively. DEL, deletion-like rearrangement; DUP, duplication-like rearrangement; h2hINV, head-to-head inversion; t2tINV, tail-to-tail inversion.

**Fig. S13.**
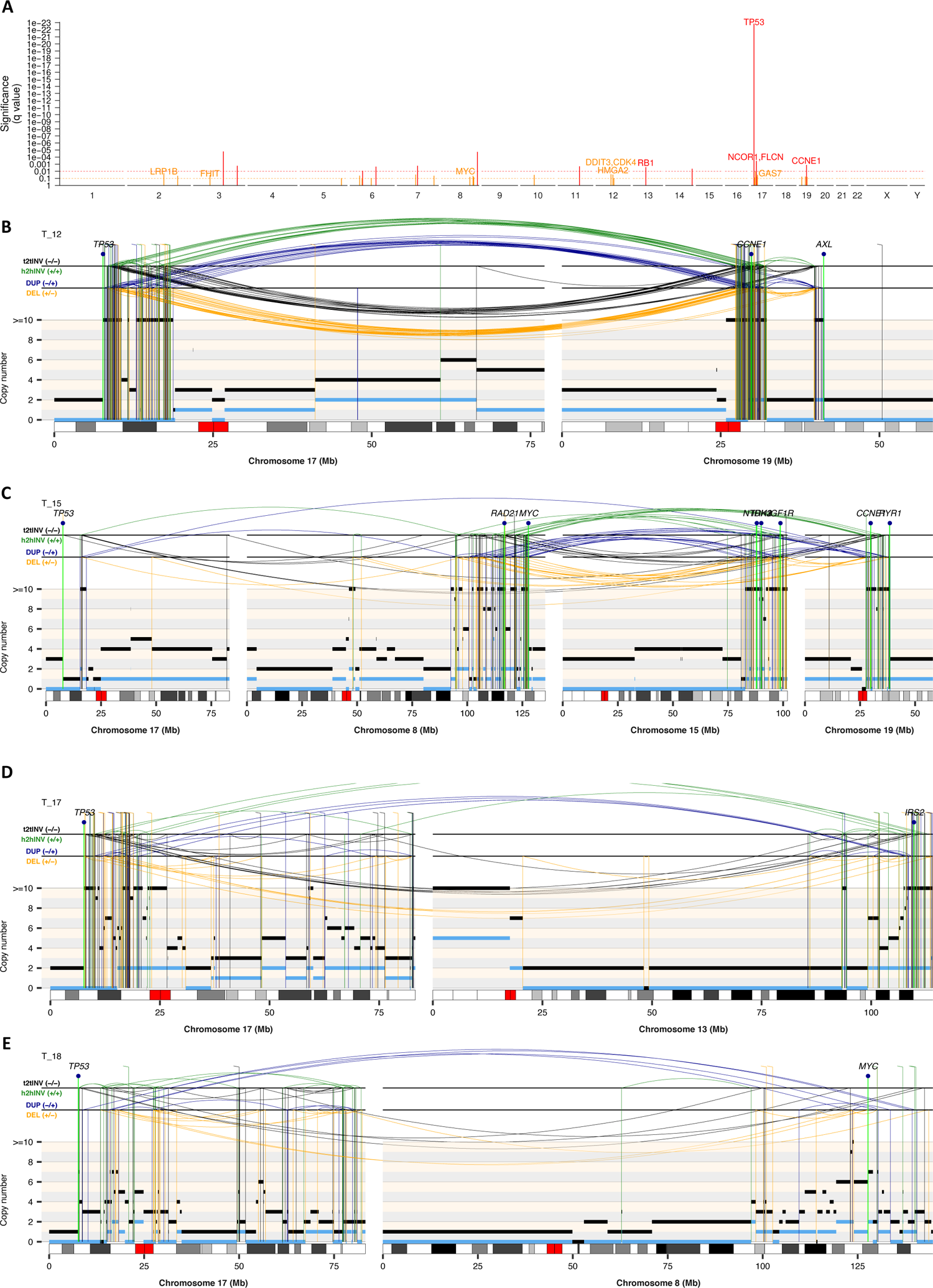
Genome-wide analysis of breakpoint enrichment and examples of LTA events. **(A)** Genome-wide enrichment analysis of breakpoints in high-grade osteosarcomas computed using non-overlapping windows of 50Kb. **(B-E)** Representative rearrangement profiles of cases from the TARGET (T) cohort showing loss of *TP53* and oncogene amplification via Loss-Translocation-Amplification mechanism. The total and minor copy-number data in **B-E** are represented in black and blue, respectively. DEL, deletion-like rearrangement; DUP, duplication-like rearrangement; h2hINV, head-to-head inversion; t2tINV, tail-to-tail inversion.

**Fig. S14.**
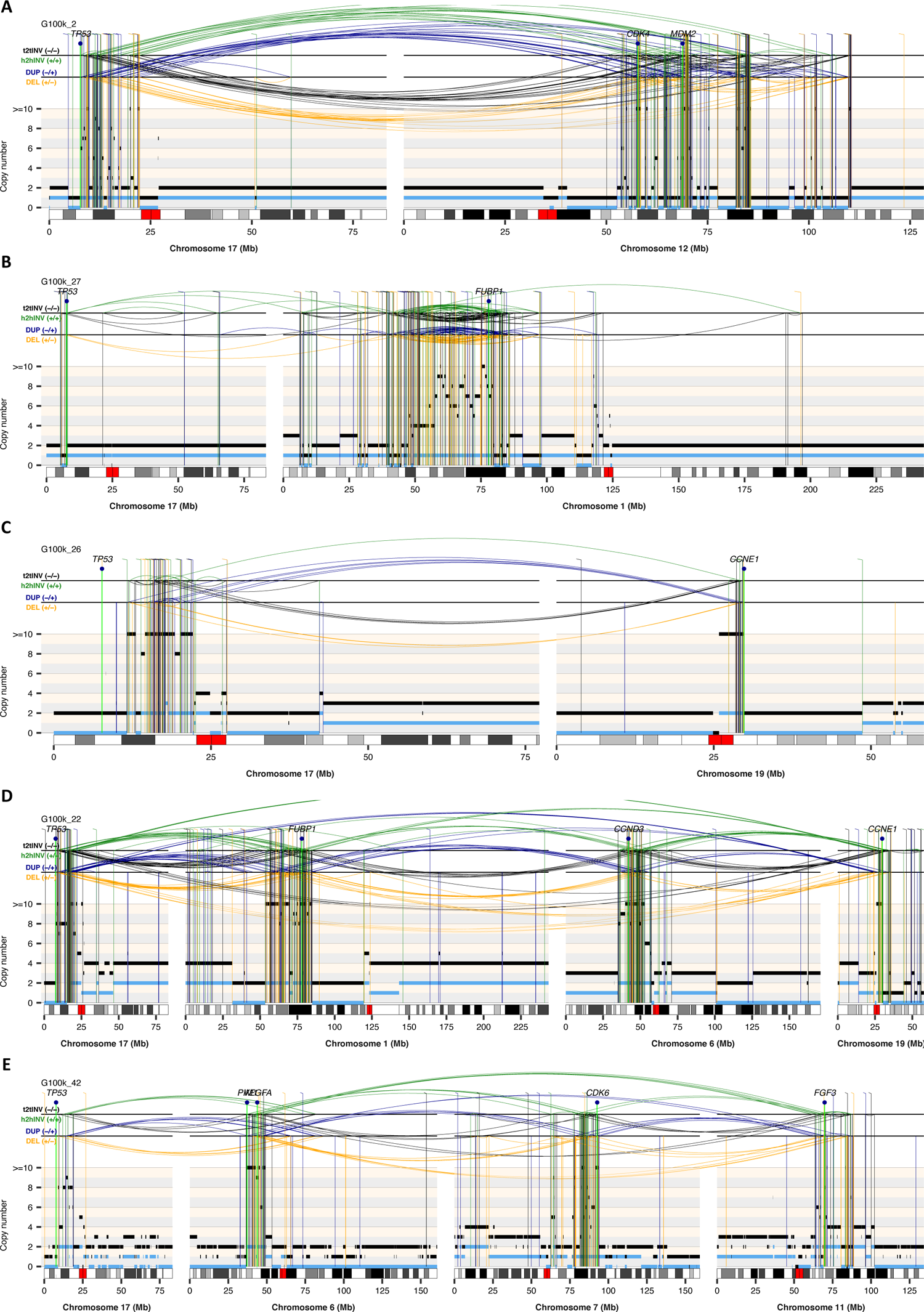
Representative rearrangement profiles of cases from the Genomics England 100,000 Genomes Project (G100k) cohort showing loss of *TP53* and oncogene amplification via LTA chromothripsis. The total and minor allele copy-number data in **A-E** are represented in black and blue, respectively. DEL,

**Fig. S15.**
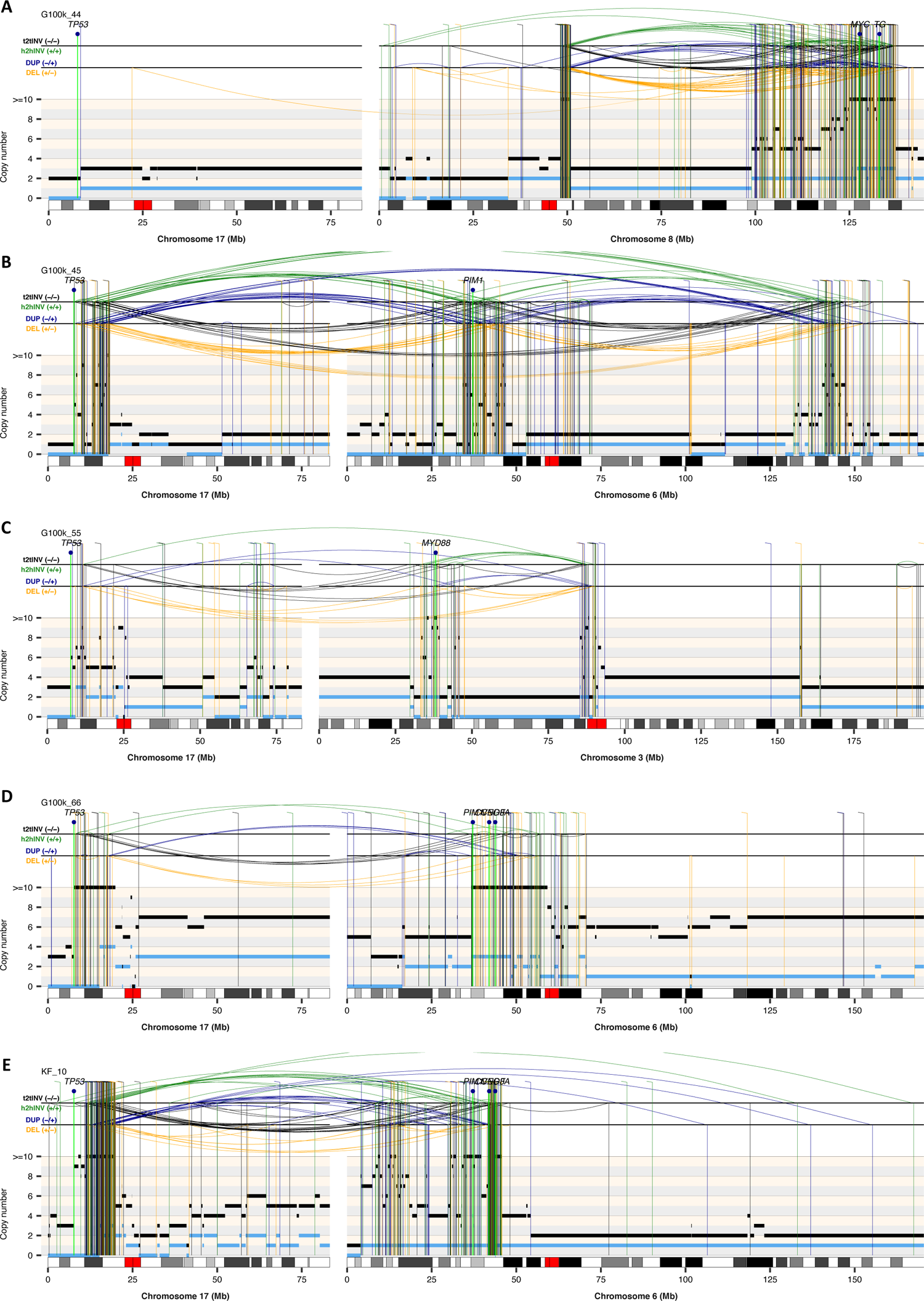
Representative rearrangement profiles of cases from the Genomics England 100,000 Genomes Project (G100k) and Kids First (KF) cohorts showing loss of *TP53* and oncogene amplification via LTA chromothripsis. The total and minor copy-number data in **A-E** are represented in black and blue, respectively.

**Fig. S16.**
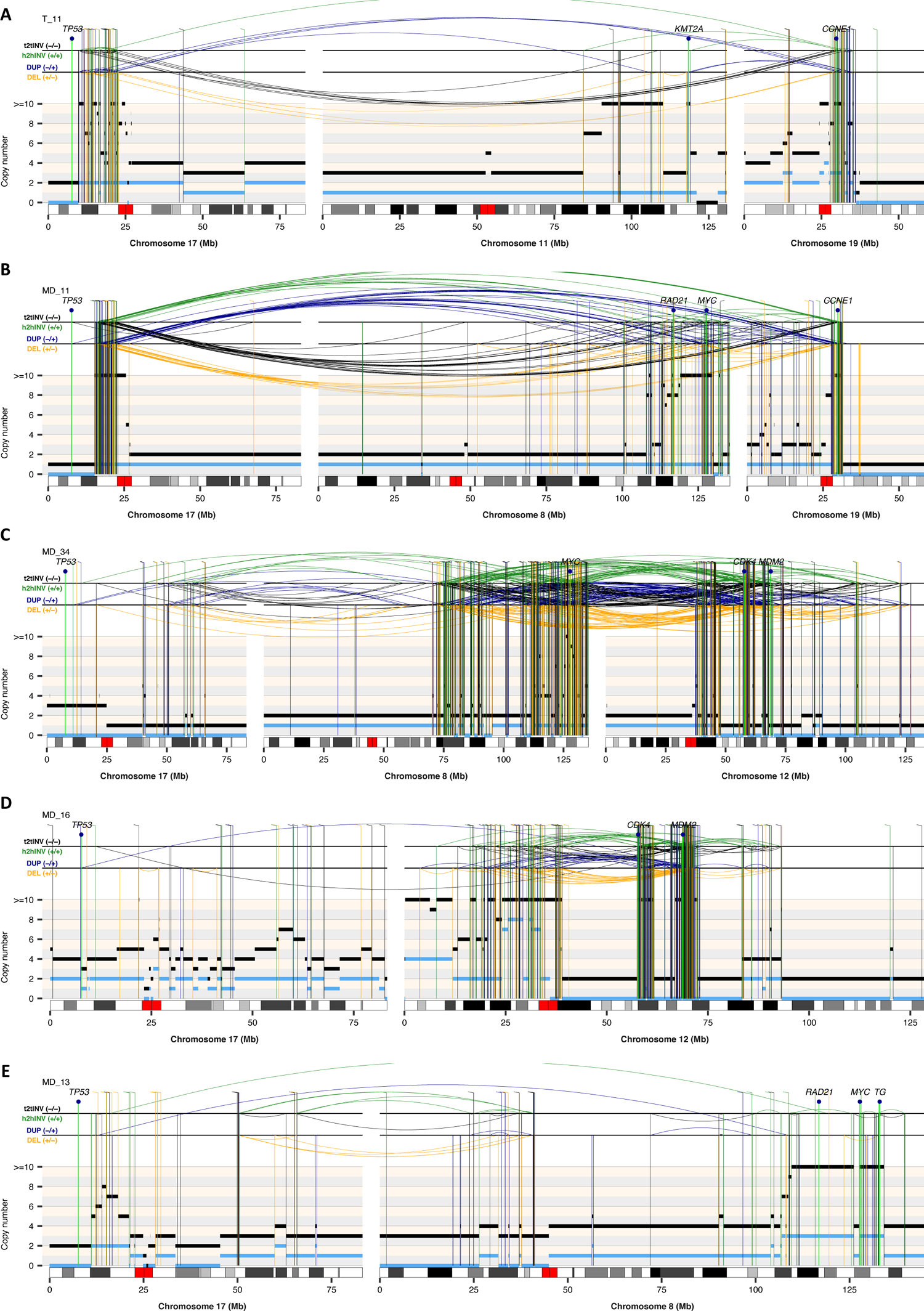
Representative rearrangement profiles of cases from the TARGET and MD (MD Anderson Cancer Center, Wu et al., *Nature Communications* 2020) cohorts showing loss of *TP53* and oncogene amplification via LTA chromothripsis. The total and minor allele copy-number data in **A-E** are represented in black and blue, respectively. DEL, deletion-like rearrangement; DUP, duplication-like rearrangement; h2hINV, head-to-head inversion; t2tINV, tail-to-tail inversion.

**Fig. S17.**
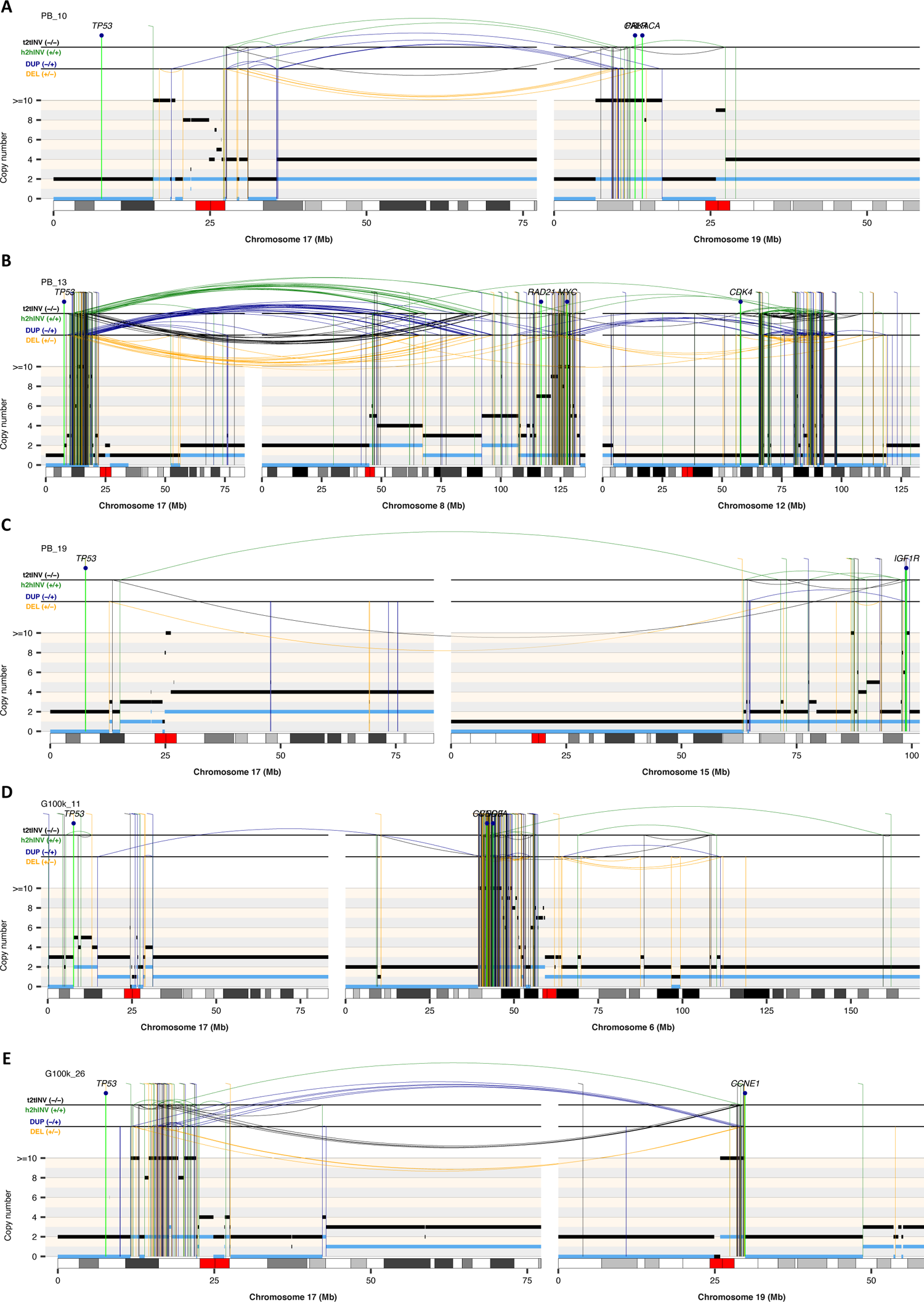
Representative rearrangement profiles of cases from the PB (PCAWG cases from Behjati et al. *Nature Communications*, 2017) cohort showing loss of *TP53* and oncogene amplification via LTA chromothripsis. The total and minor copy-number data in **A-E** are represented in black and blue, respectively. DEL, deletion-like rearrangement; DUP, duplication-like rearrangement; h2hINV, head-to-head inversion; t2tINV, tail-to-tail inversion.

**Fig. S18.**
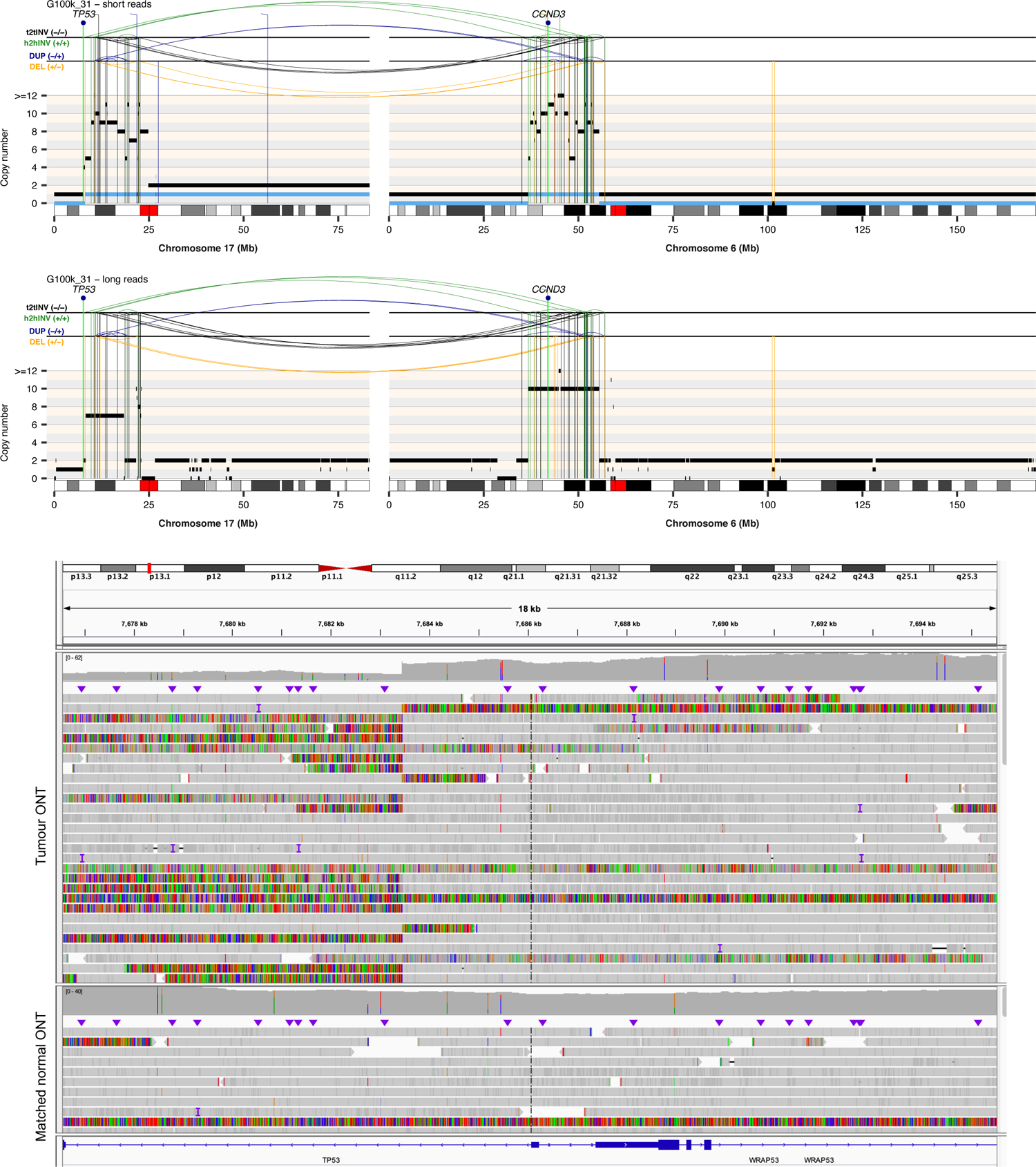
Rearrangement profiles of chromosome 17 computed using Illumina (**A**) and nanopore WGS (**B**) data for case G100k_31. The total and minor allele copy-number data in **A-B** are represented in black and blue, respectively. DEL, deletion-like rearrangement; DUP, duplication-like rearrangement; h2hINV, head-to-head inversion; t2tINV, tail-to-tail inversion. (**C**) Raw nanopore sequencing reads showing the breakpoint in *TP53* that triggers LTA in case G100k_31.

**Fig. S19.**
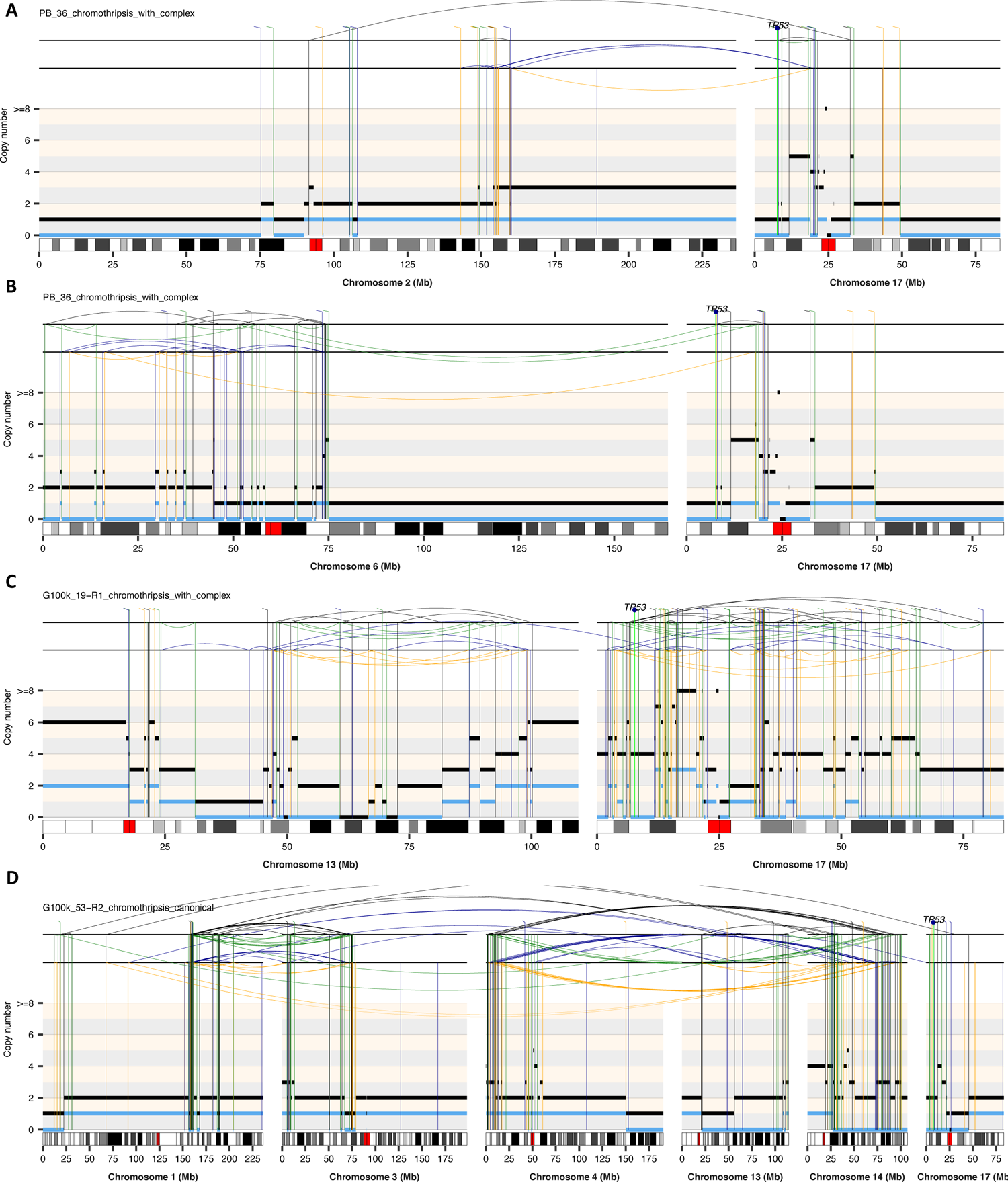
Representative rearrangement profiles of cases showing loss of *TP53* via LTA chromothripsis and segmental amplifications not overlapping recurrently amplified oncogenes. The total and minor copy-number data in **A-D** are represented in black and blue, respectively. DEL, deletion-like rearrangement; DUP, duplication-like rearrangement; h2hINV, head-to-head inversion; t2tINV, tail-to-tail inversion.

**Fig. S20.**
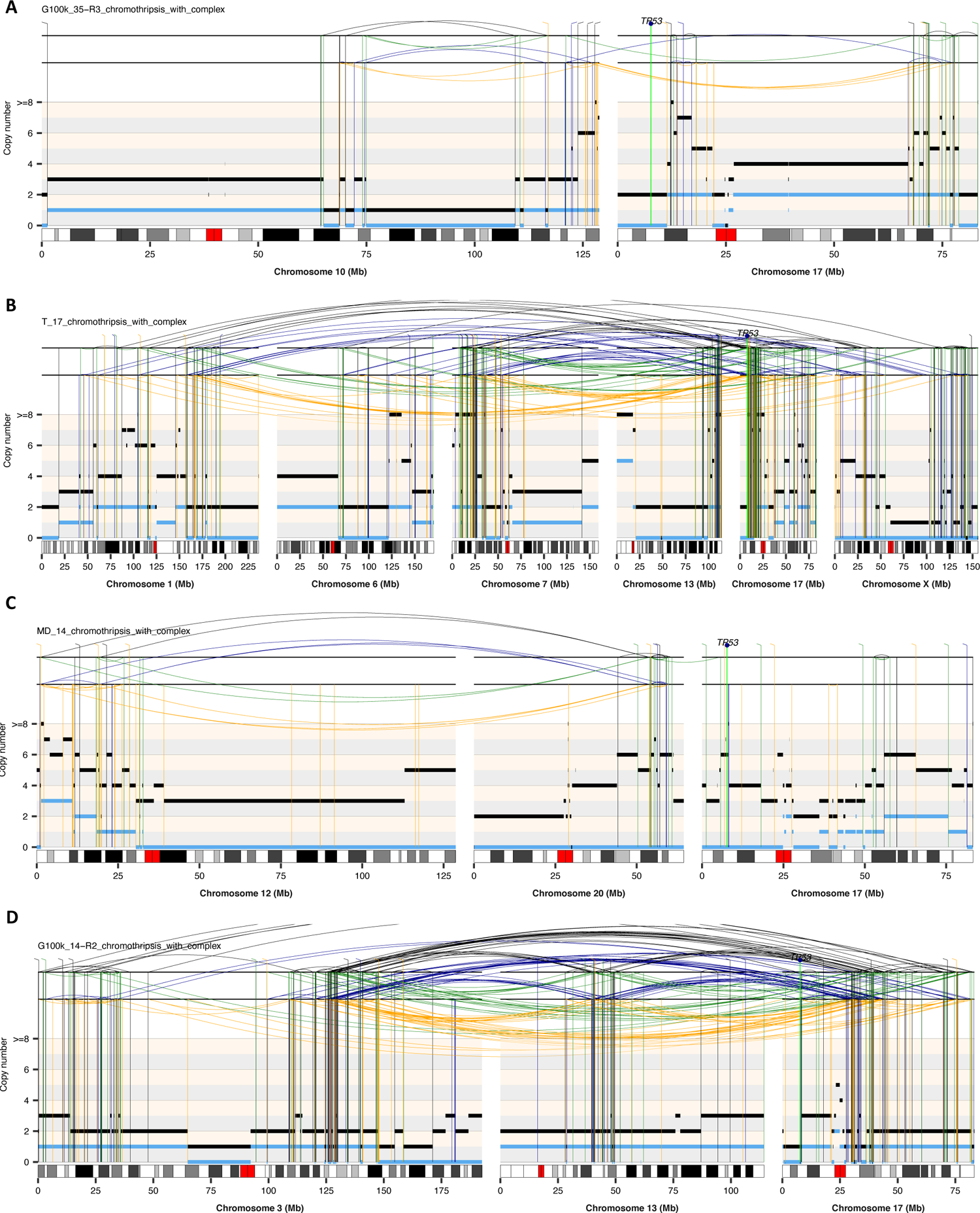
Representative rearrangement profiles of cases showing loss of *TP53* via LTA chromothripsis and segmental amplifications not overlapping recurrently amplified oncogenes. The total and minor allele copy-number data in **A-D** are represented in black and blue, respectively. DEL, deletion-like rearrangement; DUP, duplication-like rearrangement; h2hINV, head-to-head inversion; t2tINV, tail-to-tail inversion.

**Fig. S21.**
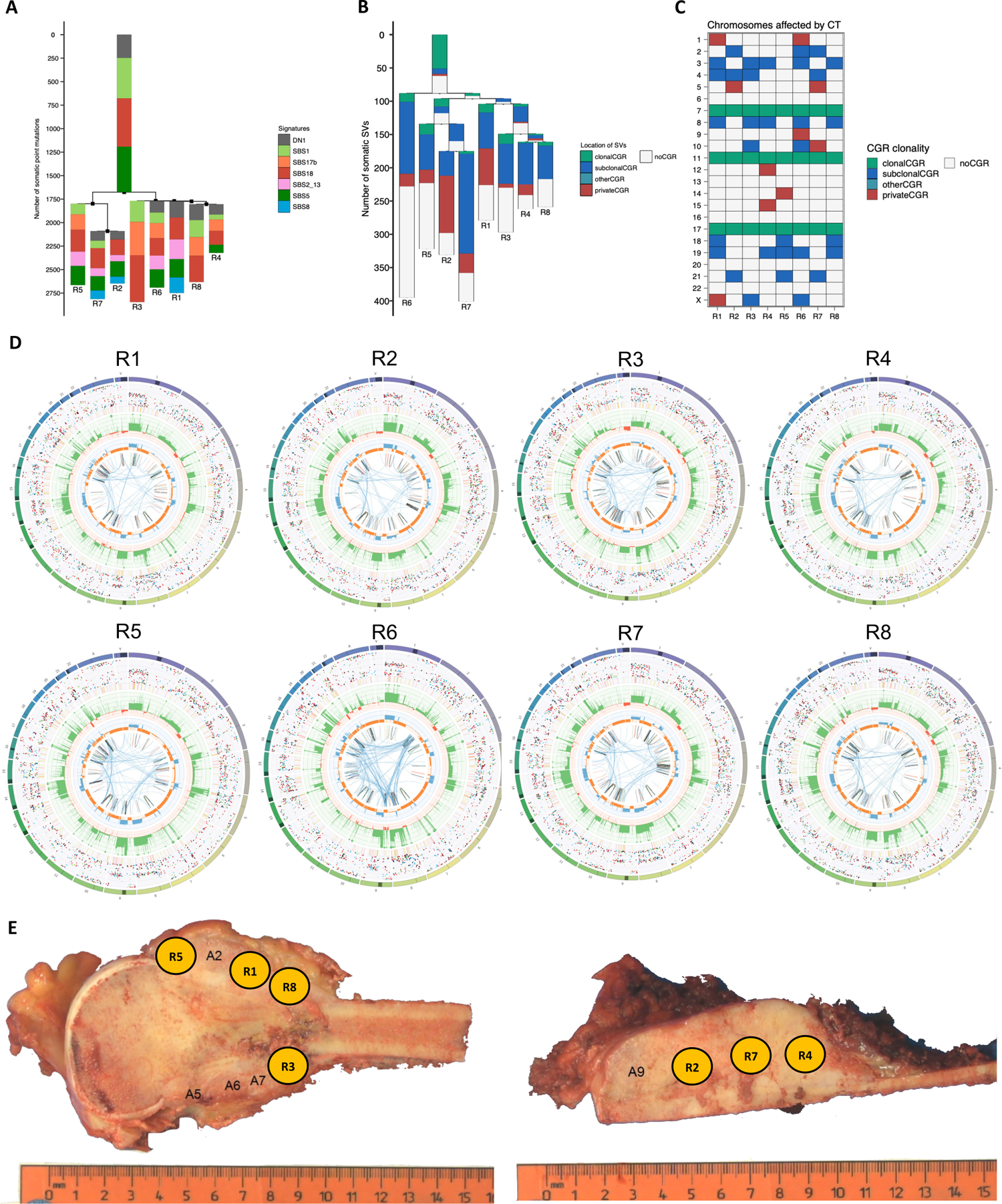
Intra-tumour heterogeneity in high-grade osteosarcoma G100k_30. Phylogenetic trees constructed using somatic SNVs (**A**) or SVs (**B**) detected in multi-region WGS data from case G100k_30. The colours of the bars in the SNV tree represent the contribution of different SBS mutational signatures to the SNVs detected. The colours in the SV tree indicate whether the SVs in the tree map to clonal, subclonal or private CGRs (green, blue, and red), respectively, or outside CGR regions (grey). (C) Distribution of clonal, subclonal and private CGRs across chromosomes and tumour regions. (D) Circos plots representing the landscape of somatic aberrations detected in the WGS data from 8 tumour regions (R) case G100k_30.(E) Macroscopic image of the primary tumour from case G100k_30. Sampling regions 1-8 are labelled. Region 6 corresponds to a biopsy sample, whose location within the primary tumour was not recorded. The side of the ruler is marked in centimeters.

**Fig. S22.**
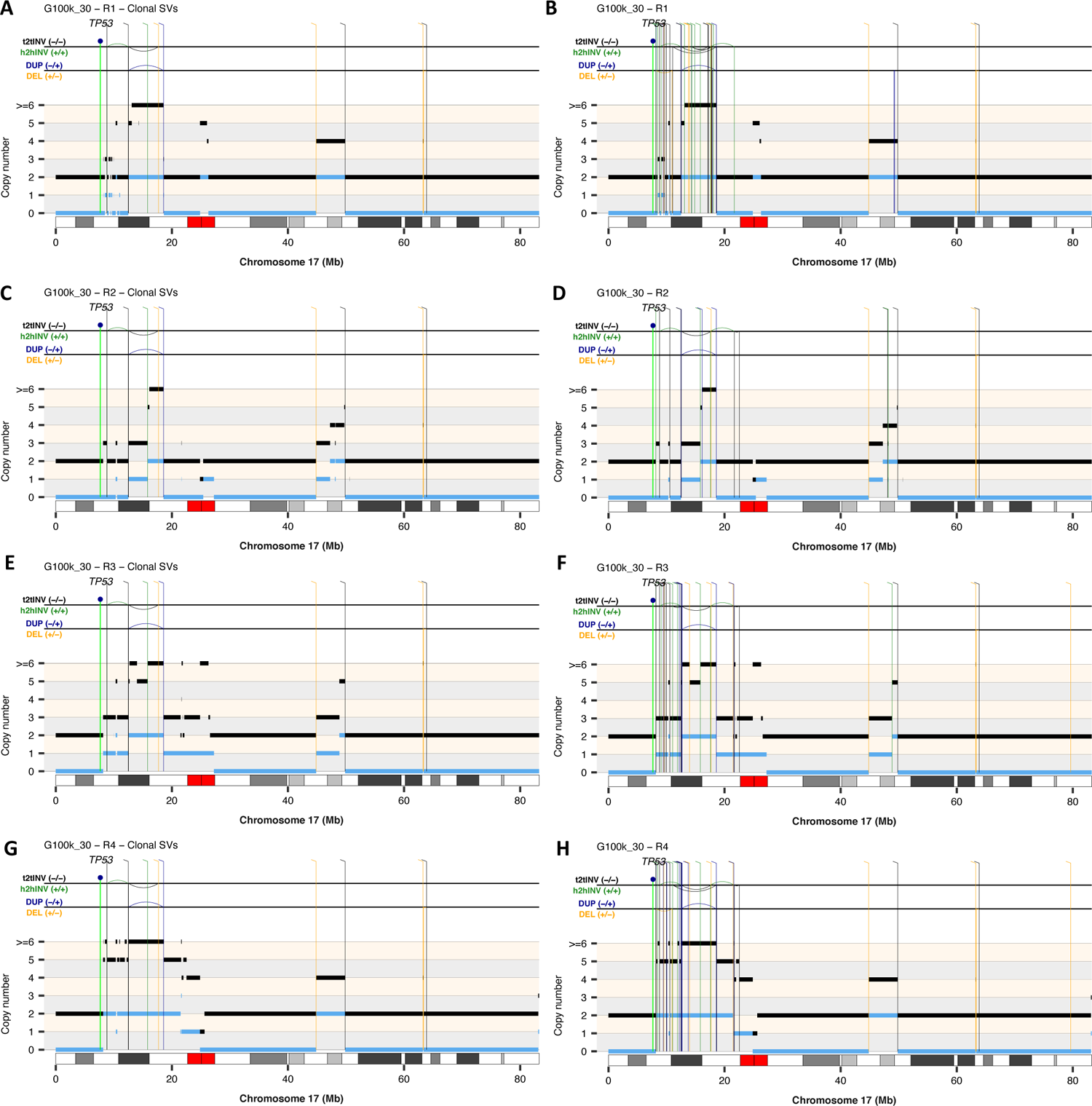
Evolutionary trajectory of chromosomes affected by CGRs in osteosarcoma G100k_30. Rearrangement profiles of chromosome 17 computed using somatic SV and copy-number information for tumour regions R1, R2, R3 and R4 from case G100k_30. In **A**, **C, E** and **G**, only somatic SVs detected in the eight regions from G100k_30 analysed are shown. All SVs detected in each region, including clonal SVs, are shown in **B**, **D, F** and **H.** The total and minor copy-number data in **A-H** are represented in black and blue, respectively. DEL, deletion-like rearrangement; DUP, duplication-like rearrangement; h2hINV, head-to-head inversion; t2tINV, tail-to-tail inversion.

**Fig. S23.**
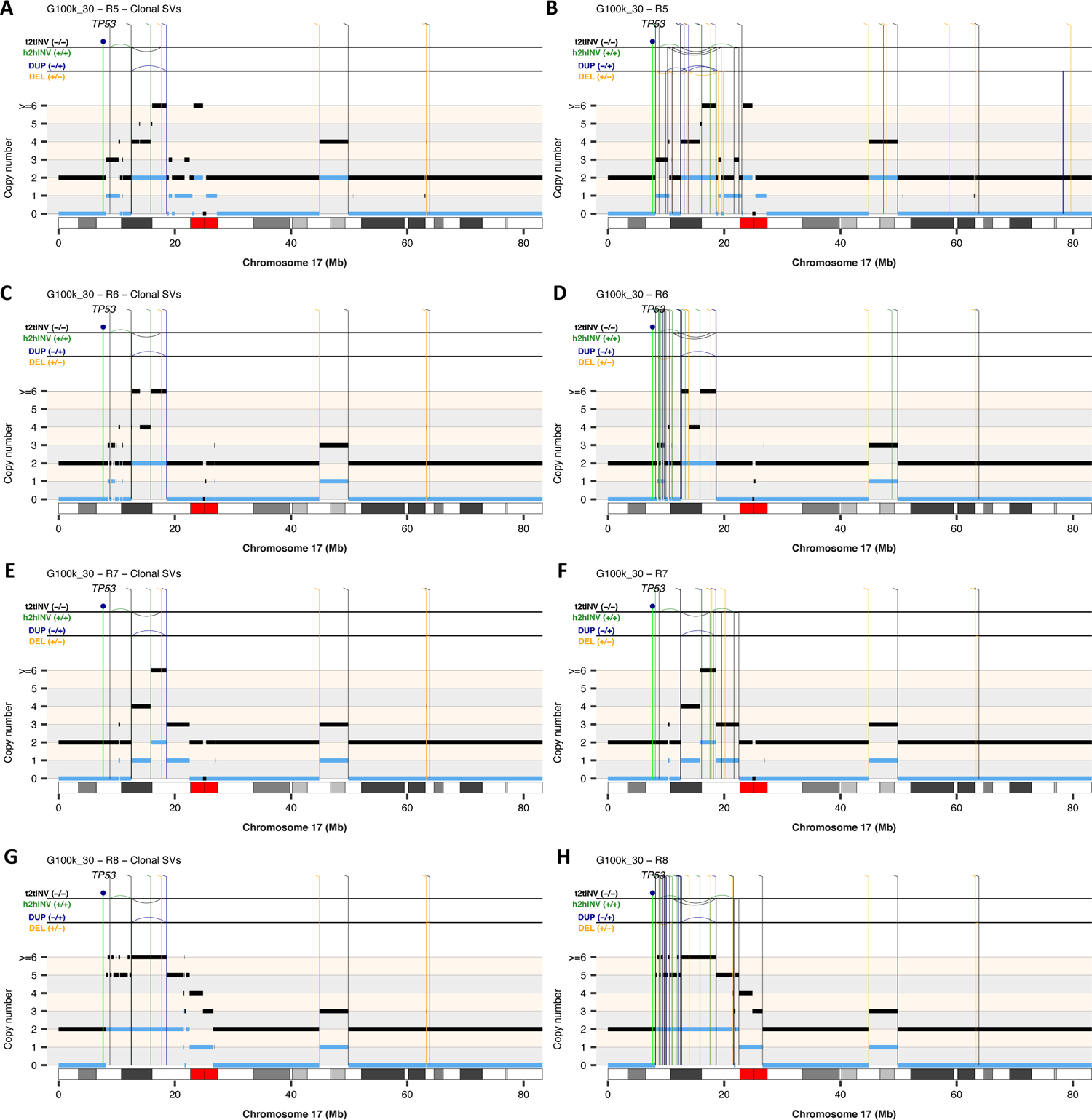
Evolutionary trajectory of chromosomes affected by CGRs in osteosarcoma G100k_30. Rearrangement profiles of chromosome 17 computed using somatic SV and copy-number information for tumour regions R5, R6, R7 and R8 from case G100k_30. In **A**, **C, E** and **G**, only somatic SVs detected in the eight regions from G100k_30 analysed are shown. All SVs detected in each region, including clonal SVs, are shown in **B**, **D, F** and **H.** The total and minor allele copy-number data in **A-H** are represented in black and blue, respectively. DEL, deletion-like rearrangement; DUP, duplication-like rearrangement; h2hINV, head-to-head inversion; t2tINV, tail-to-tail inversion.

**Fig. S24.**
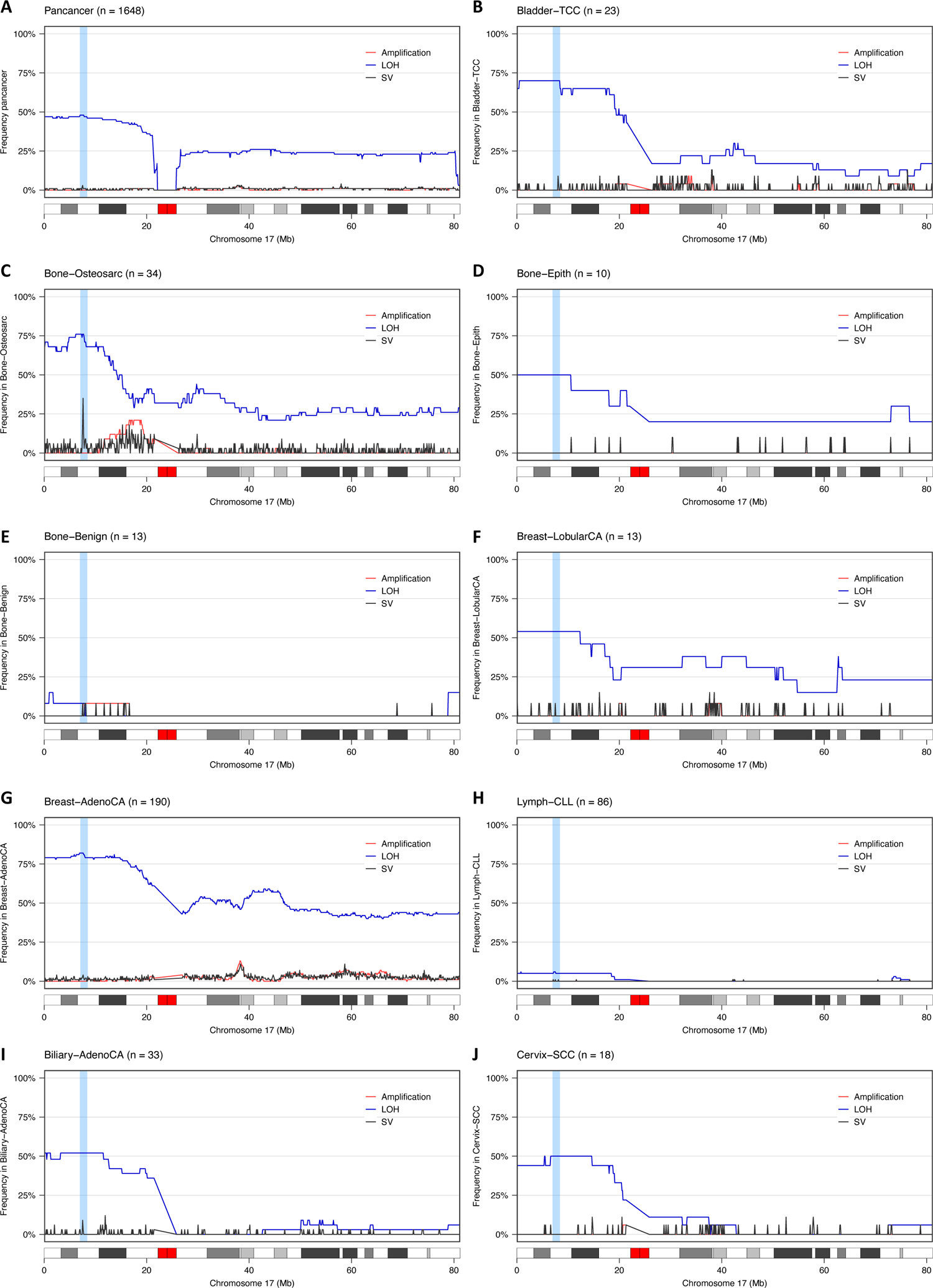
Pan-cancer analysis of rearrangements and copy number aberrations in chromosome 17. Analysis of the frequency of amplification (red), LOH (blue) and structural variants in chromosome 17 across cancer types in the ICGC/TCGA PCAWG cohort with at least 10 tumour samples computed using non-overlapping windows of 100 kilobases. (**A**) Pan-cancer analysis, **(B**) Bladder transitional cell carcinoma (Bladder-TCC), (**C**) Bone osteosarcoma (Bone-Osteosarcoma), (**D**) bone neoplasm, epithelioid (Bone-Epith), (**E**) bone cartilaginous neoplasm, osteoblastoma and bone osteofibrous dysplasia (Bone-Benign), (**F**) Breast lobular carcinoma (Breast-LobCA), (**G**) Breast adenocarcinoma (Breast-AdenoCA), (**H**) Lymphoid chronic lymphocytic leukemia (Lymph. CLL), (**I**) Billiary adenocarcinoma (Billiary-AdenoCA), and (**J**) cervix squamous cell carcinoma (Cervix-SCC).

**Fig. S25.**
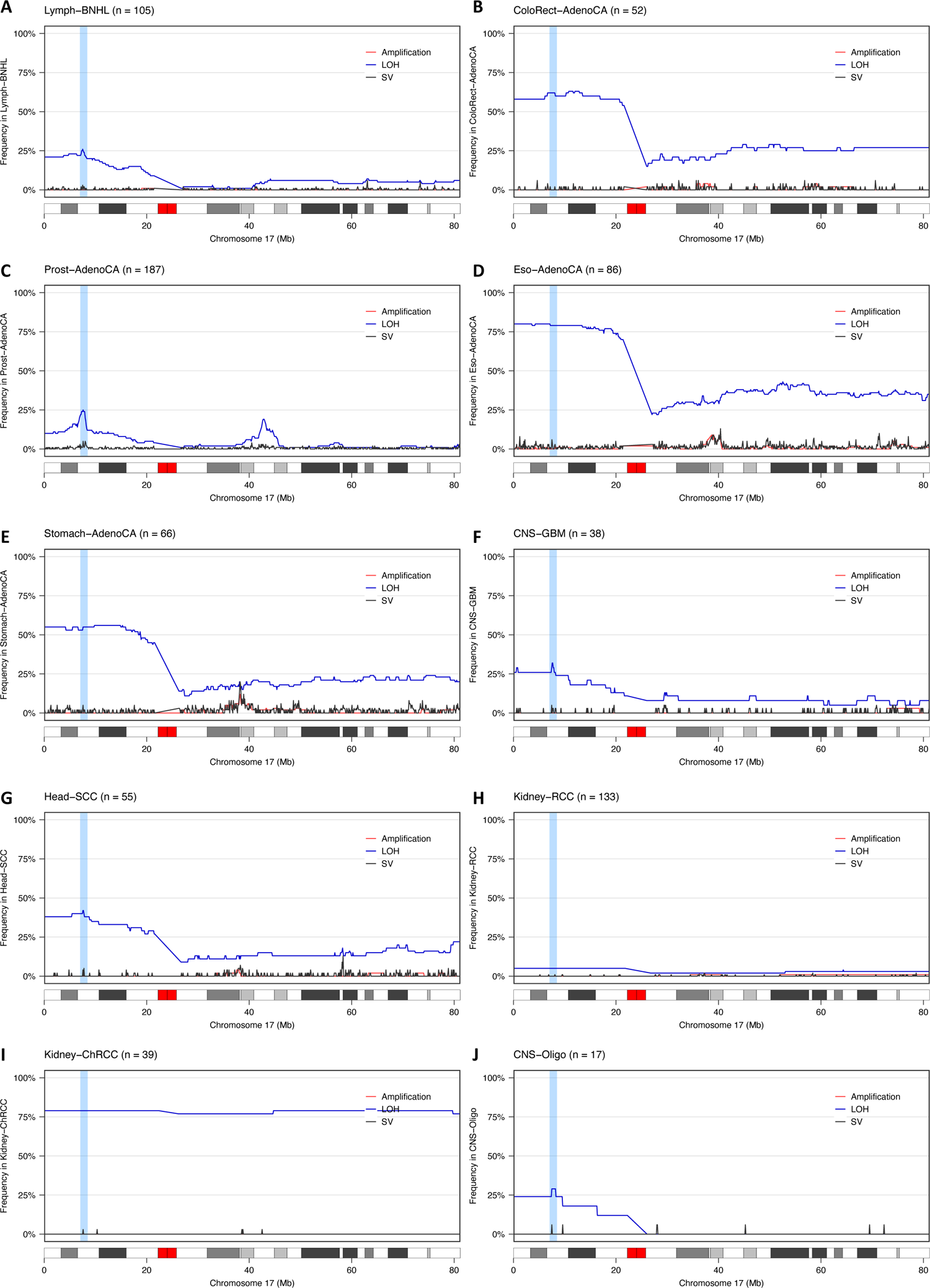
Pan-cancer analysis of rearrangements and copy number aberrations in chromosome 17. Analysis of the frequency of amplification (red), LOH (blue) and structural variants in chromosome 17 across cancer types in the ICGC/TCGA PCAWG cohort with at least 10 tumour samples computed using non-overlapping windows of 100 kilobases. (**A**) lymphoid mature B-cell lymphoma (Lymph-BNHL), (**B**) colorectal adenocarcinoma (ColoRect-AdenoCA), (**C**) prostate adenocarcinoma (Prost-AdenoCA), (**D**) esophagus adenocarcinoma (Eso-AdenoCA), (**E**) stomach adenocarcinoma (Stomach-AdenoCA), (**F**) central nervous system glioblastoma (CNS-GBM), (**G**) head-and-neck squamous cell carcinoma (Head-SCC), (**H**) kidney renal cell carcinoma (Kidney-RCC), (**I**) kidney chromophobe renal cell carcinoma (Kidney-ChRCC), and (**J**) CNS oligodenroglioma (CNS-Oligo).

**Fig. S26.**
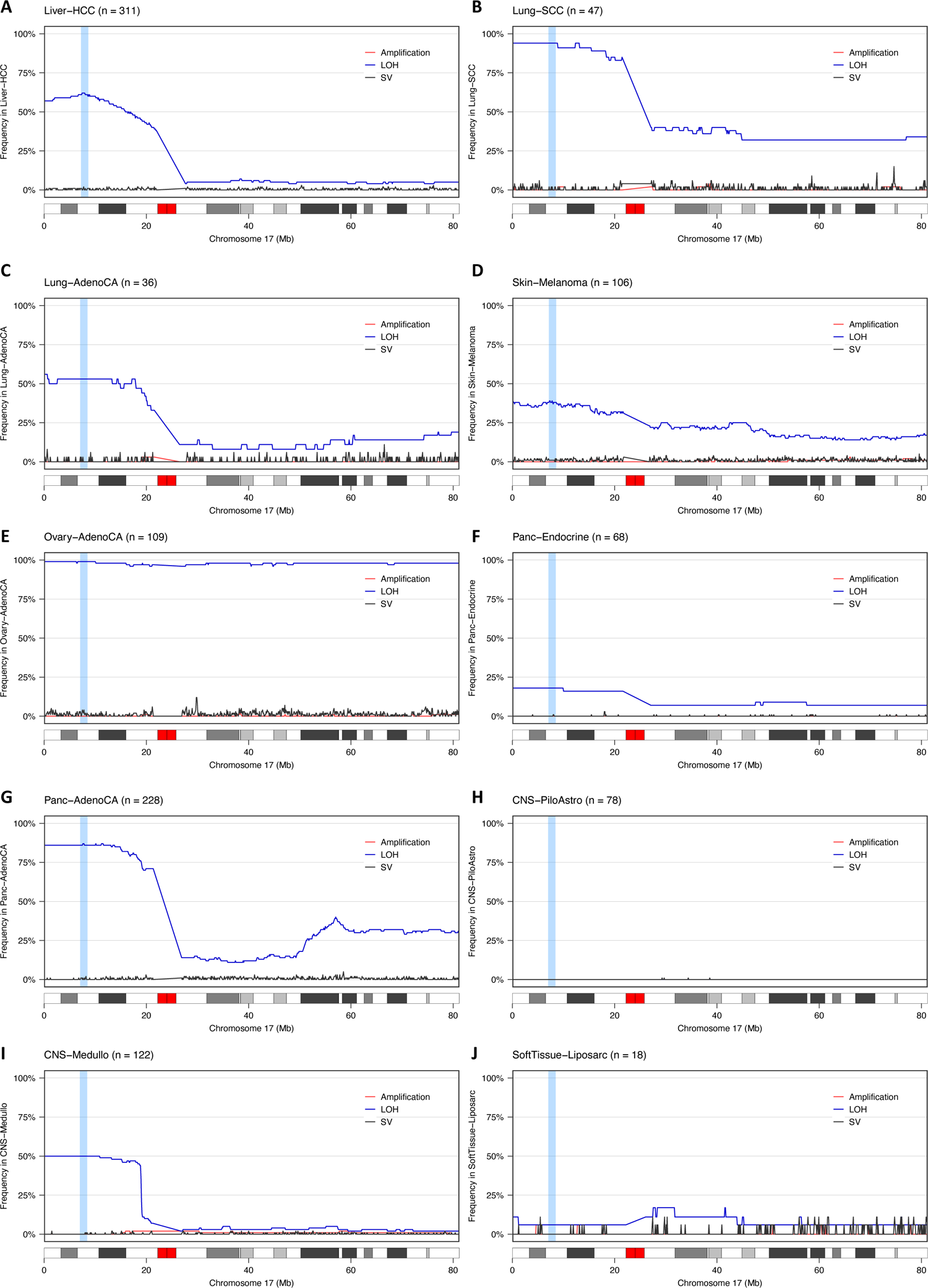
Pan-cancer analysis of rearrangements and copy number aberrations in chromosome 17. Analysis of the frequency of amplification (red), LOH (blue) and structural variants in chromosome 17 across cancer types in the ICGC/TCGA PCAWG cohort with at least 10 tumour samples computed using non-overlapping windows of 100 kilobases. (**A**) liver hepatocellular carcinoma (Liver-HCC), (**B**) lung squamous cell carcinoma (Lung-SCC), (**C**) lung adenocarcinoma (Lung-AdenoCA), (**D**) skin melanoma (Skin-Melanoma), (**E**) ovary adenocarcinoma (Ovary-AdenoCA), (**F**) pancreatic neuroendocrine tumor (Panc-Endocrine), (**G**) pancreatic adenocarcinoma (Panc-AdenoCA), (**H**) CNS pilocytic astrocytoma (CNS-PiloAstro), (**I**) CNS medulloblastoma (CNS-Medullo), and (**J**) liposarcoma (SoftTissue-Liposarc).

**Fig. S27.**
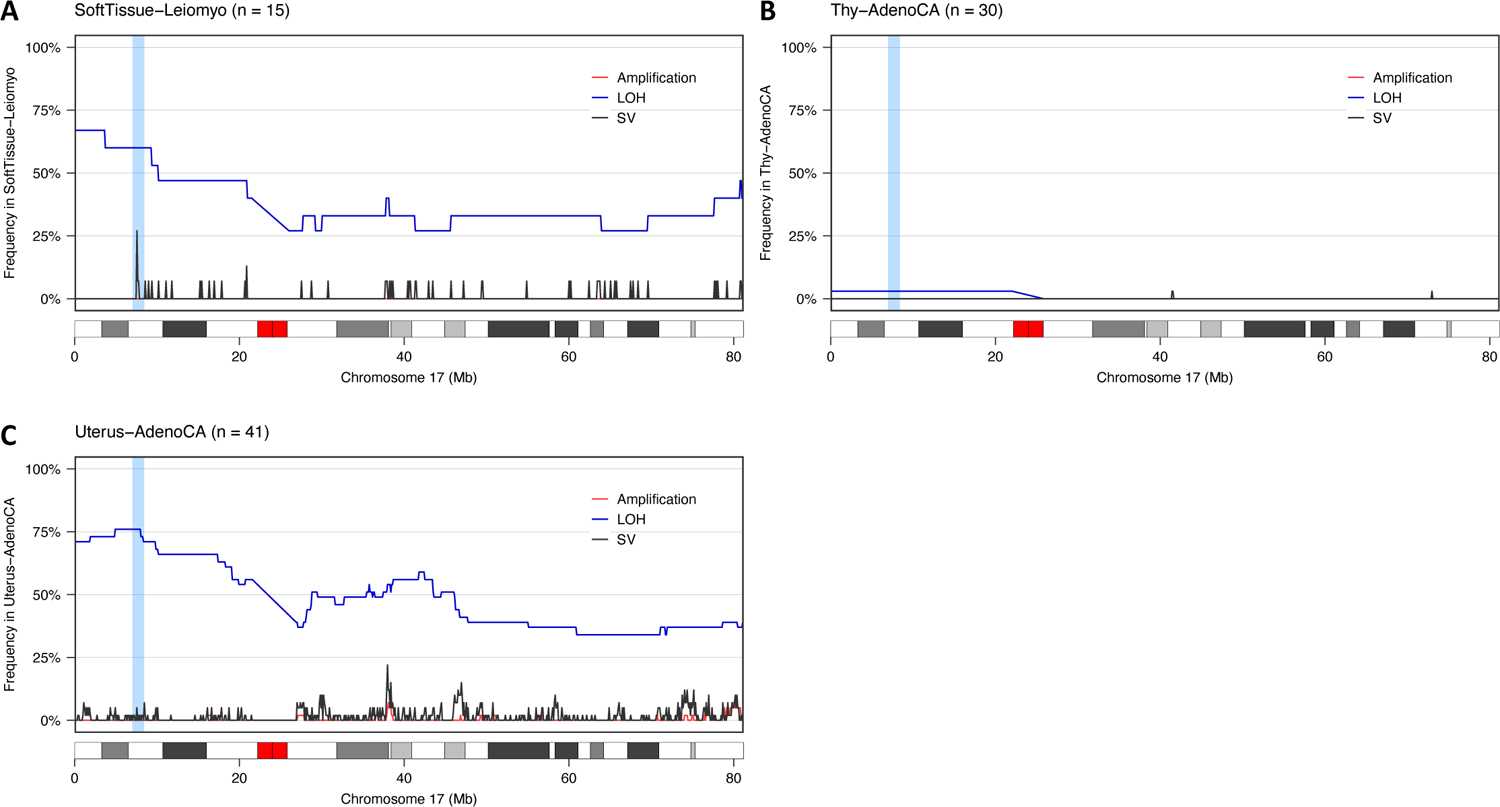
Pan-cancer analysis of rearrangements and copy number aberrations in chromosome 17. Analysis of the frequency of amplification (red), LOH (blue) and structural variants in chromosome 17 across cancer types in the ICGC/TCGA PCAWG cohort with at least 10 tumour samples computed using non-overlapping windows of 100 kilobases. (**A**) leiomyosarcoma, soft tissue (SoftTissue-Leiomyo), (**B**) thyroid low-grade adenocarcinoma (Thy-AdenoCA), and (**C**) uterus adenocarcinoma (Uterus-AdenoCA).

**Fig. S28.**
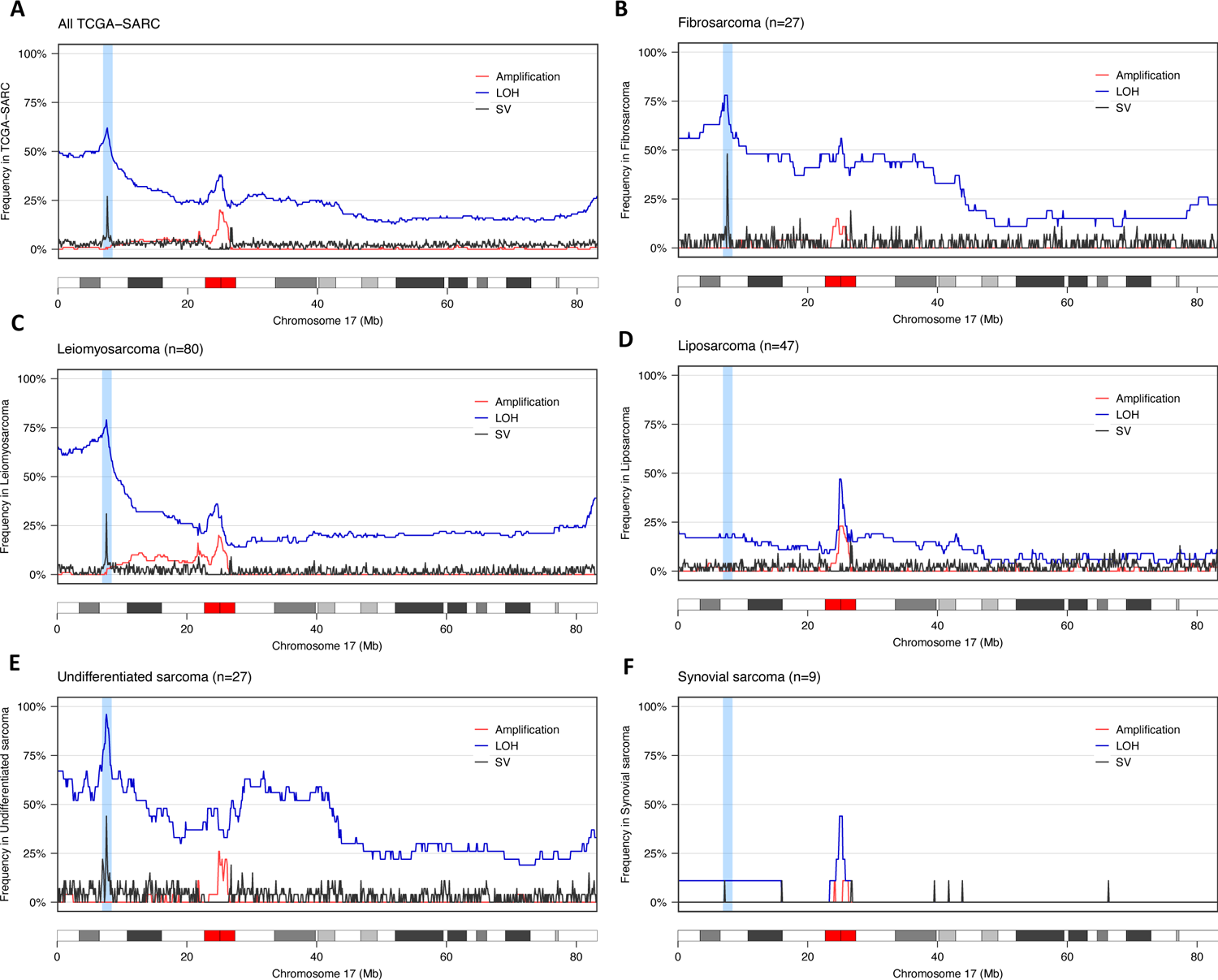
Analysis of rearrangements and copy number aberrations in chromosome 17 in sarcomas from TCGA (TCGA-SARC). Analysis of the frequency of amplification (red), LOH (blue) and structural variants in chromosome 17 across sarcoma subtypes from TCGA (TCGA-SARC) computed using non-overlapping windows of 100 kilobases. Only subtypes with whole-genome sequencing data for at least 5 tumour samples are shown. (**A**) All sarcomas, (**B**) fibrosarcomas, (**C**) leiomyosarcomas, (**D**) liposarcomas, (**E**) undifferentiated, and (**F**) synovial sarcoma.

**Fig. S29.**
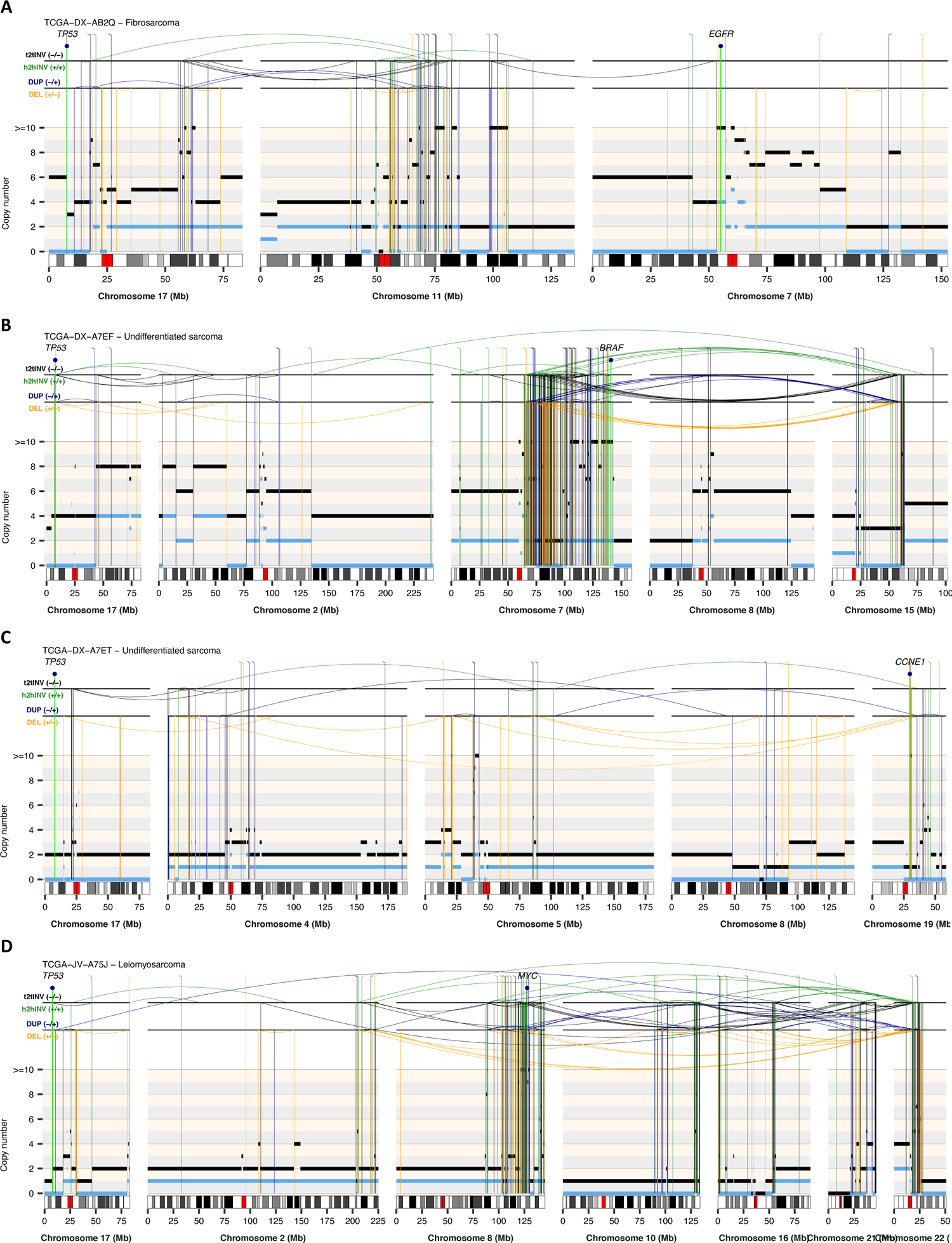
Representative rearrangement profiles of soft-tissue tumours from the TCGA-SARC cohort showing LTA chromothripsis. The total and minor copy-number data in **A-D** are represented in black and blue, respectively. DEL, deletion-like rearrangement; DUP, duplication-like rearrangement; h2hINV, head-to-head inversion; t2tINV, tail-to-tail inversion.

**Fig. S30.**
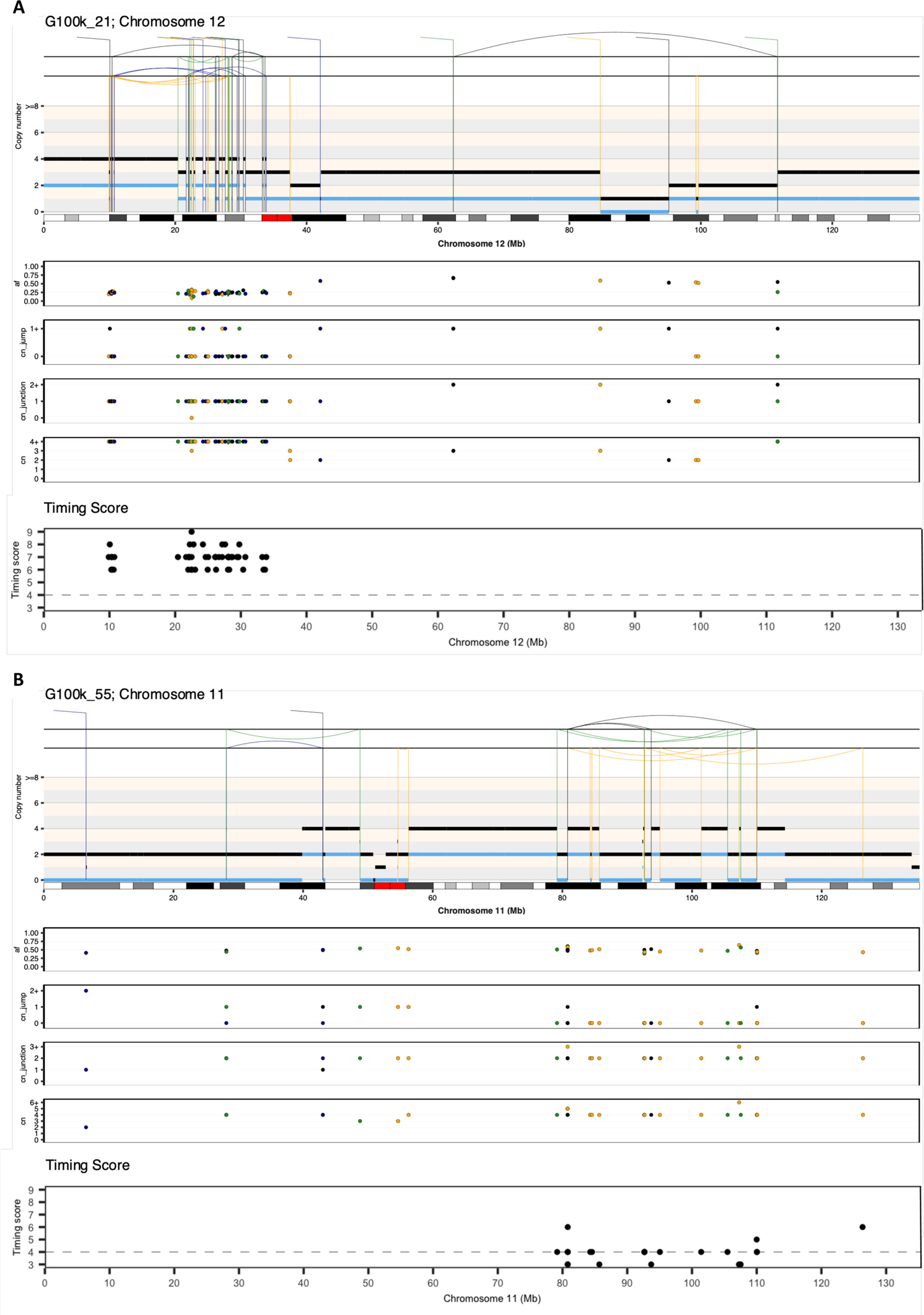
Example of canonical chromothripsis cases used to estimate the cut-off value for the timing score used to time fold-back inversions relative to WGD. (A) Example of canonical chromothripsis occurring after WGD, as evidenced by the total copy number oscillations (shown in black) between copy-number states 3 and 4. **(B)** Example of canonical chromothripsis occurring before WGD, as indicated by the presence of total copy number oscillations between copy-number states 2 and 4.

**Fig. S31.**
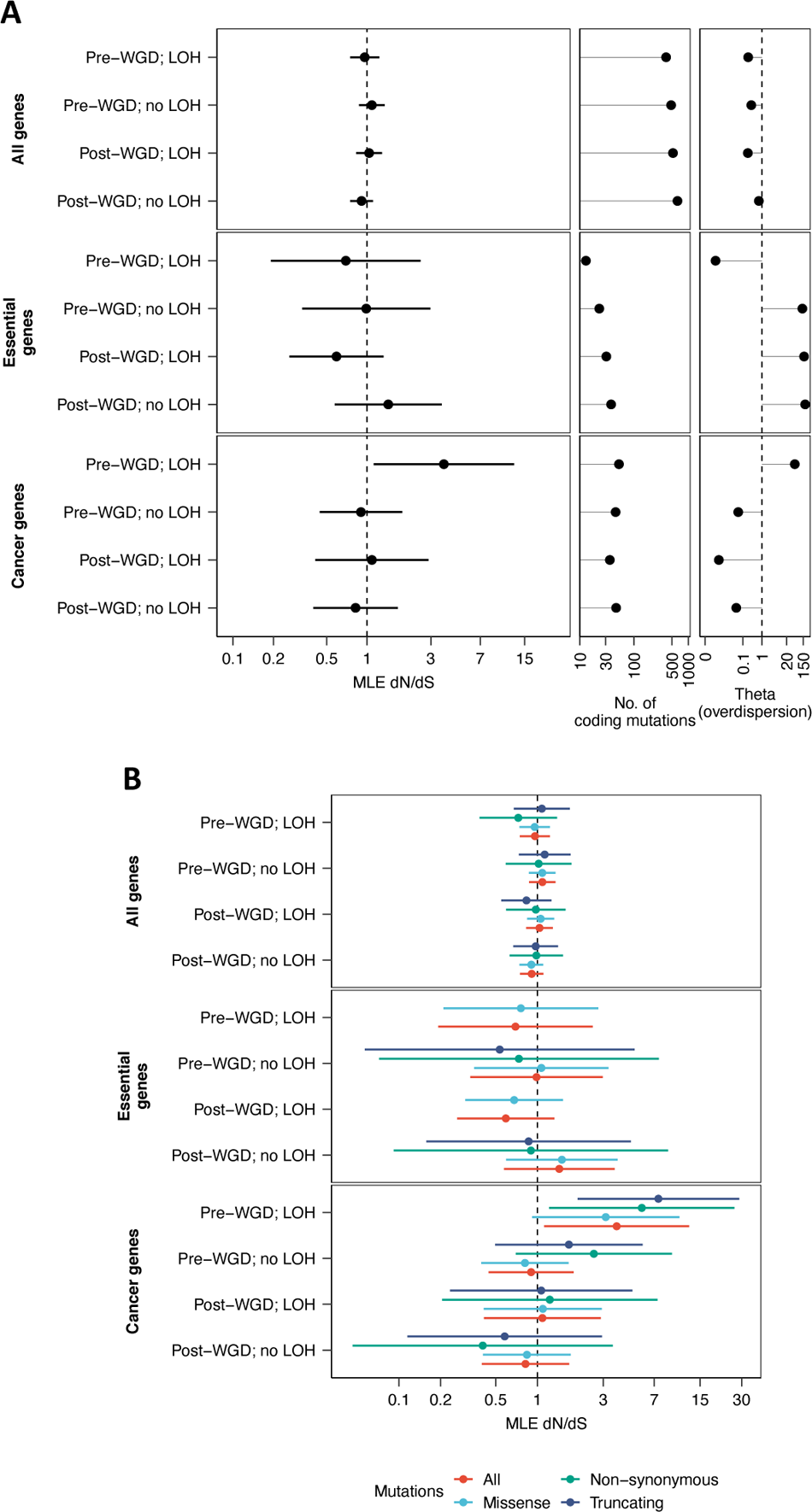
Analysis of selection for coding mutations. **(A)** dN/dS values for coding mutations in high-grade osteosarcomas with WGD calculated for all genes, essential genes, and cancer-associated genes. SNVs were stratified based on their occurrence before or after WGD and their localization to regions showing LOH. dN/dS values equal to, greater and lower than 1 are consistent with neutral evolution, positive selection, and negative selection, respectively. Point dN/dS estimates with 95% confidence intervals are shown. **(B)** Same as (**A**) with results stratified based on the functional impact of somatic coding mutations. MLE dN/dS: maximum-likelihood dN/dS estimates.

**Fig. S32.**
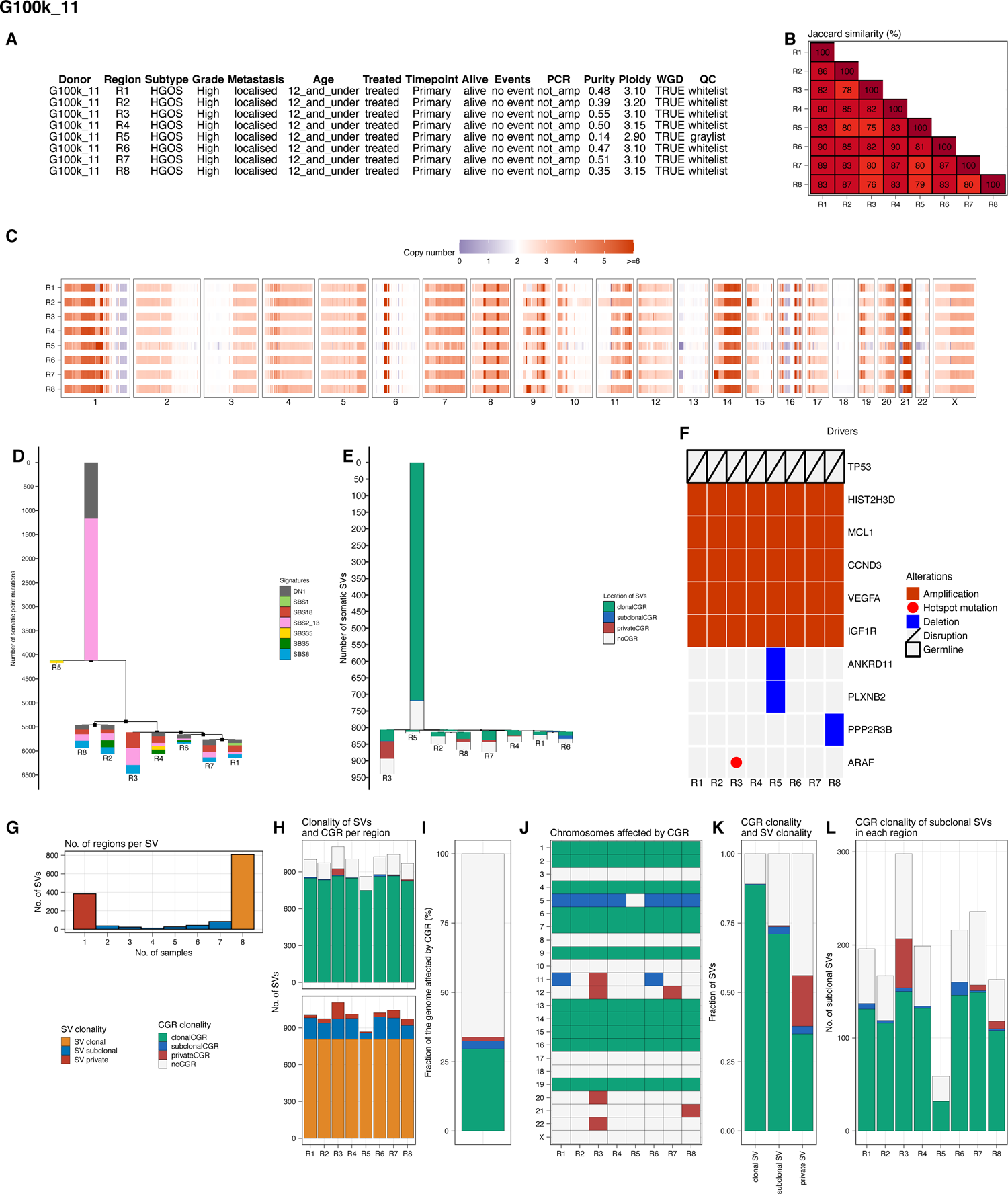
Analysis of multi-region WGS data from osteosarcomas. **(A)** Clinical and genomic data for each tumour region analysed. **(B)** Pairwise Jaccard similarity index computed using vectors recording the presence or absence of the structural variants detected across all regions that were found in each tumour region. **(C)** Absolute copy number profile for each region. **(D)** Phylogenetic tree constructed using somatic single-nucleotide variants. The colours represent the contribution of each SBS mutational signature to the mutations in each branch. **(E)** Phylogenetic trees constructed using somatic SVs detected in multi-region WGS data. The colours indicate whether the SVs in the tree map to clonal, subclonal or private CGRs (green, blue, and red), respectively, or outside CGR regions (grey). **(F)** Somatic aberrations detected in driver genes across tumour regions. **(G)** Number of SVs detected in the number of tumour regions indicated in the x axis. **(H)** Top: Distribution of the number of SVs per region stratified based on whether they map to clonal (green), subclonal (blue) or private (red) CGRs. The number of SVs mapping outside CGRs is indicated in grey. Bottom: Distribution of the number of SVs per region stratified based on clonality. **(I)** Percentage of the tumour genome affected by clonal (green), subclonal (blue) and private (red) CGRs. The fraction of the tumour genome without CGRs is shown in grey. **(J)** Distribution of clonal, subclonal and private CGRs across chromosomes and tumour regions. **(K)** Fraction of clonal, subclonal and private SVs mapping to clonal (green), subclonal (blue) and private (red) CGRs. SVs mapping outside CGRs are shown in grey. **(L)** Number of subclonal SVs in each tumour region stratified based on whether they map to clonal (green), subclonal (blue) and private (red) CGRs. The number of SVs mapping outside CGRs is shown in grey.

**Fig. S33.**
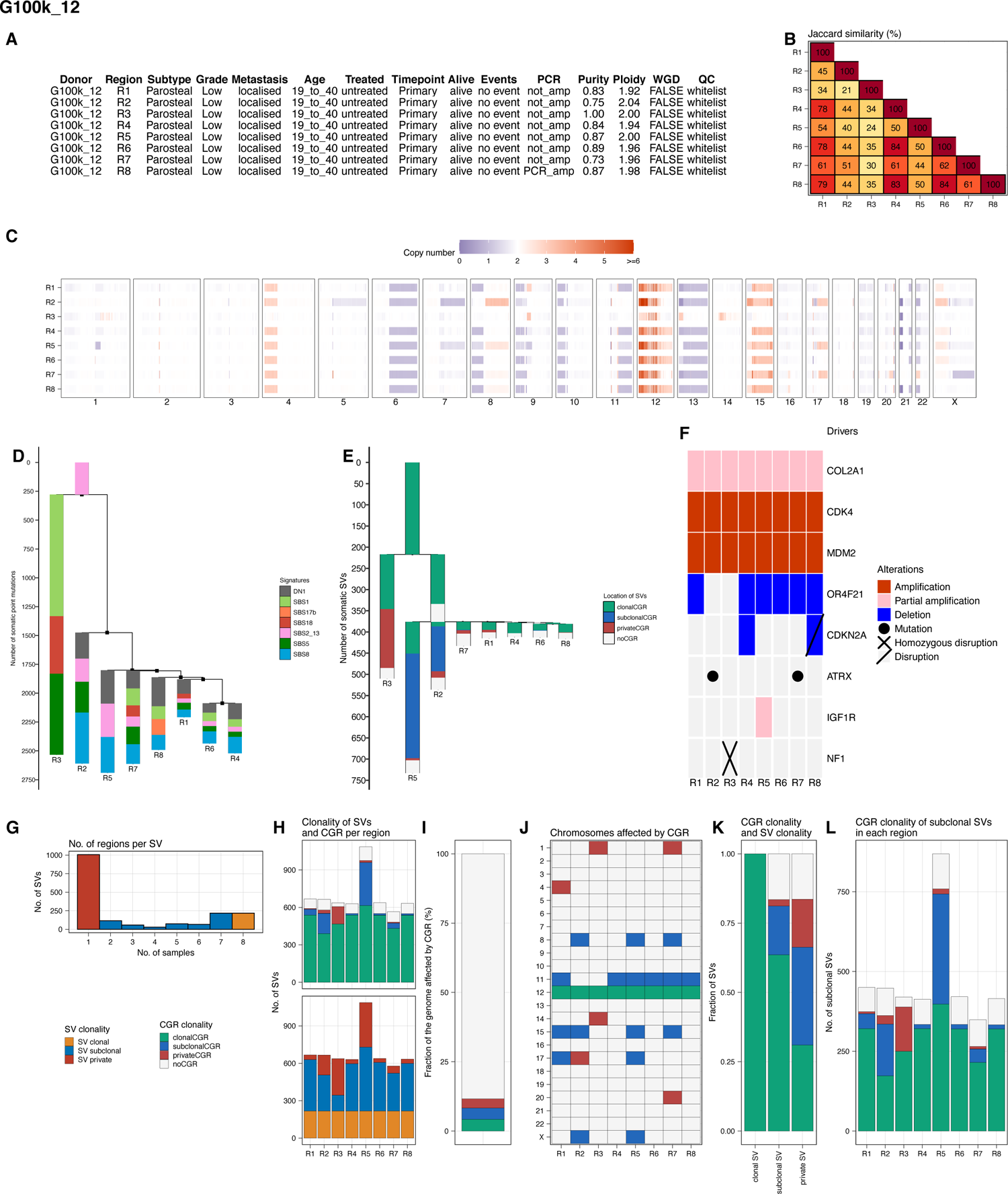

**Fig. S34.**
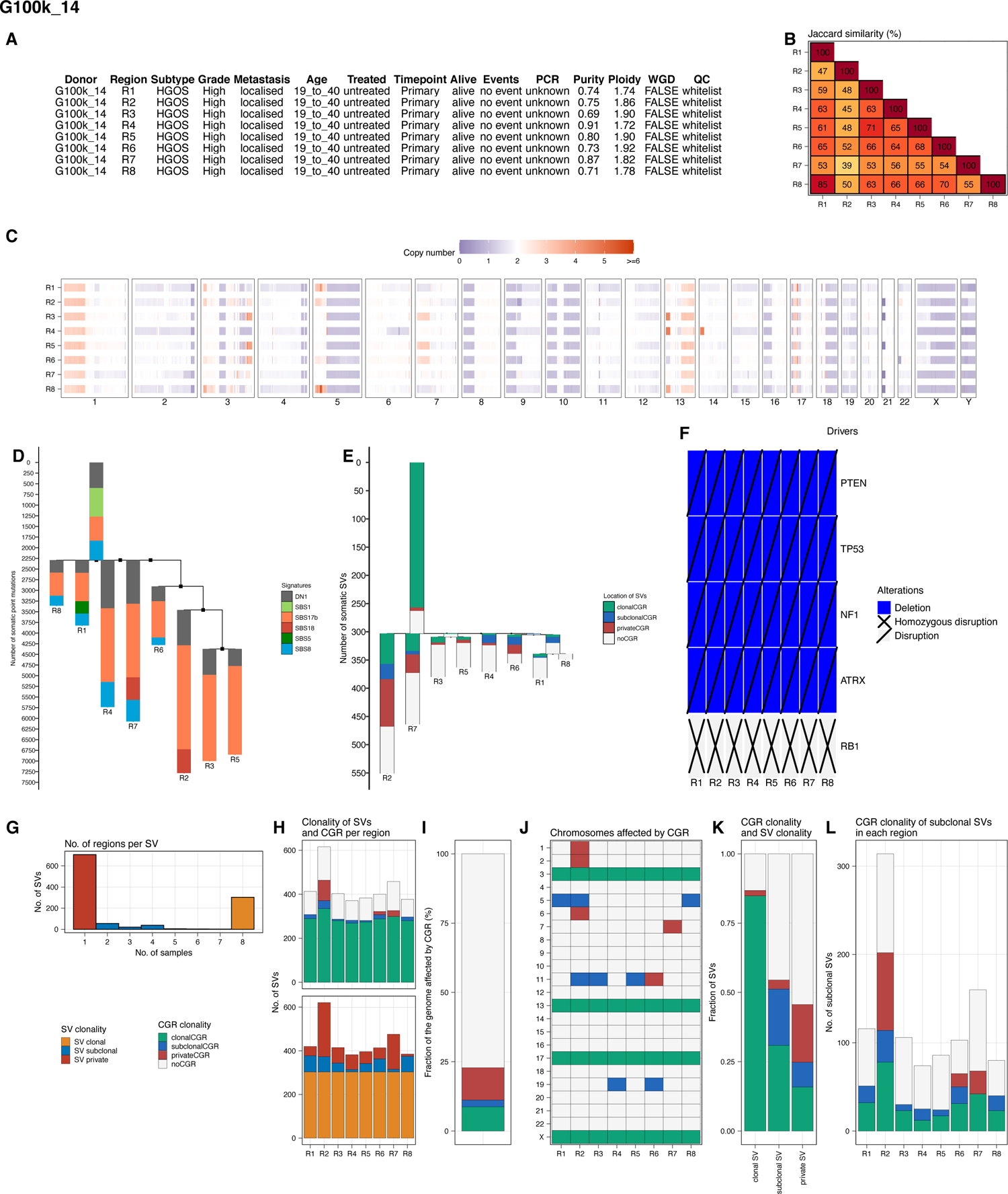

**Fig. S35.**
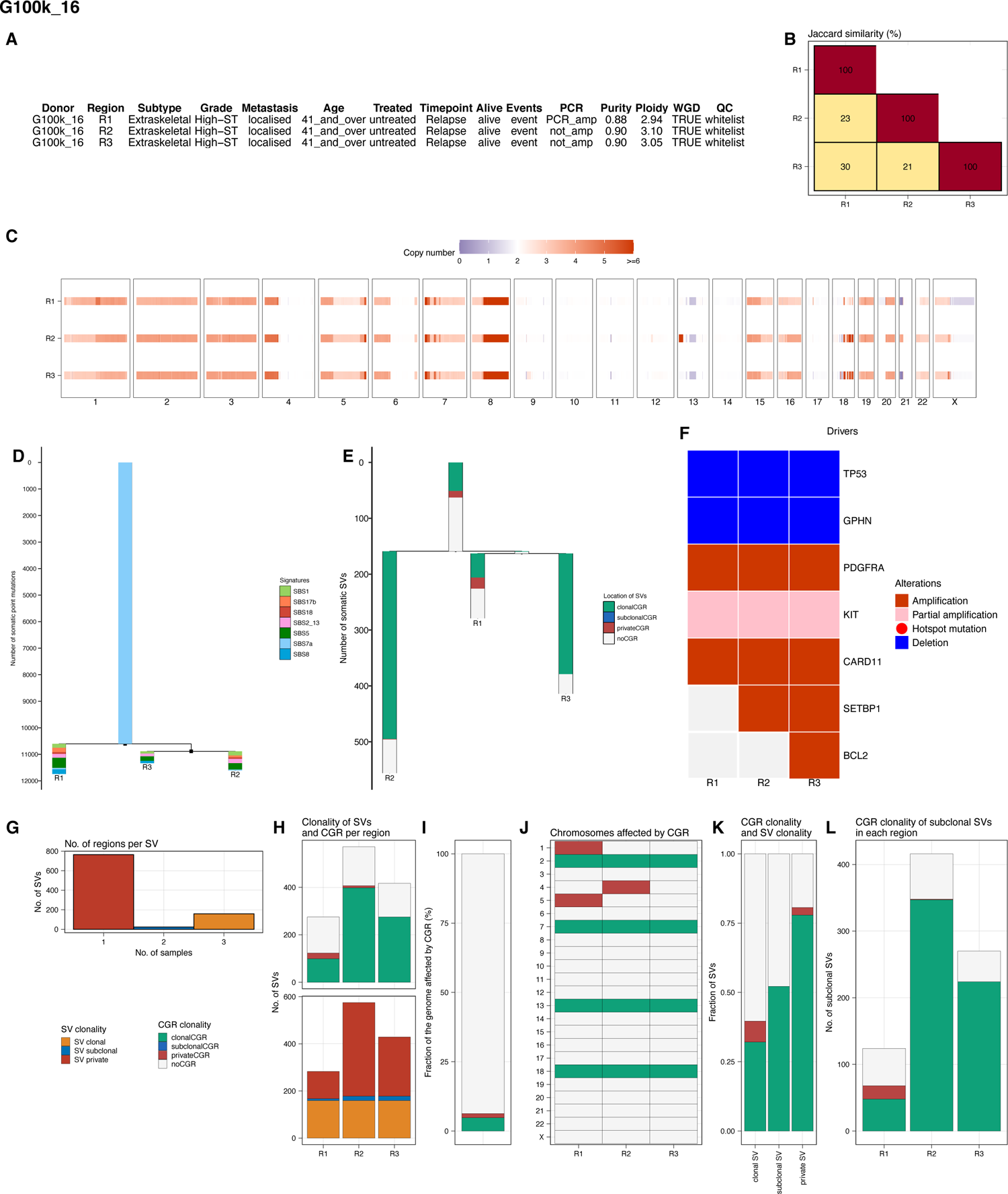

**Fig. S36.**
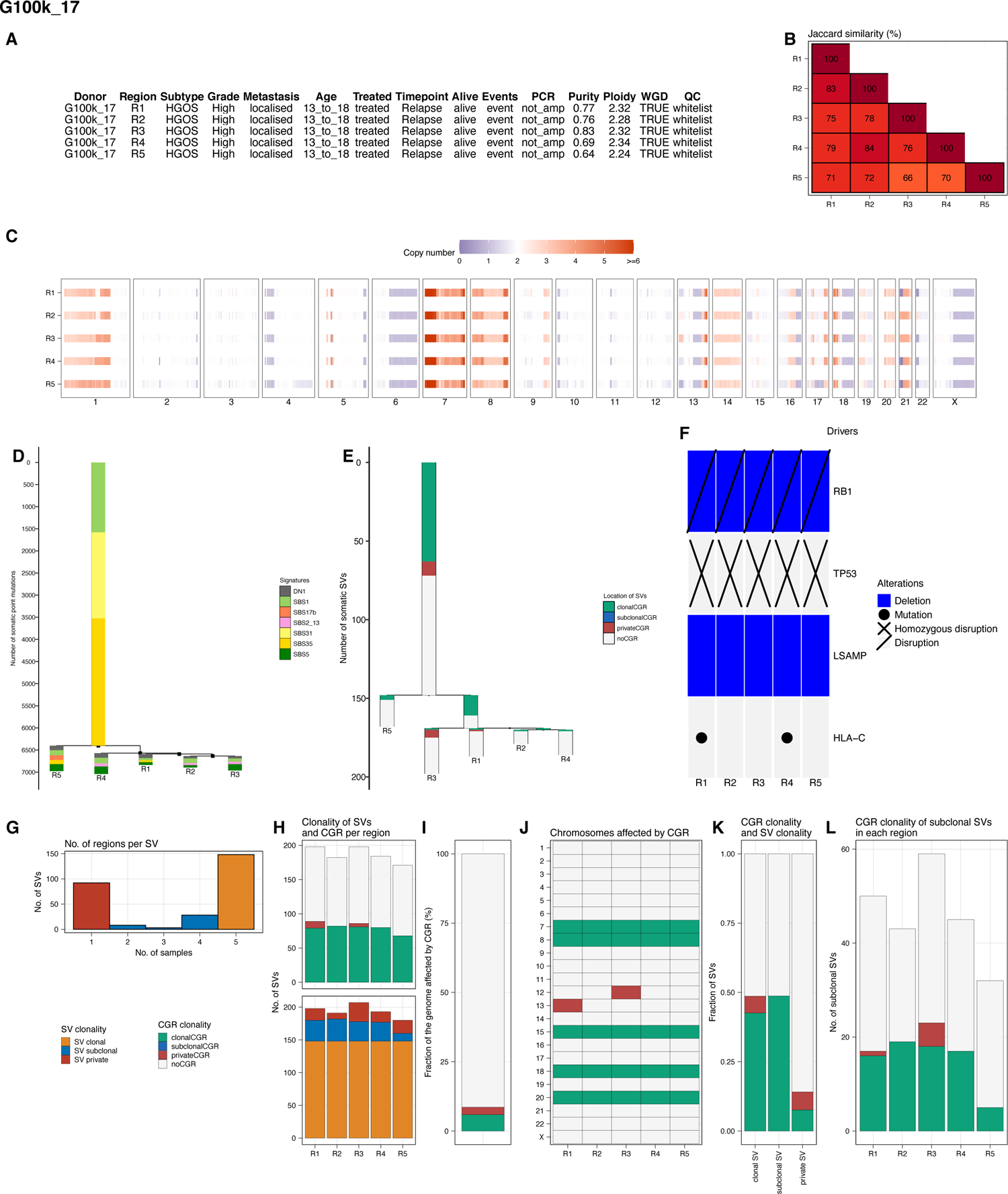

**Fig. S37.**
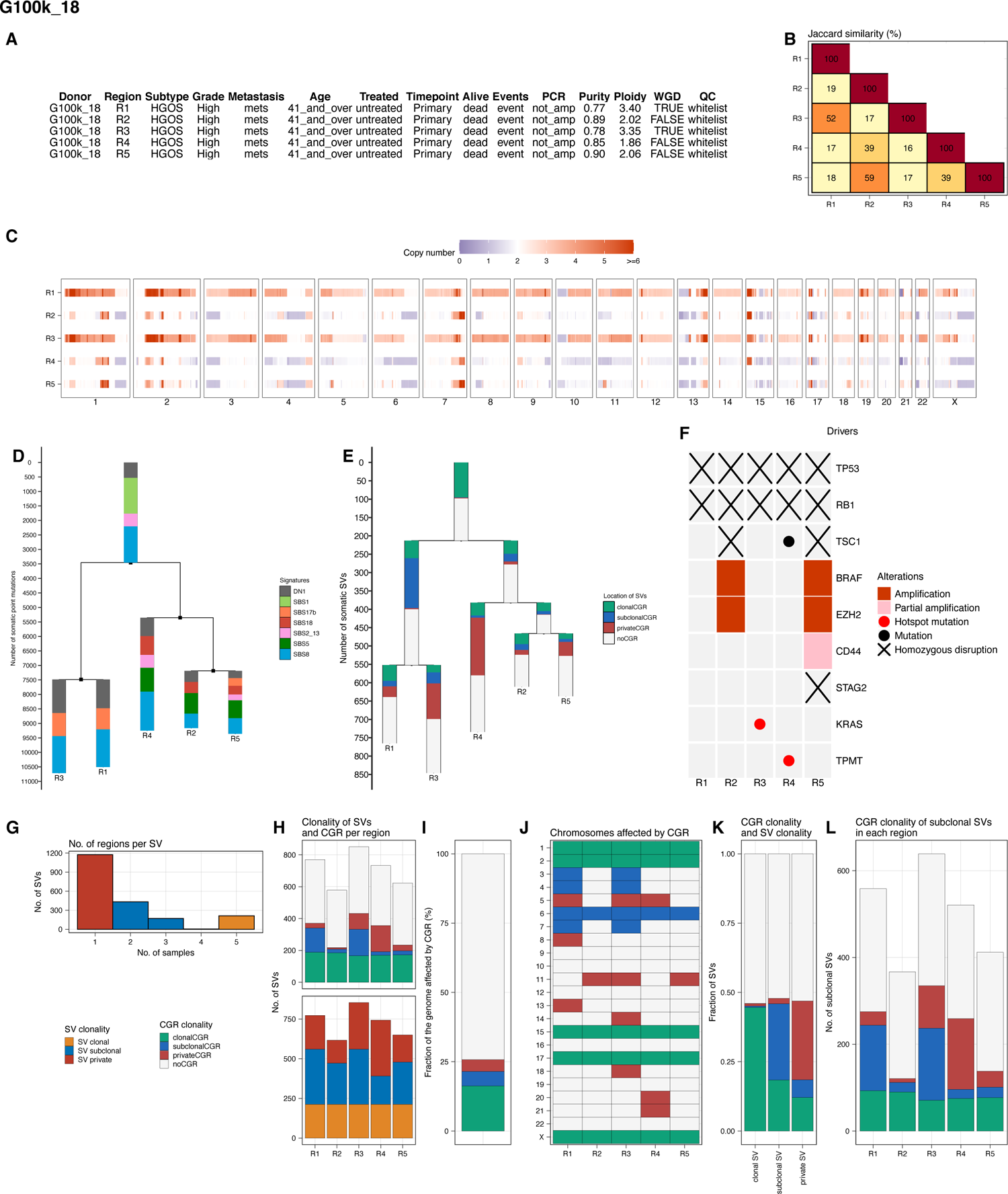

**Fig. S38.**
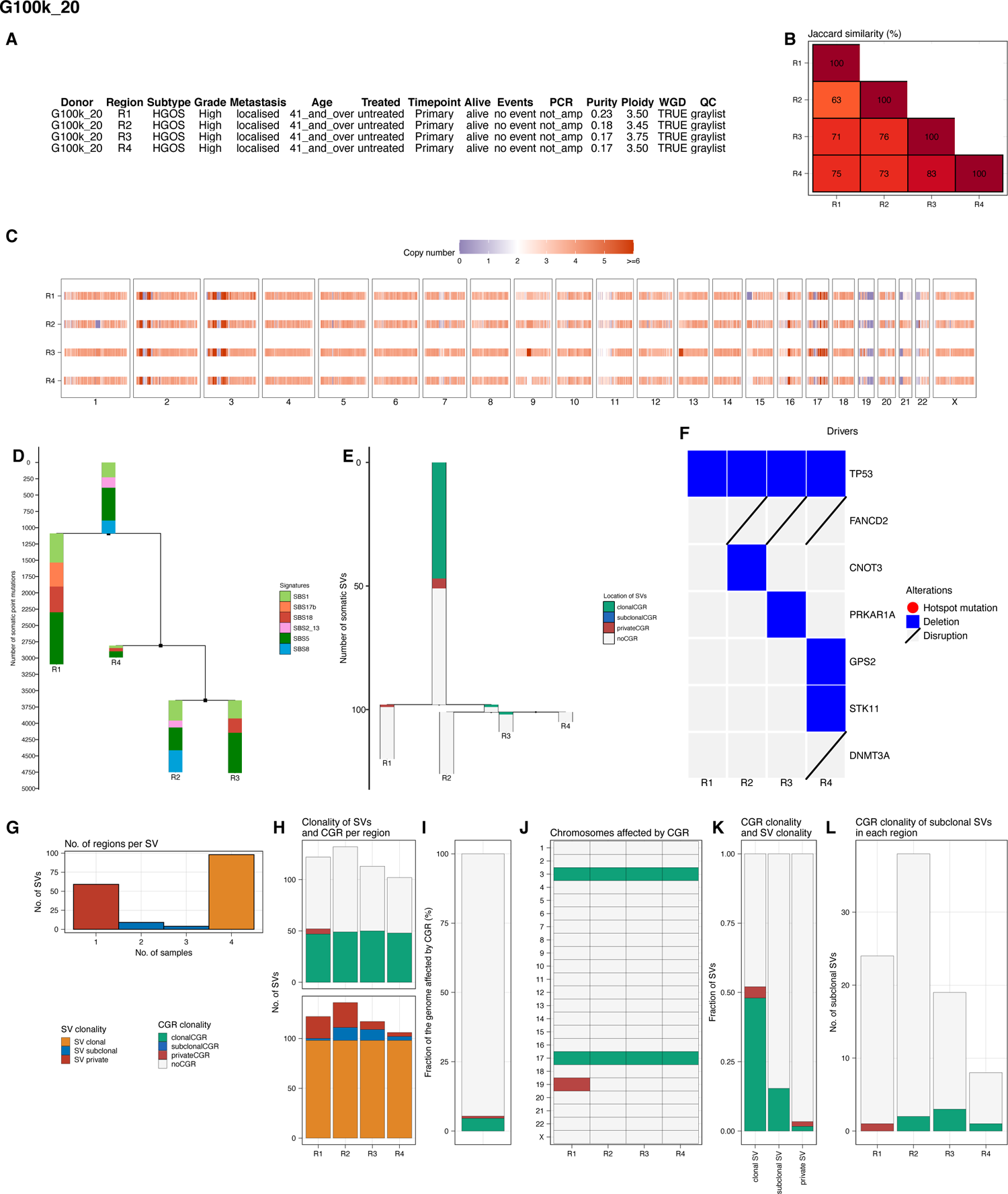

**Fig. S39.**
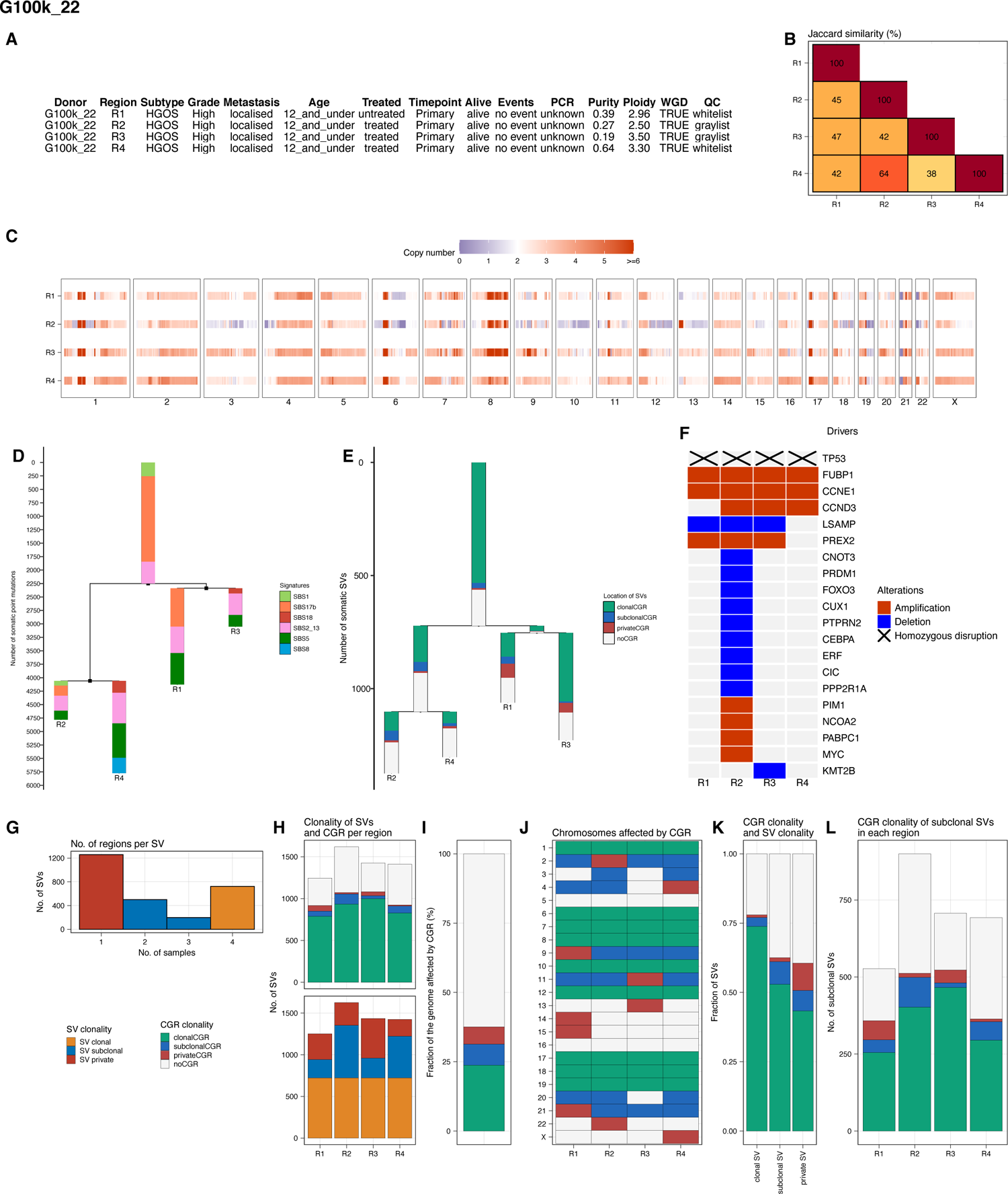

**Fig. S40.**
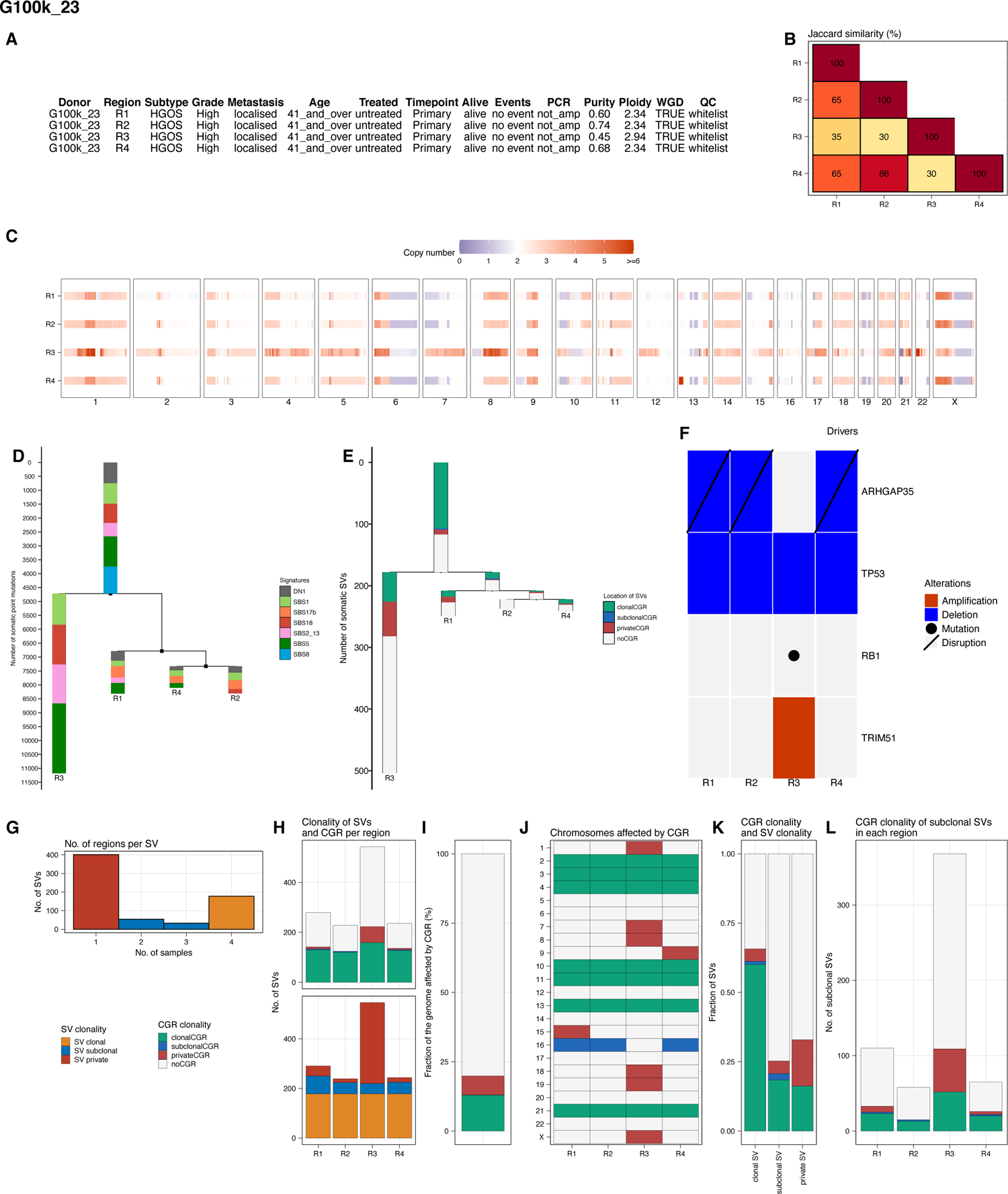

**Fig. S41.**
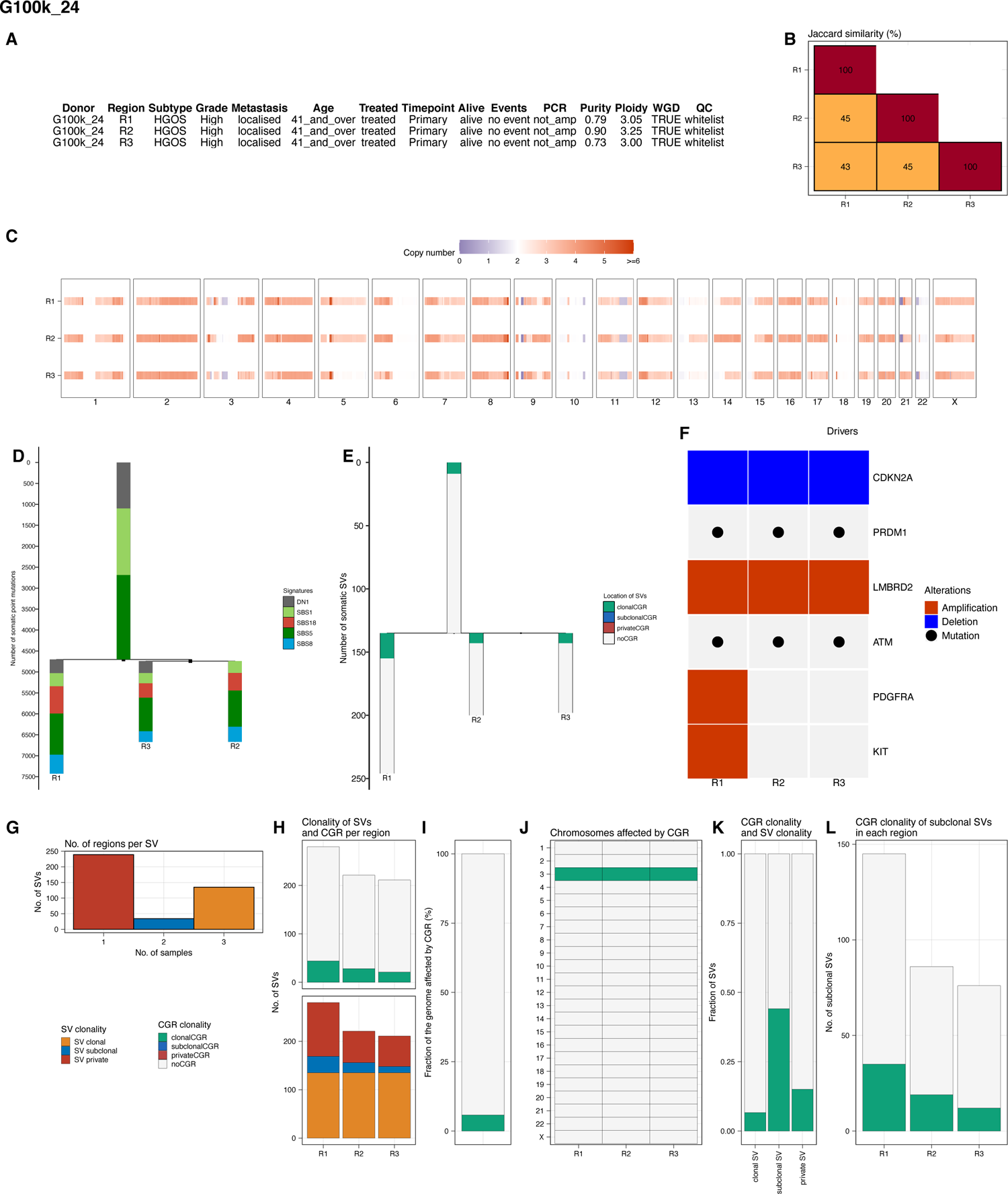

**Fig. S42.**
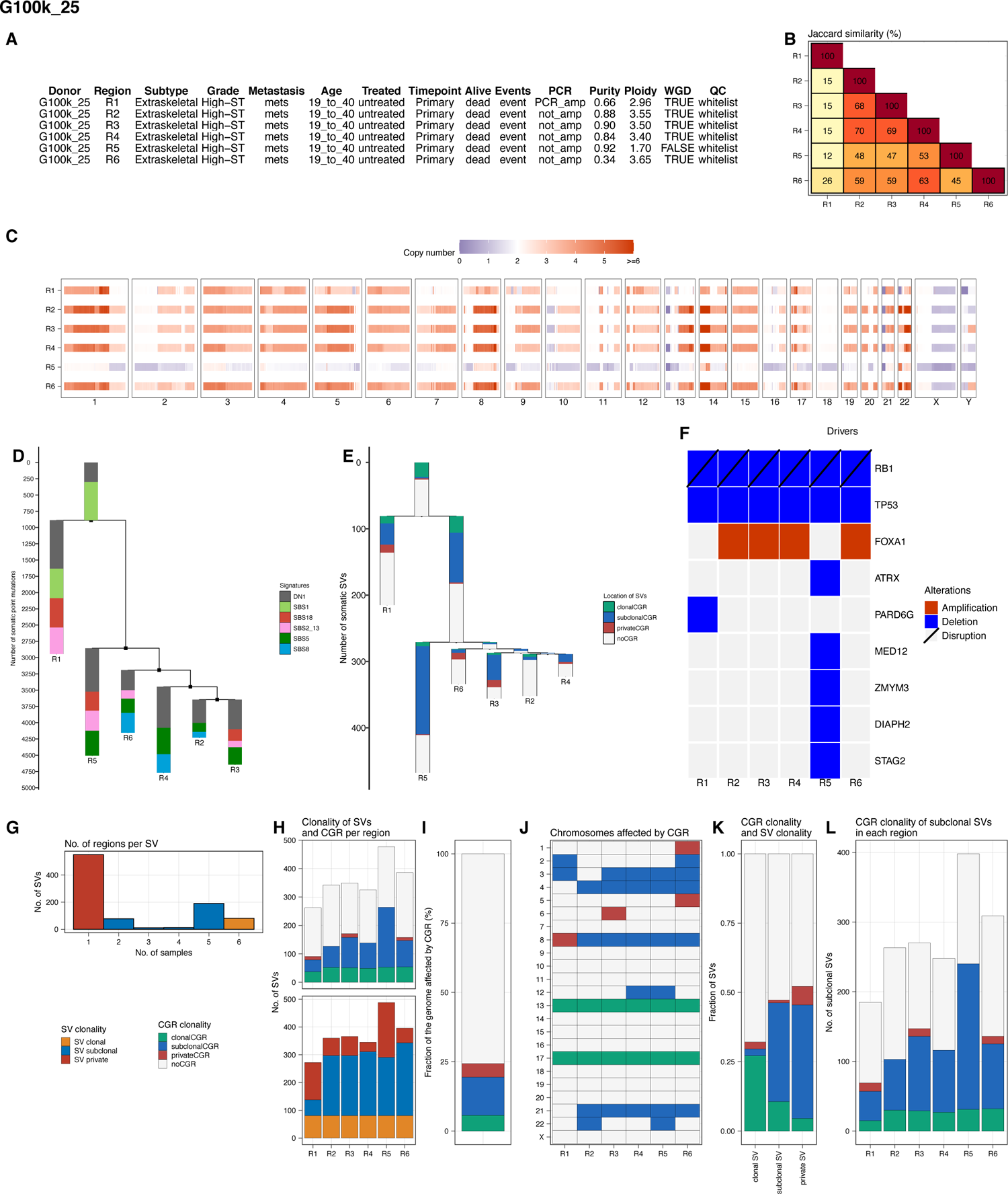

**Fig. S43.**
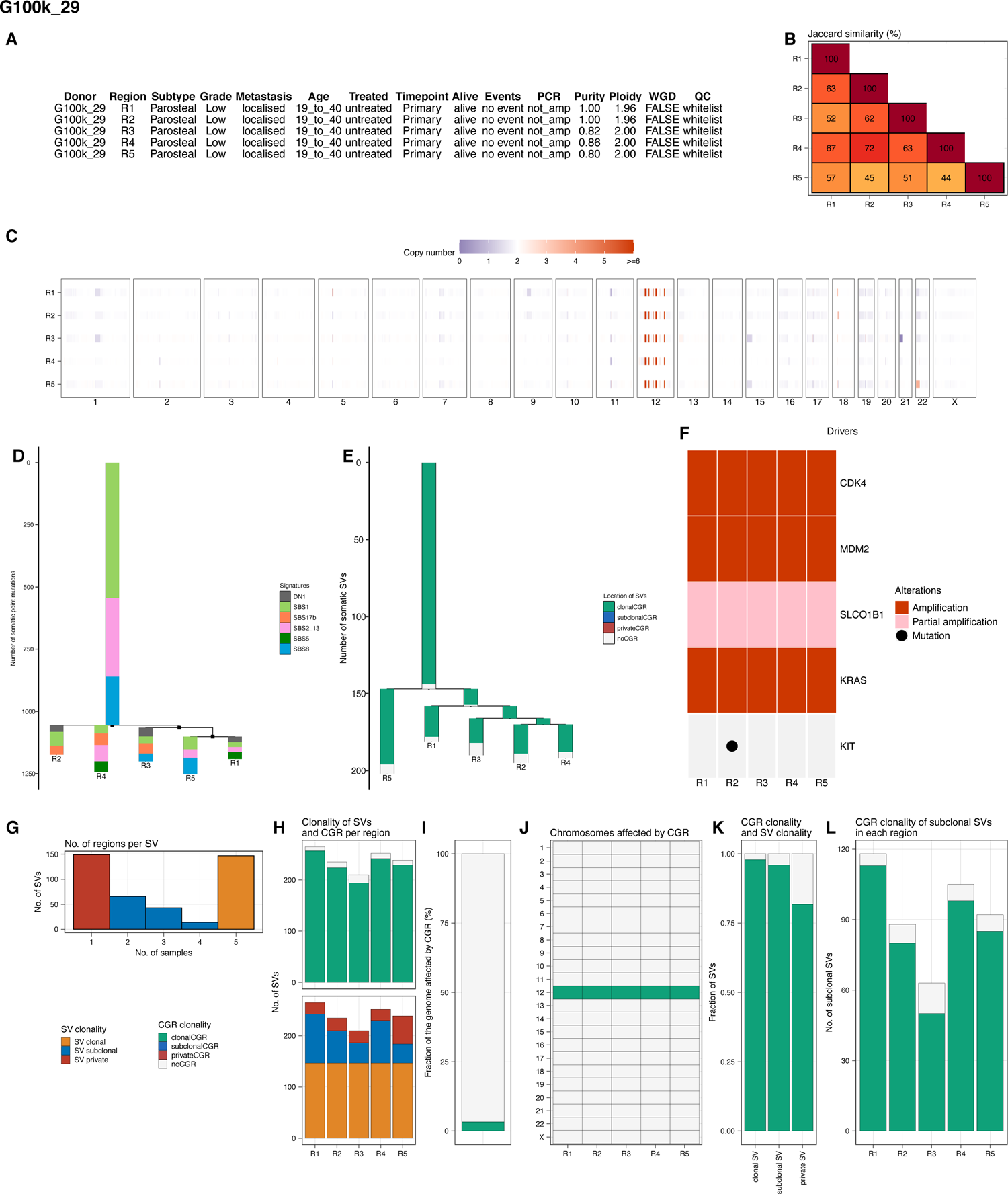

**Fig. S44.**
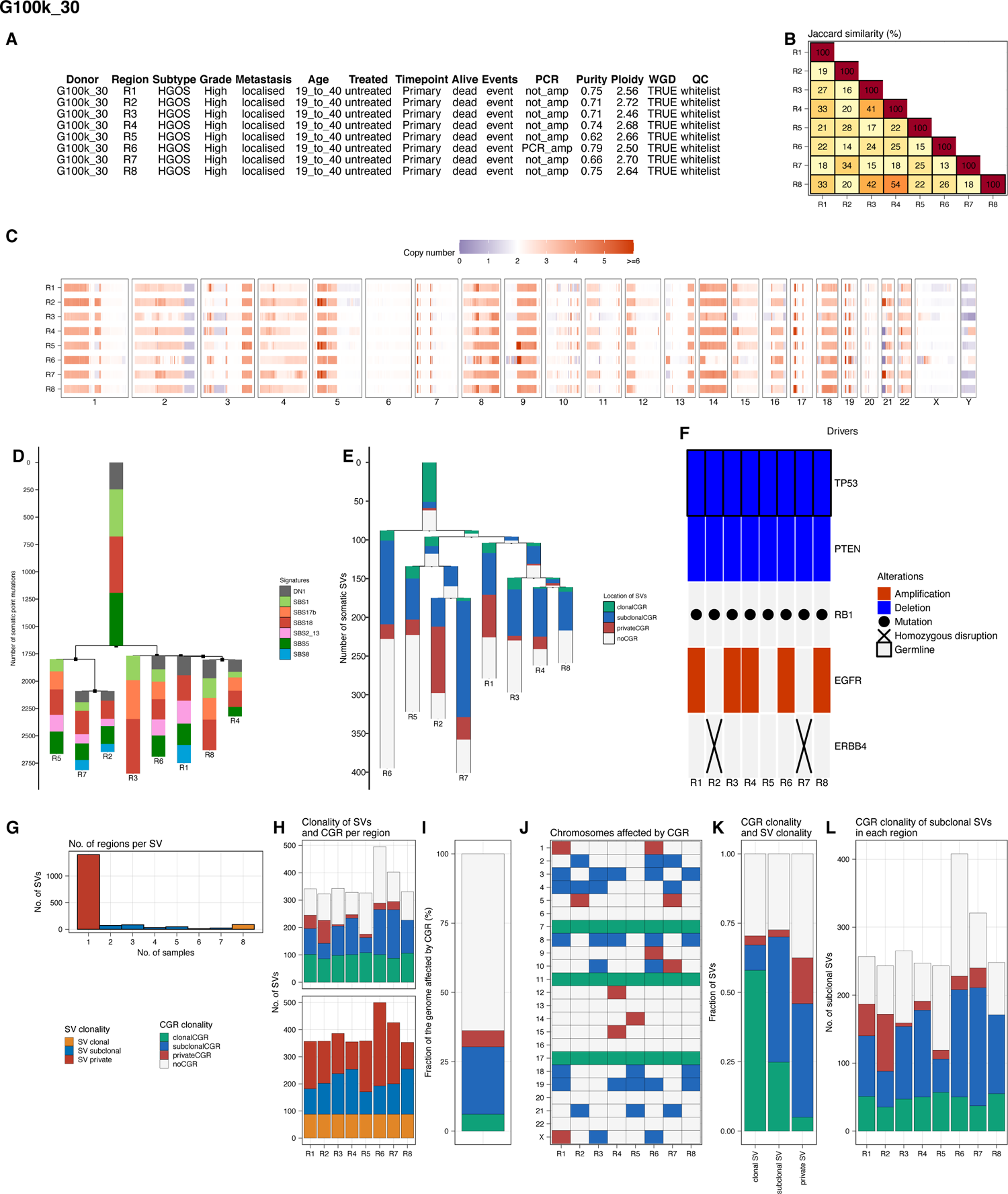

**Fig. S45.**
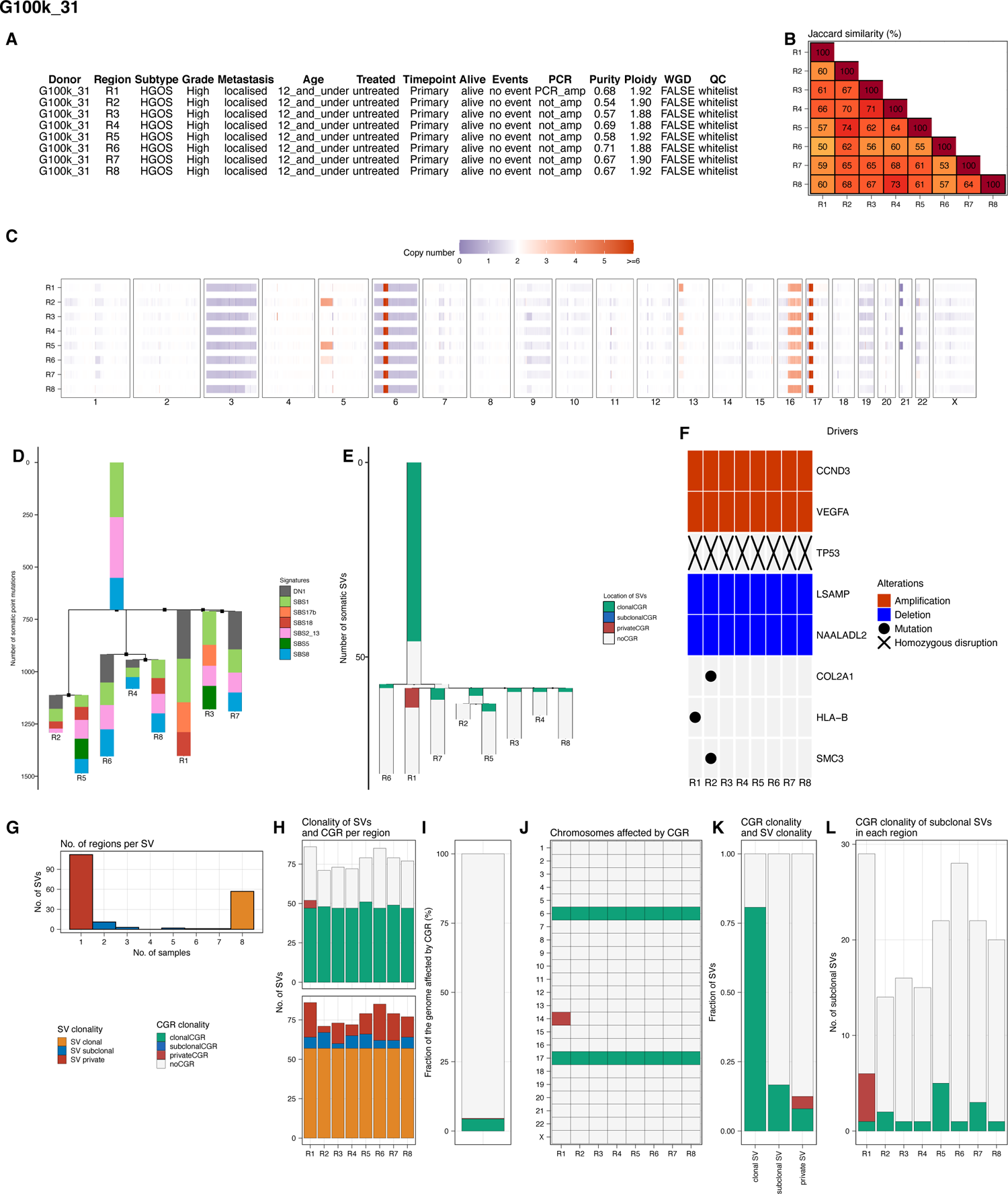

**Fig. S46.**
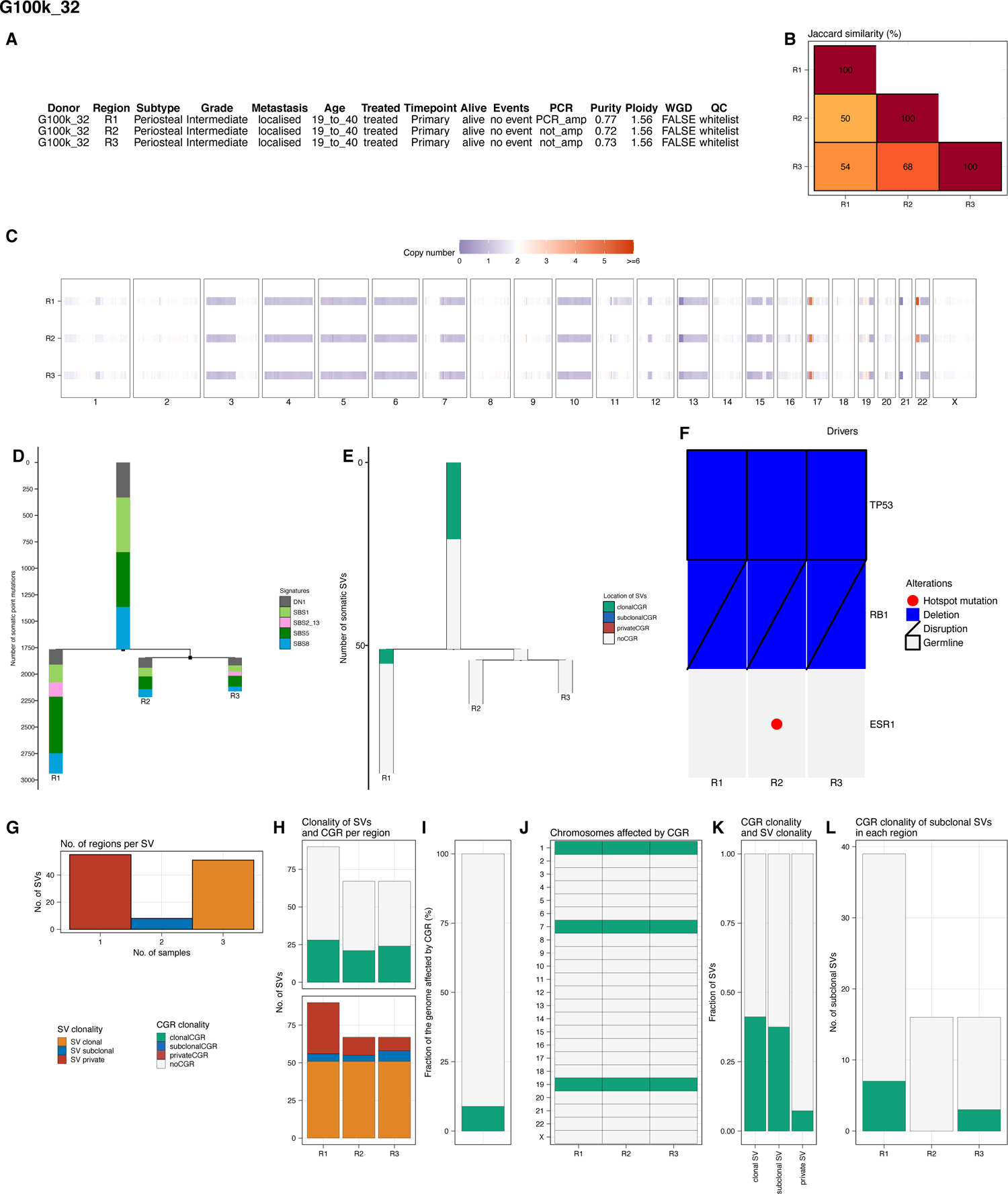

**Fig. S47.**
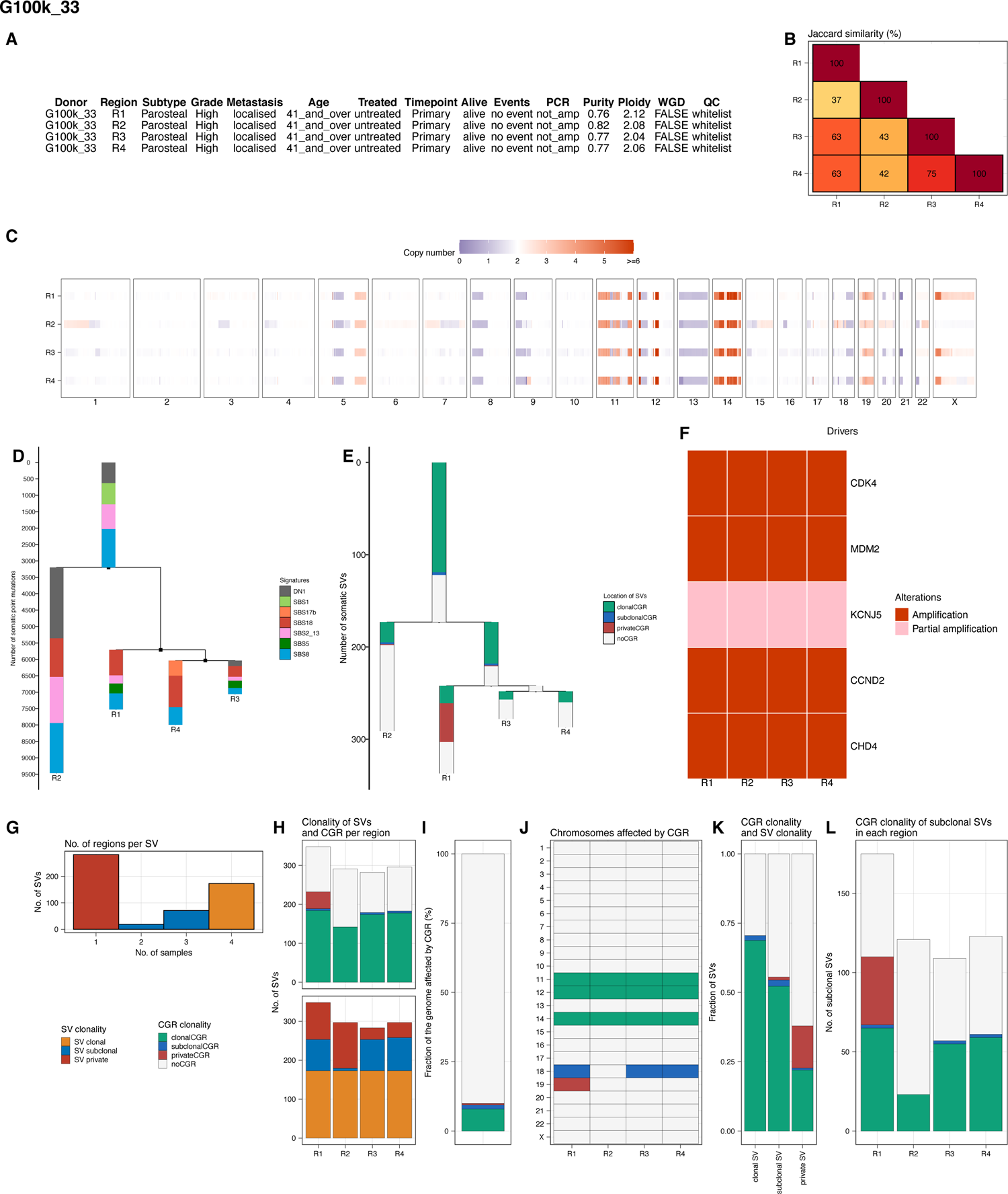

**Fig. S48.**
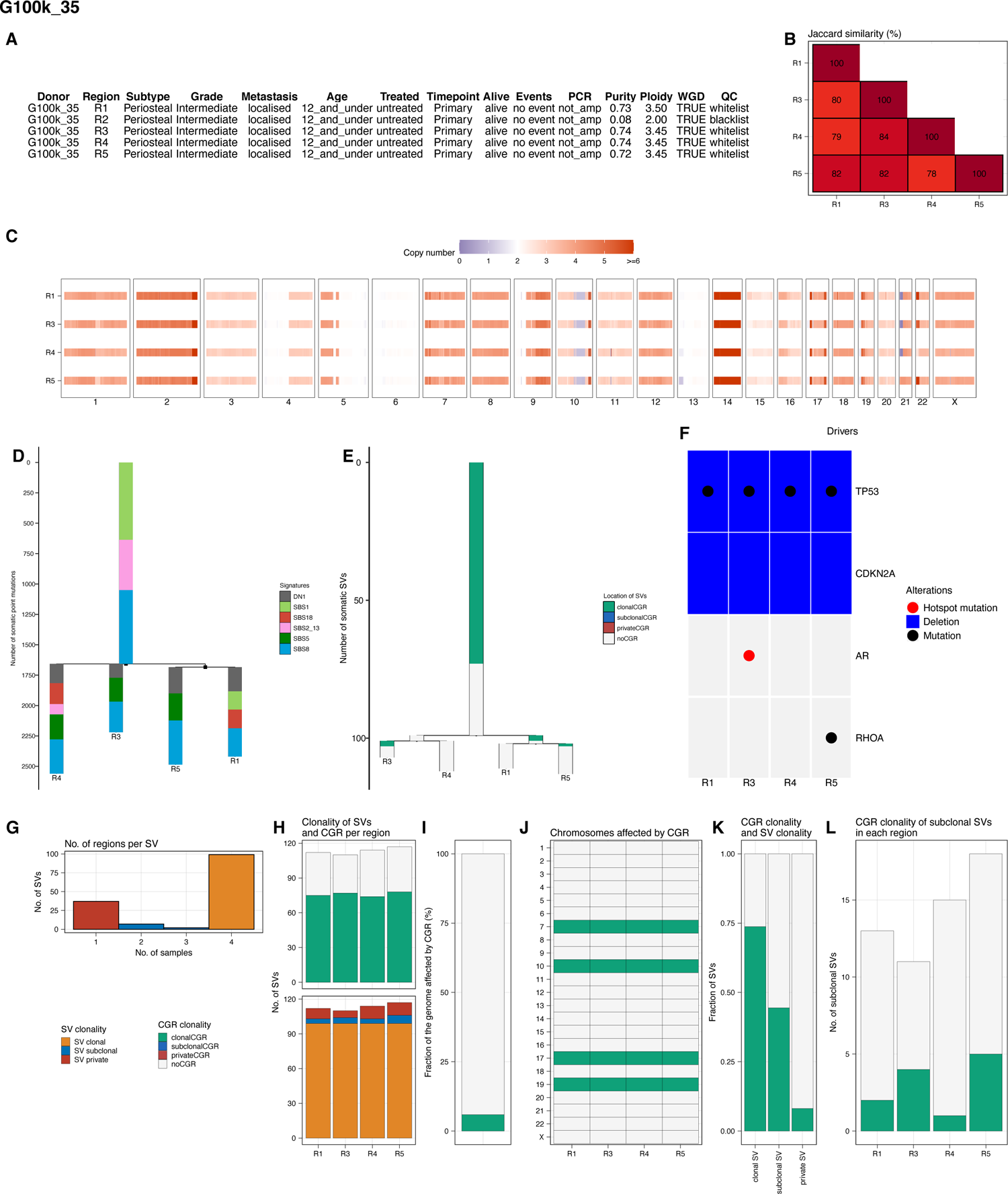

**Fig. S49.**
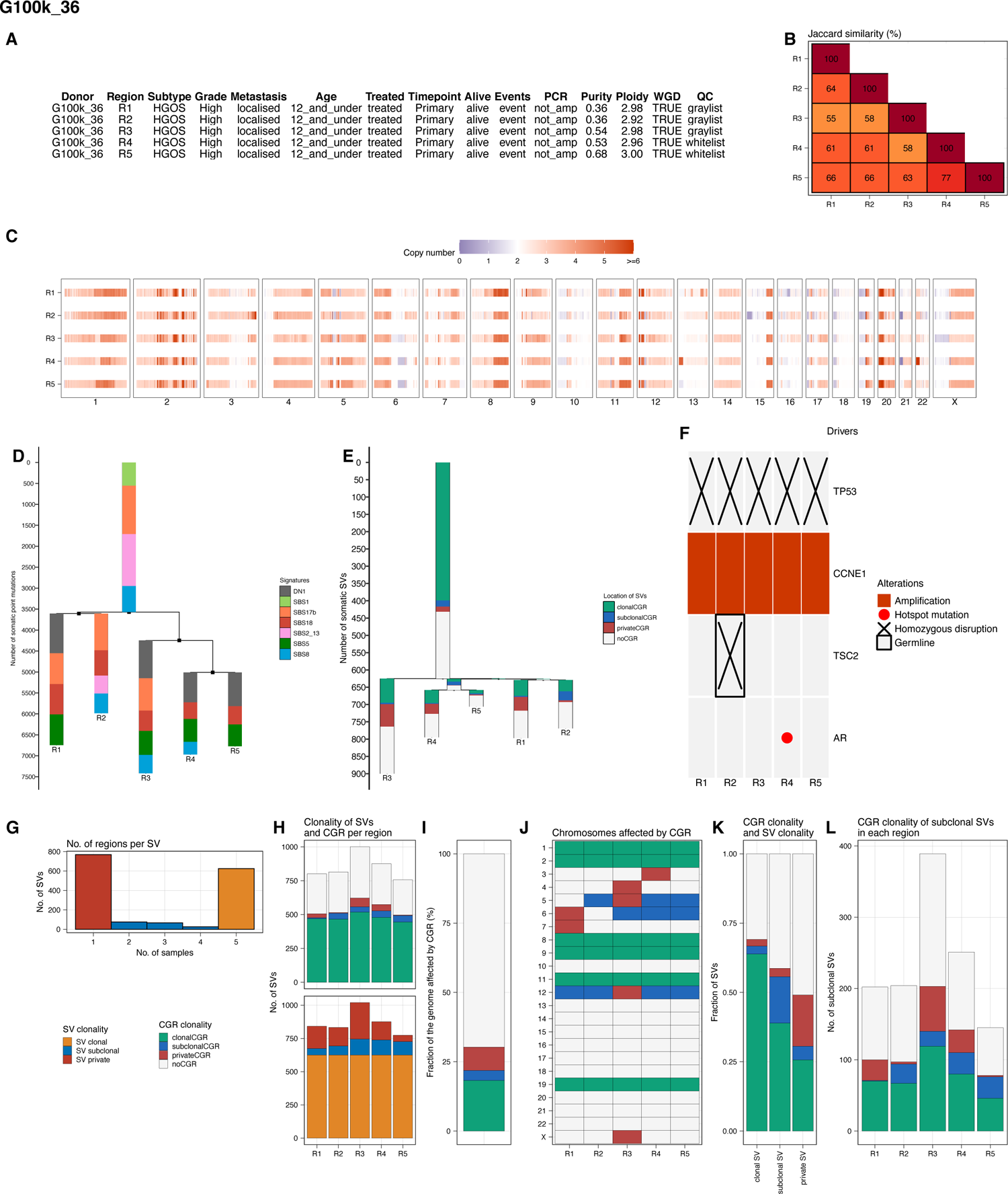

**Fig. S50.**
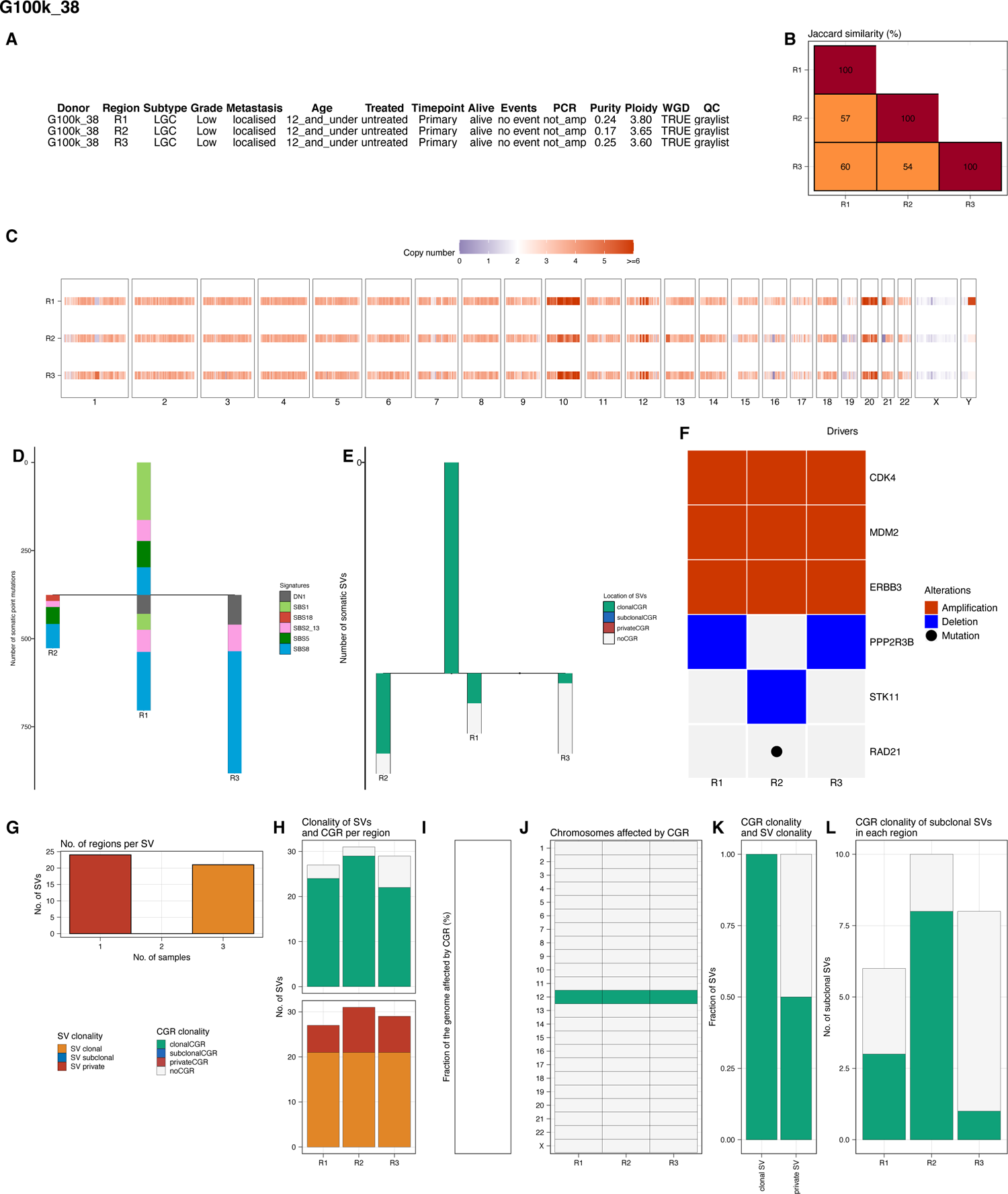

**Fig. S51.**
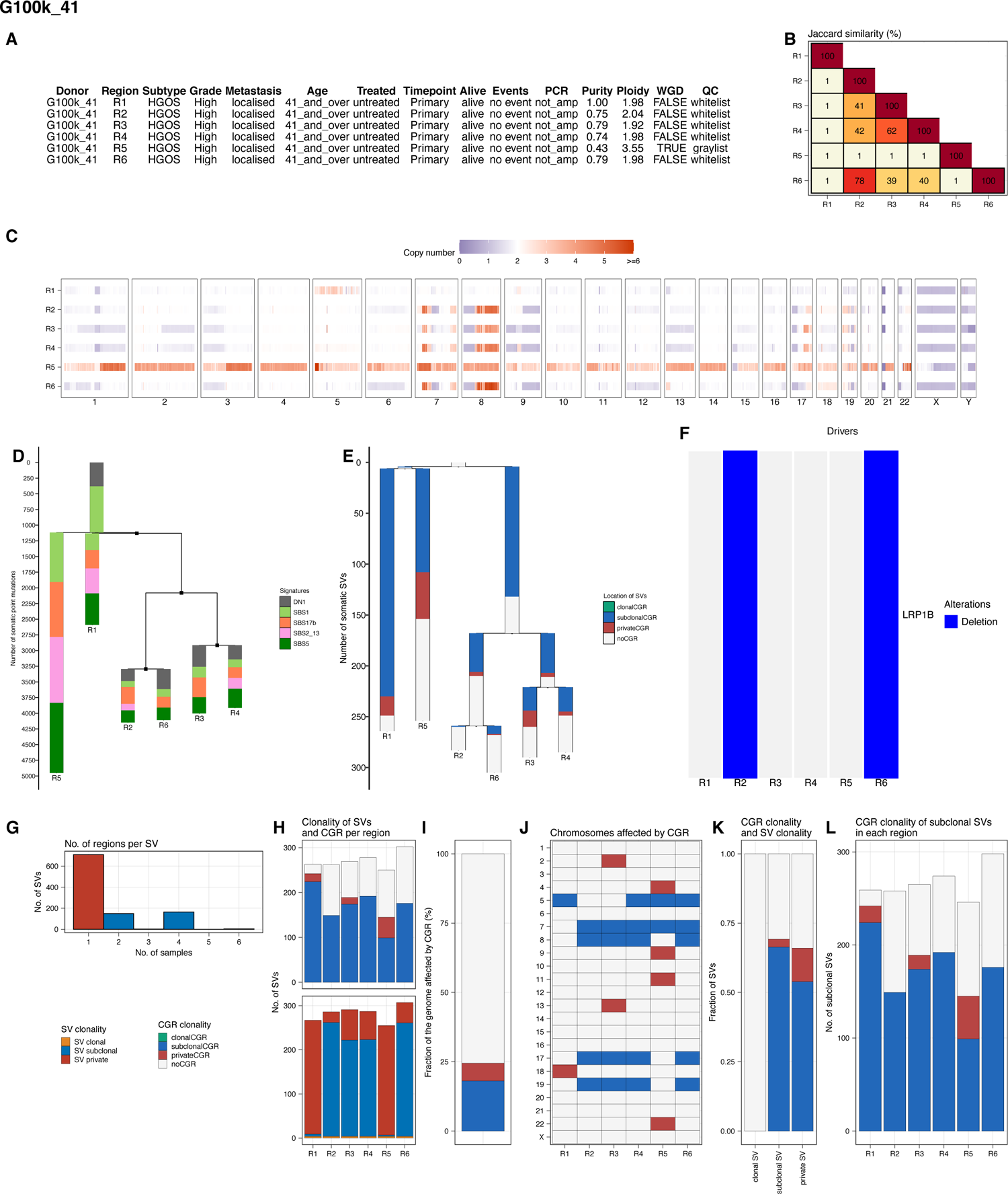

**Fig. S52.**
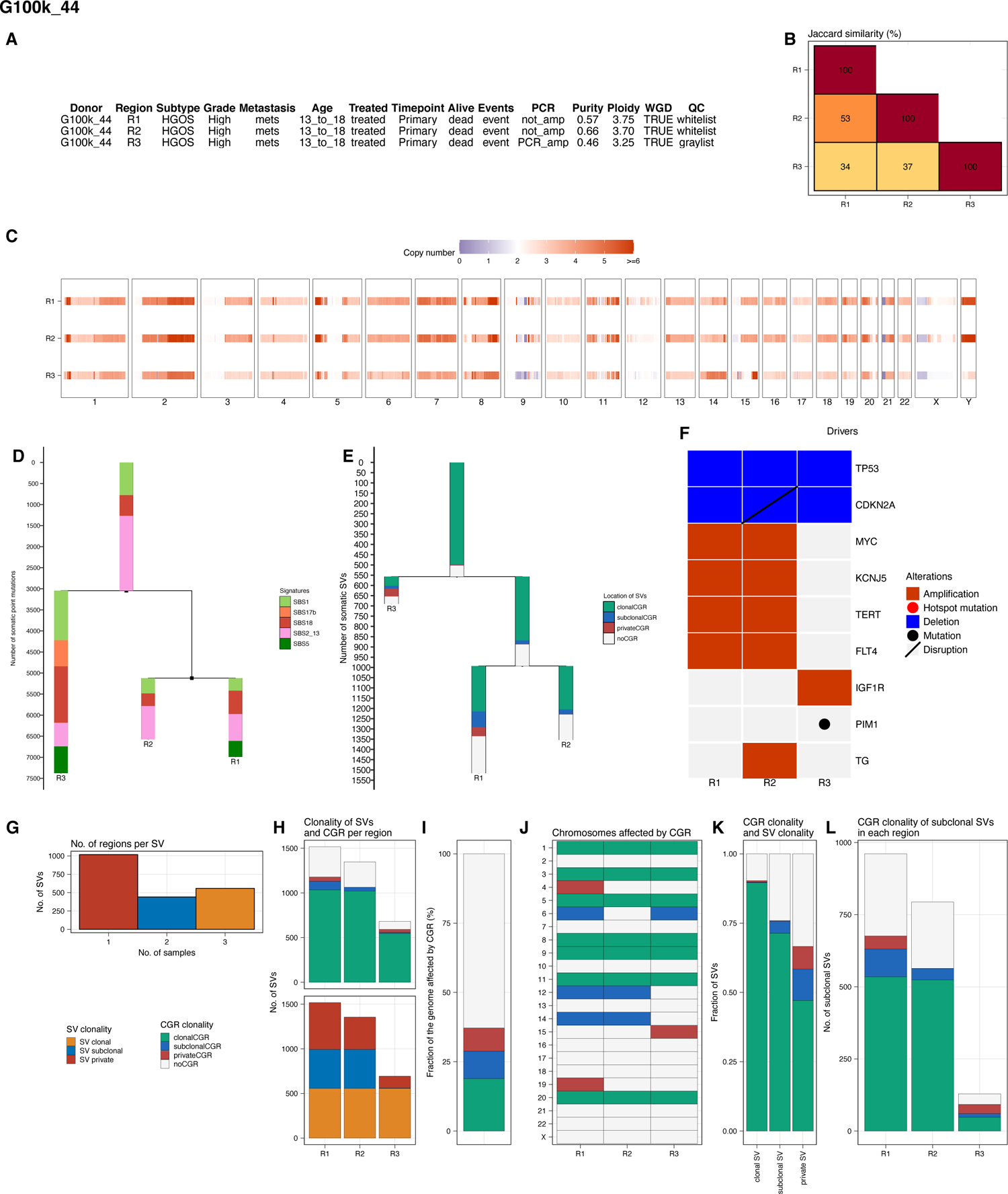

**Fig. S53.**
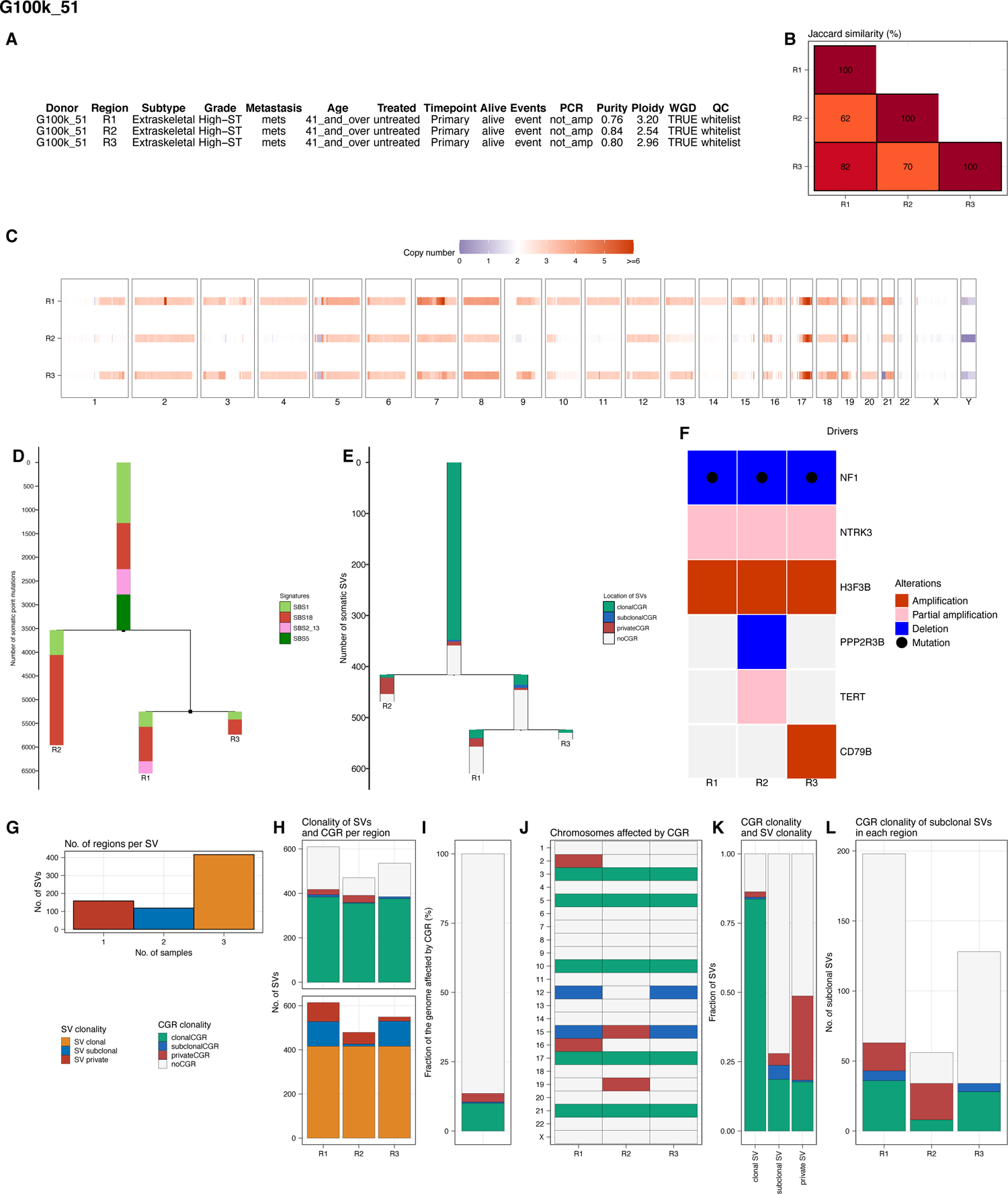

**Fig. S54.**
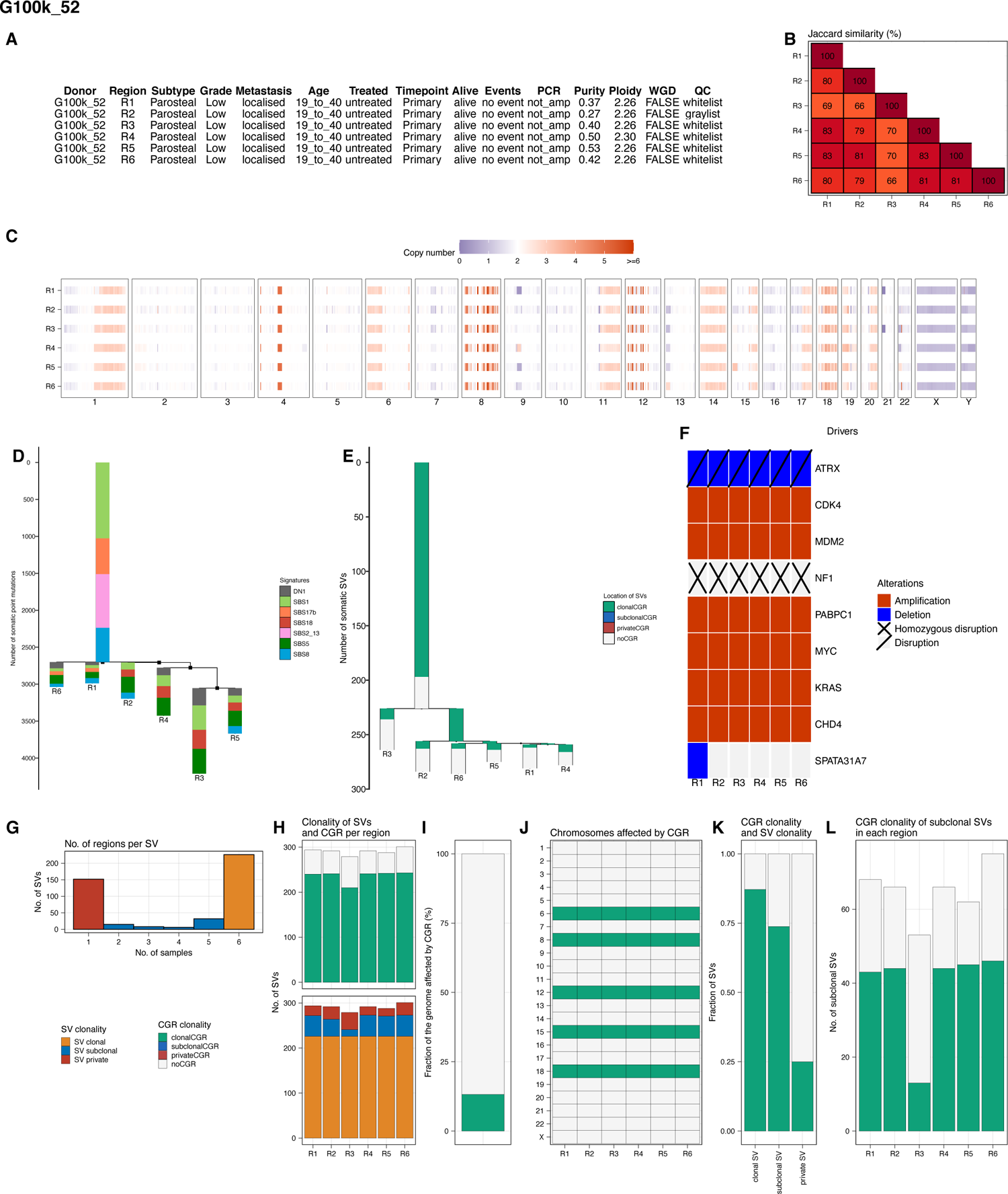

**Fig. S55.**
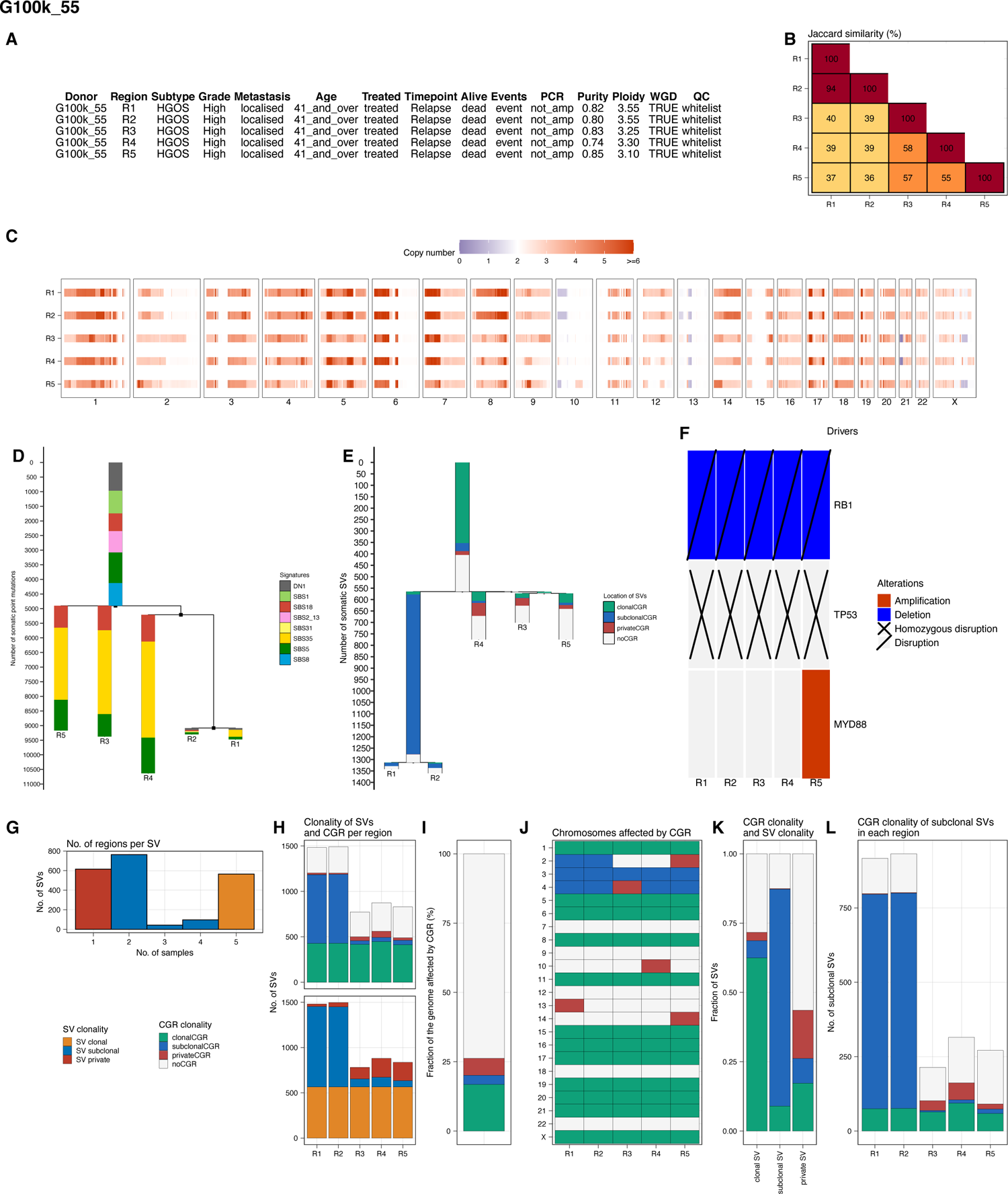

**Fig. S56.**
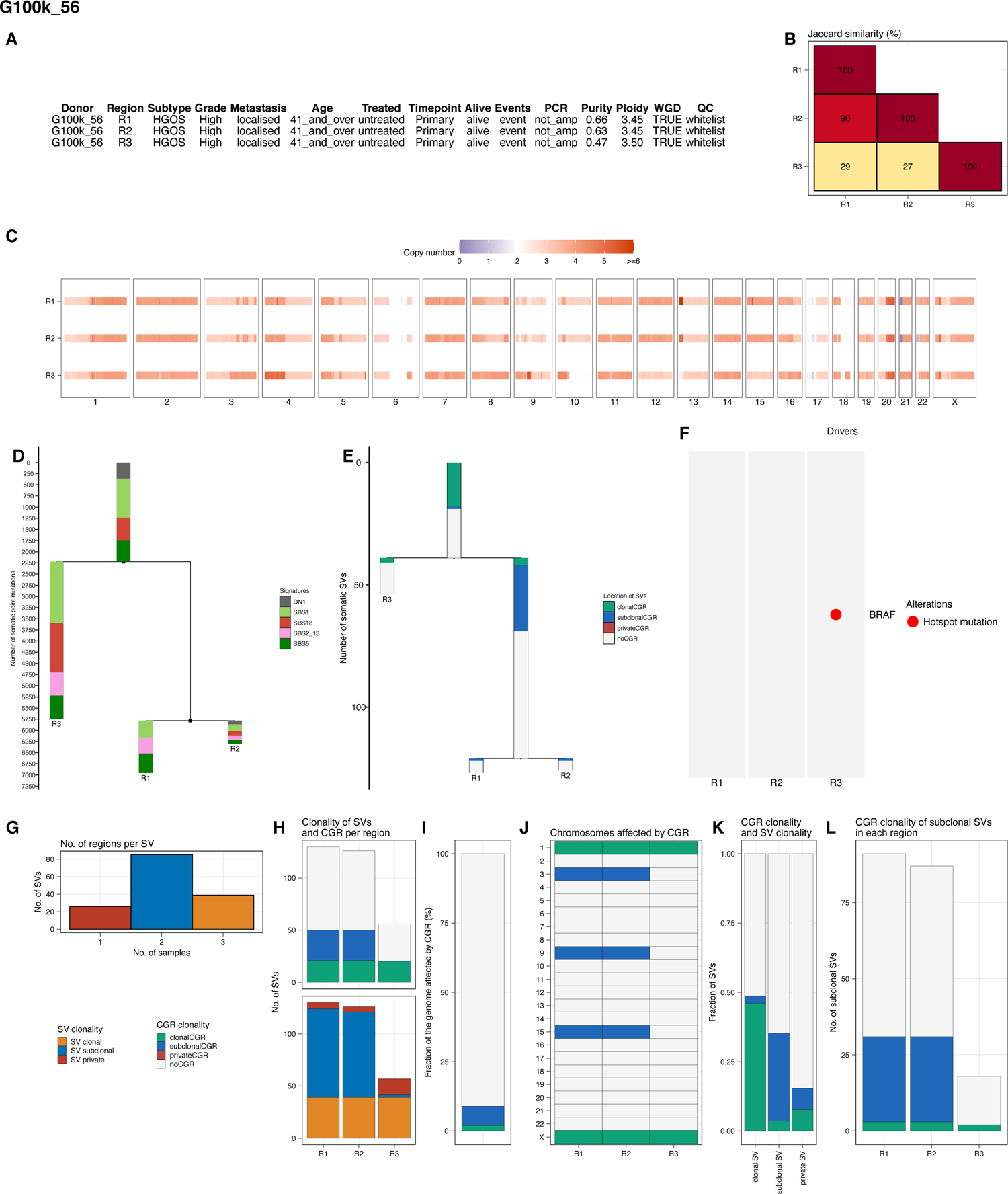

**Fig. S57.**
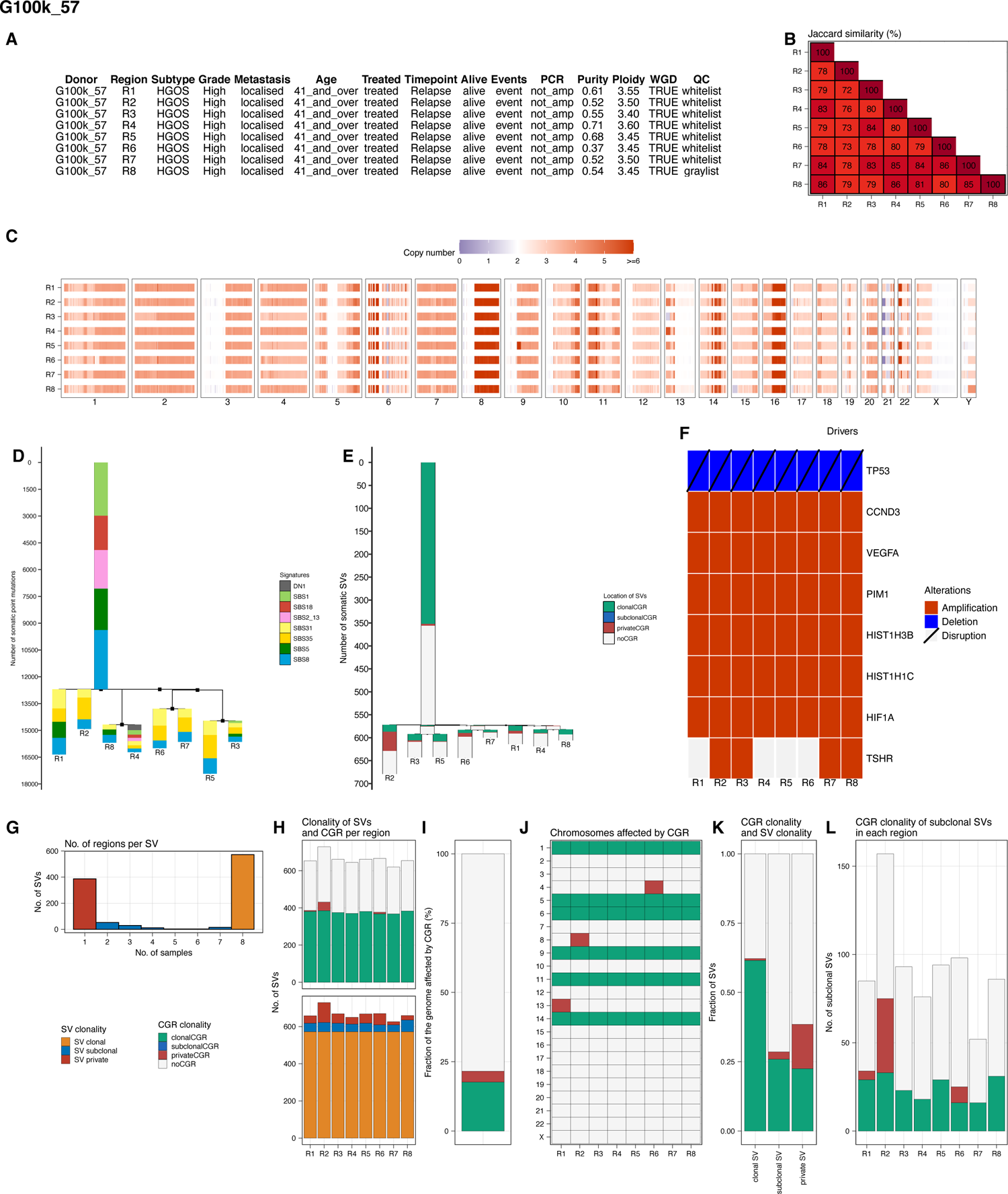

**Fig. S58.**
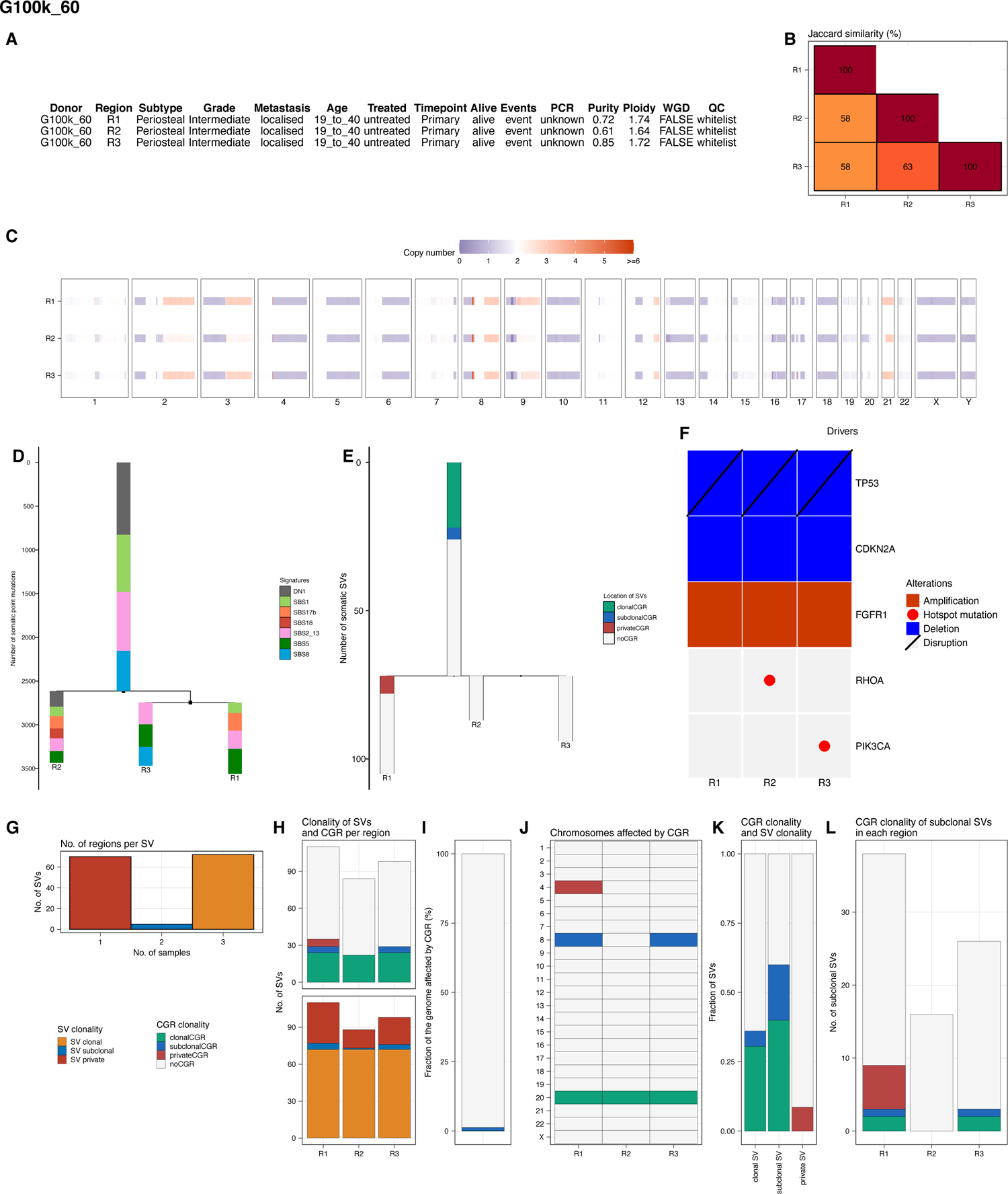

**Fig. S59.**
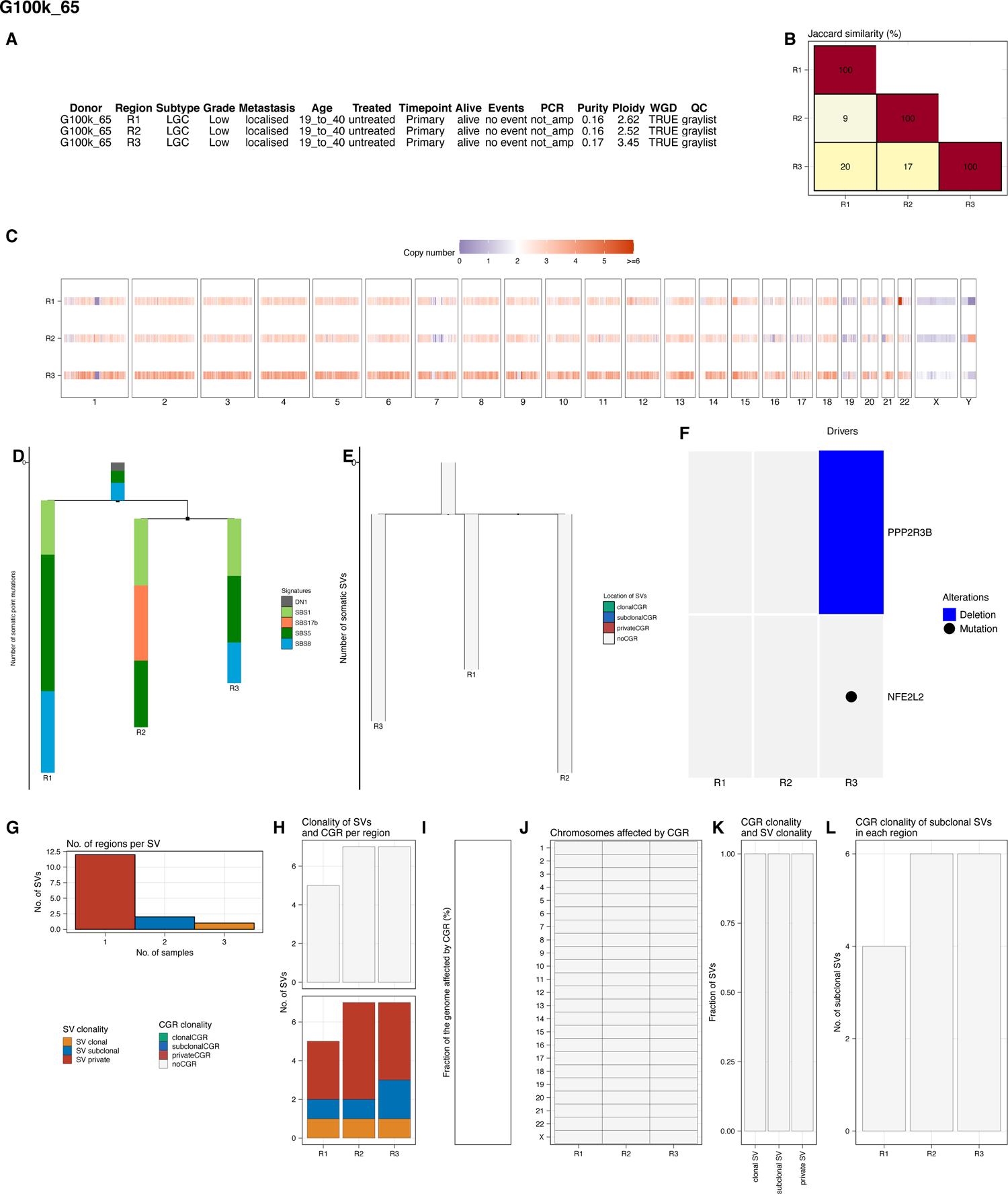

**Fig. S60.**
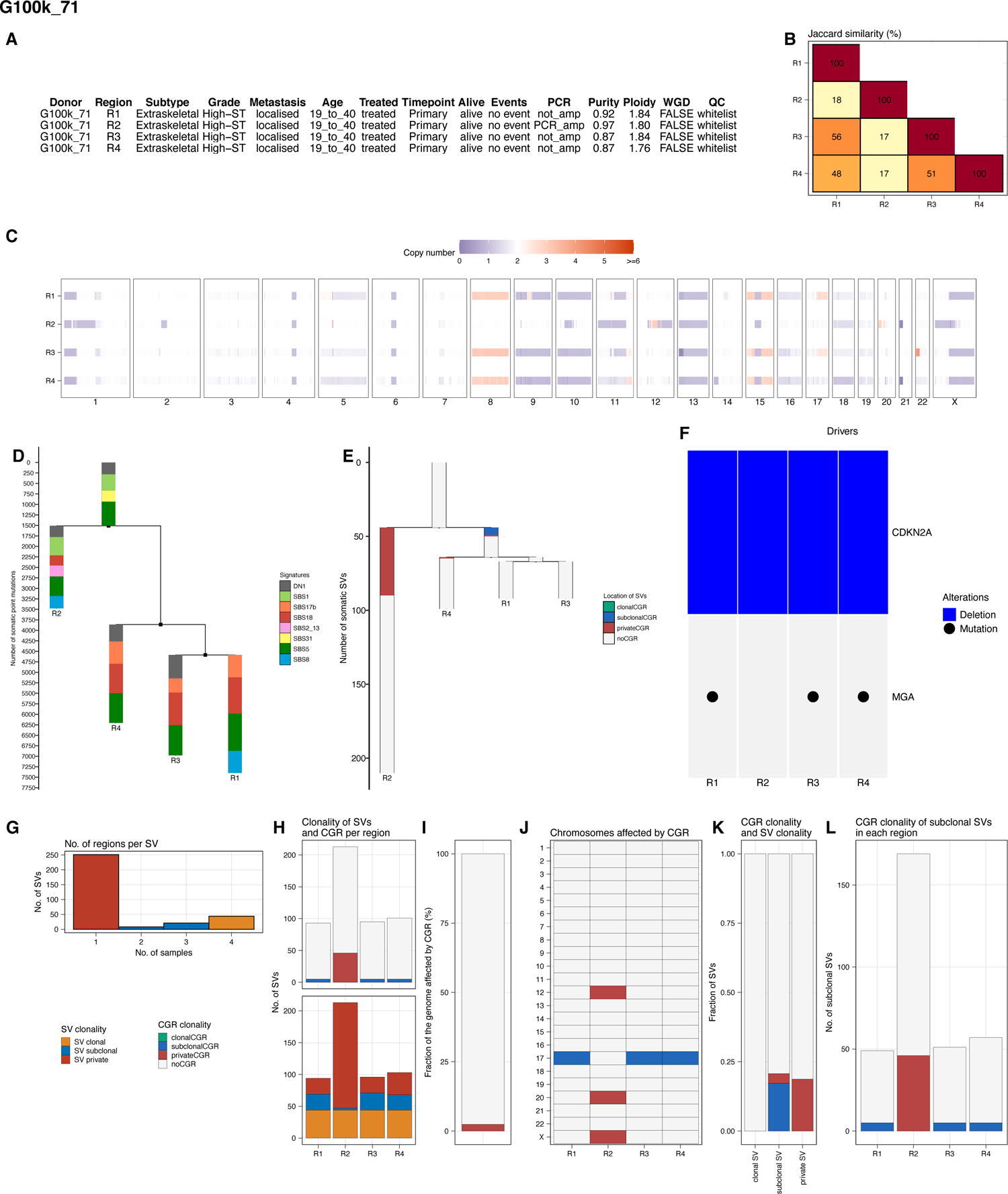

**Fig. S61.**
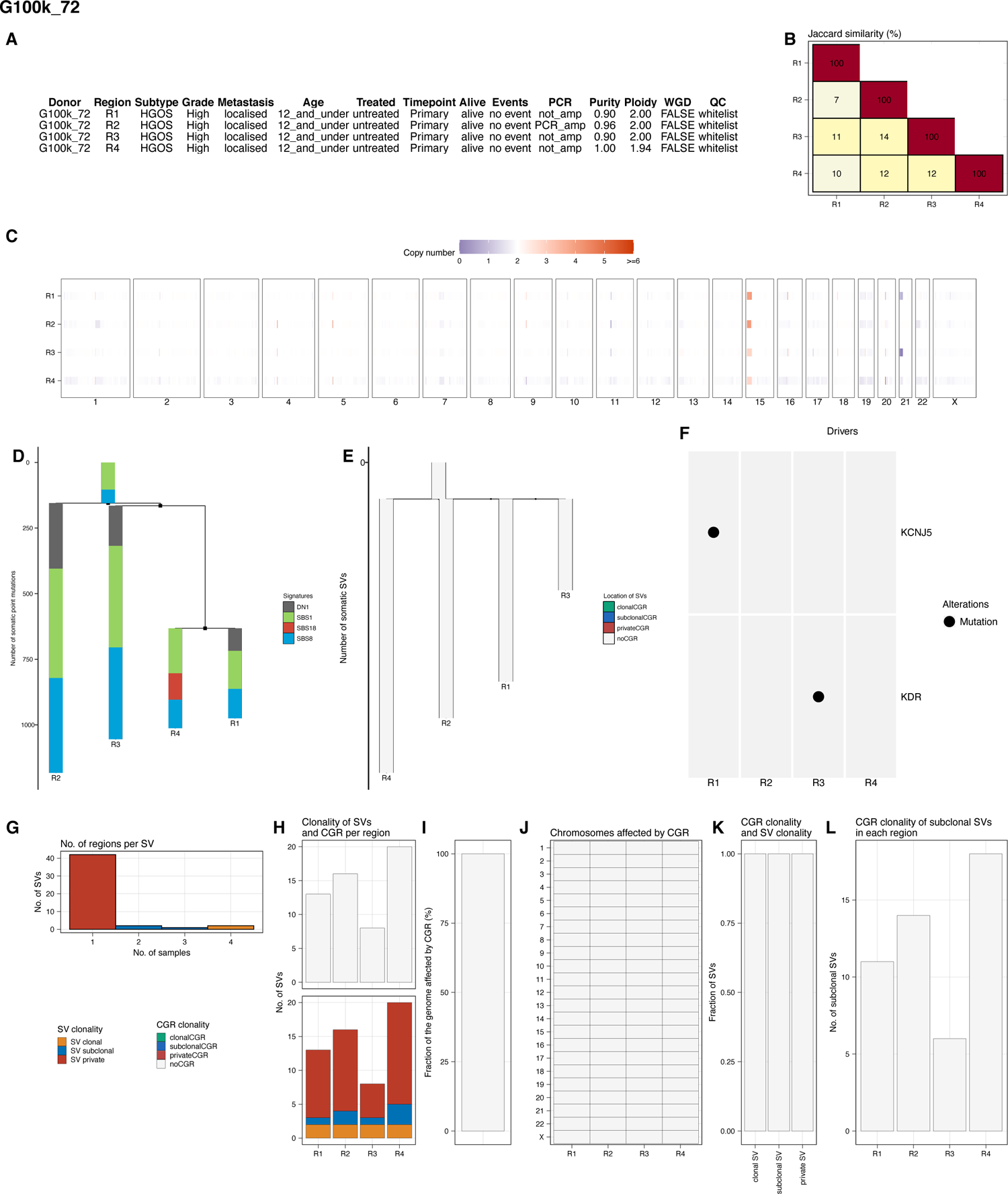

**Fig. S62.**
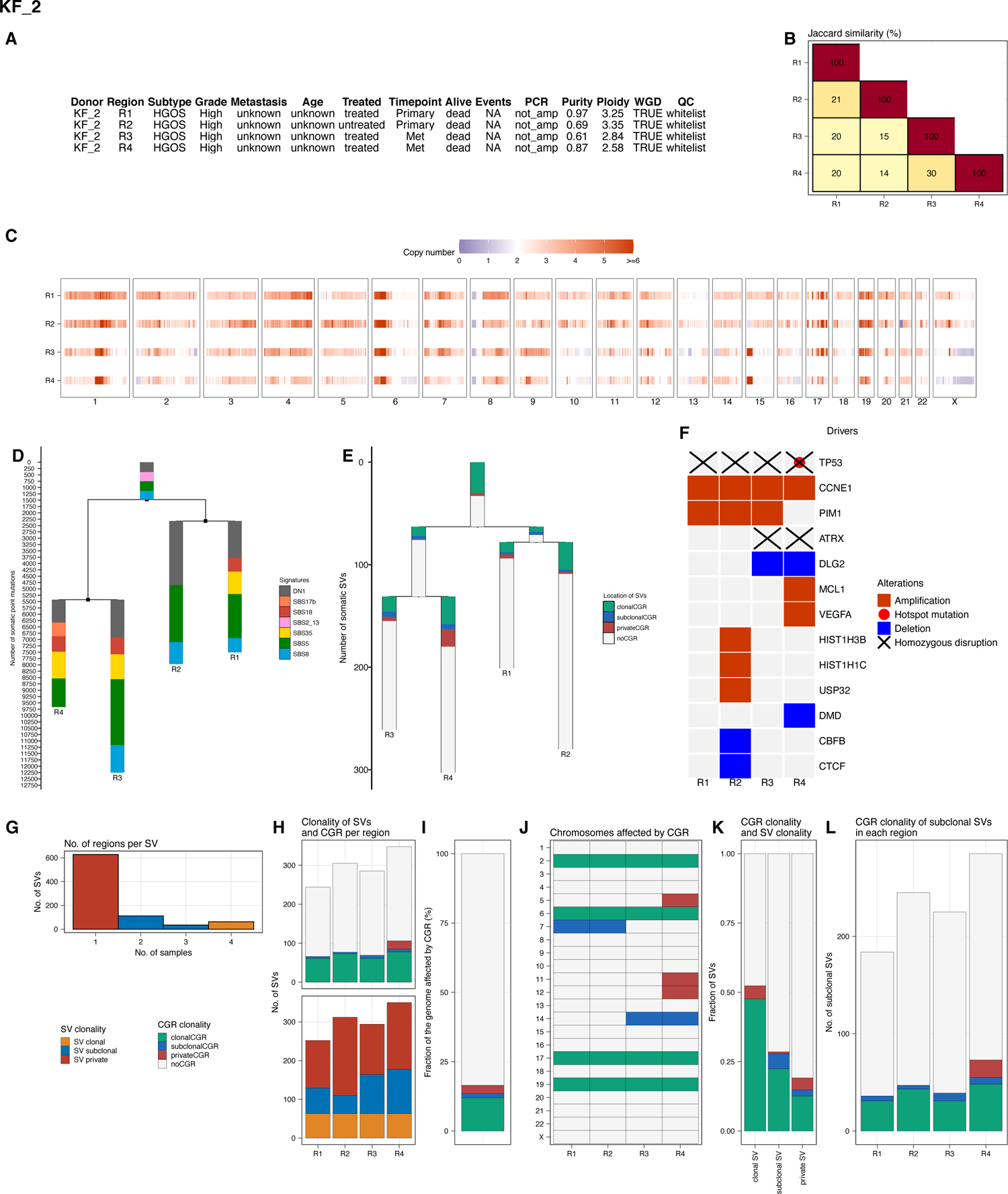

**Fig. S63.**
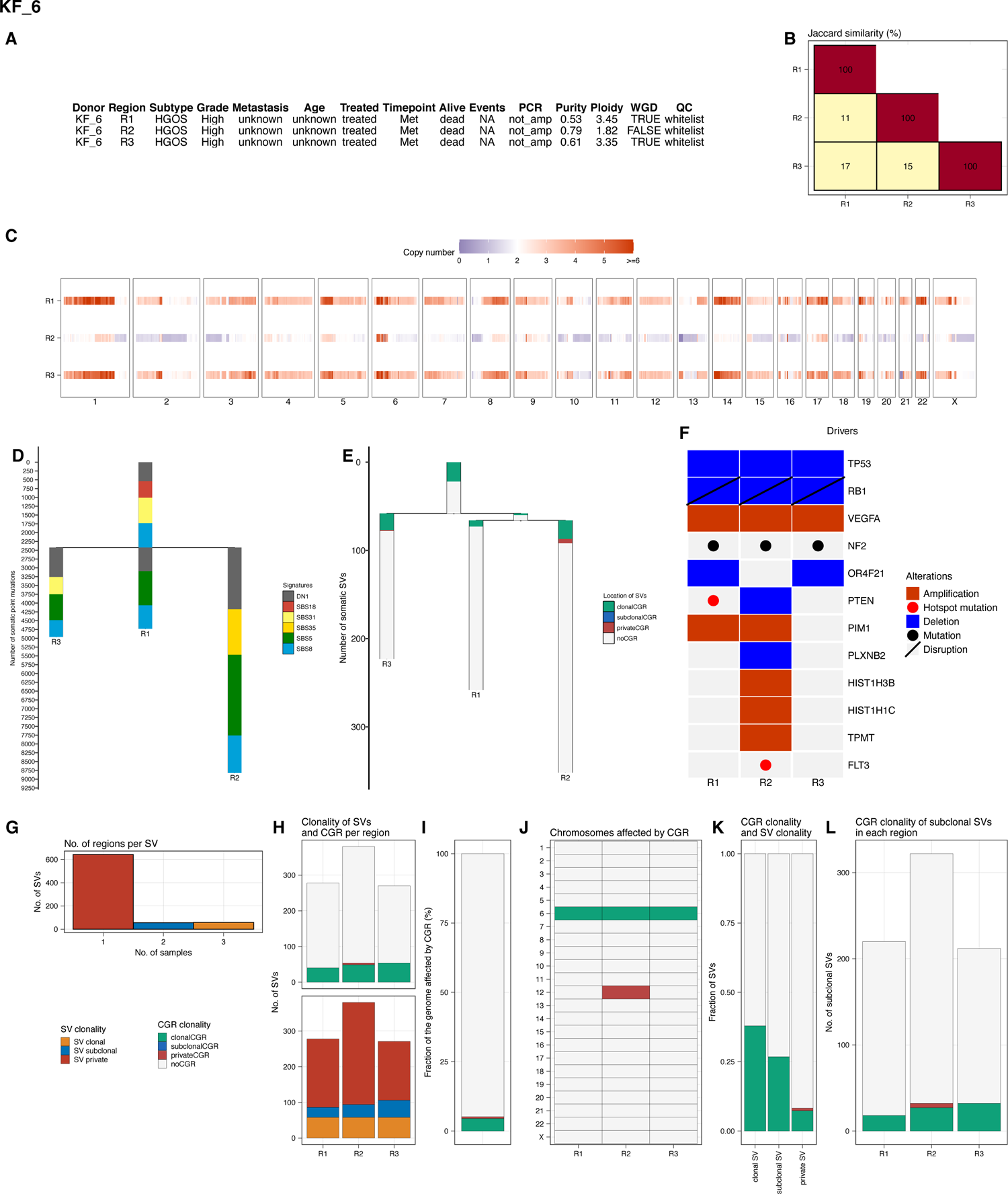

**Fig. S64.**
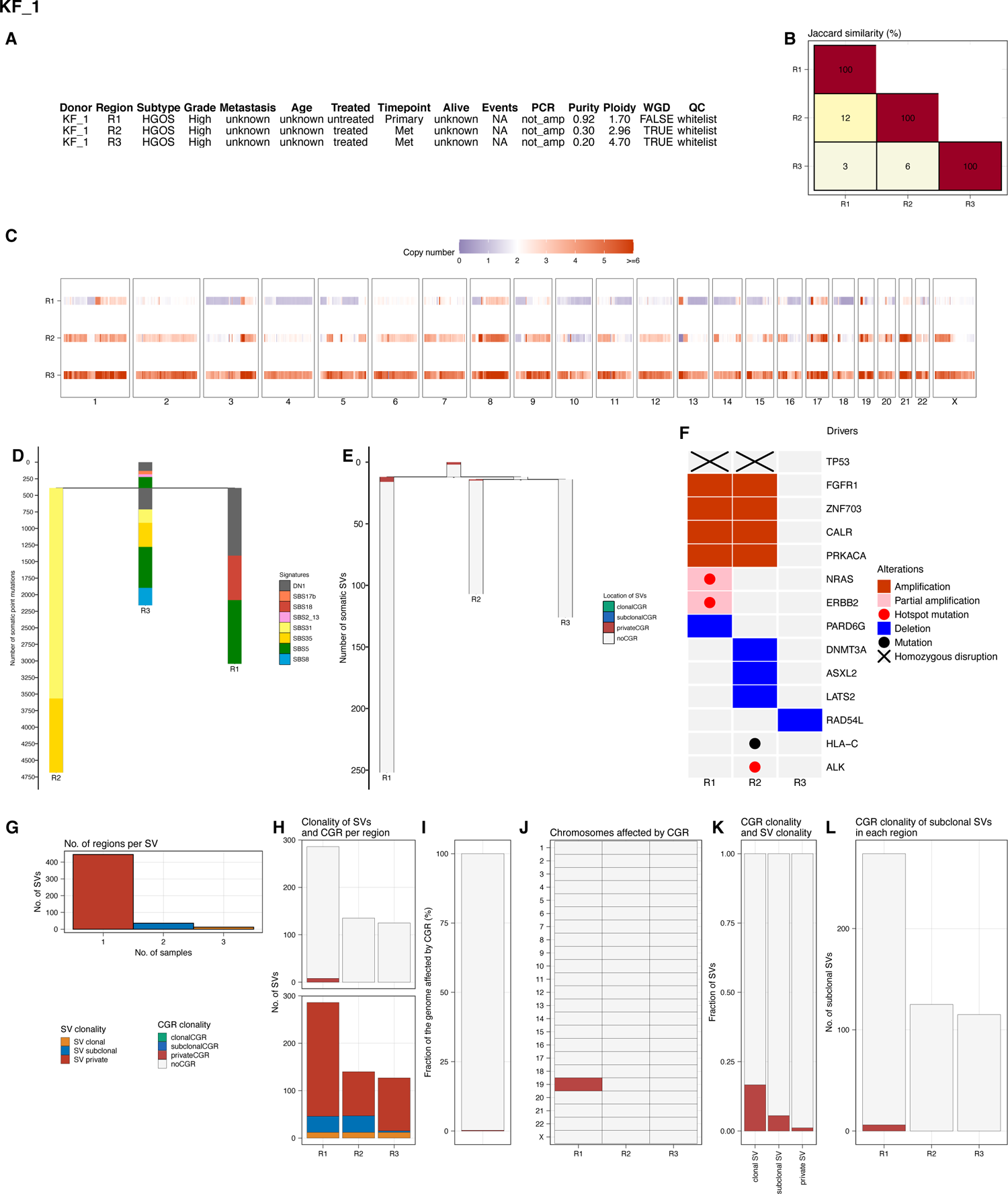

**Fig. S65.**
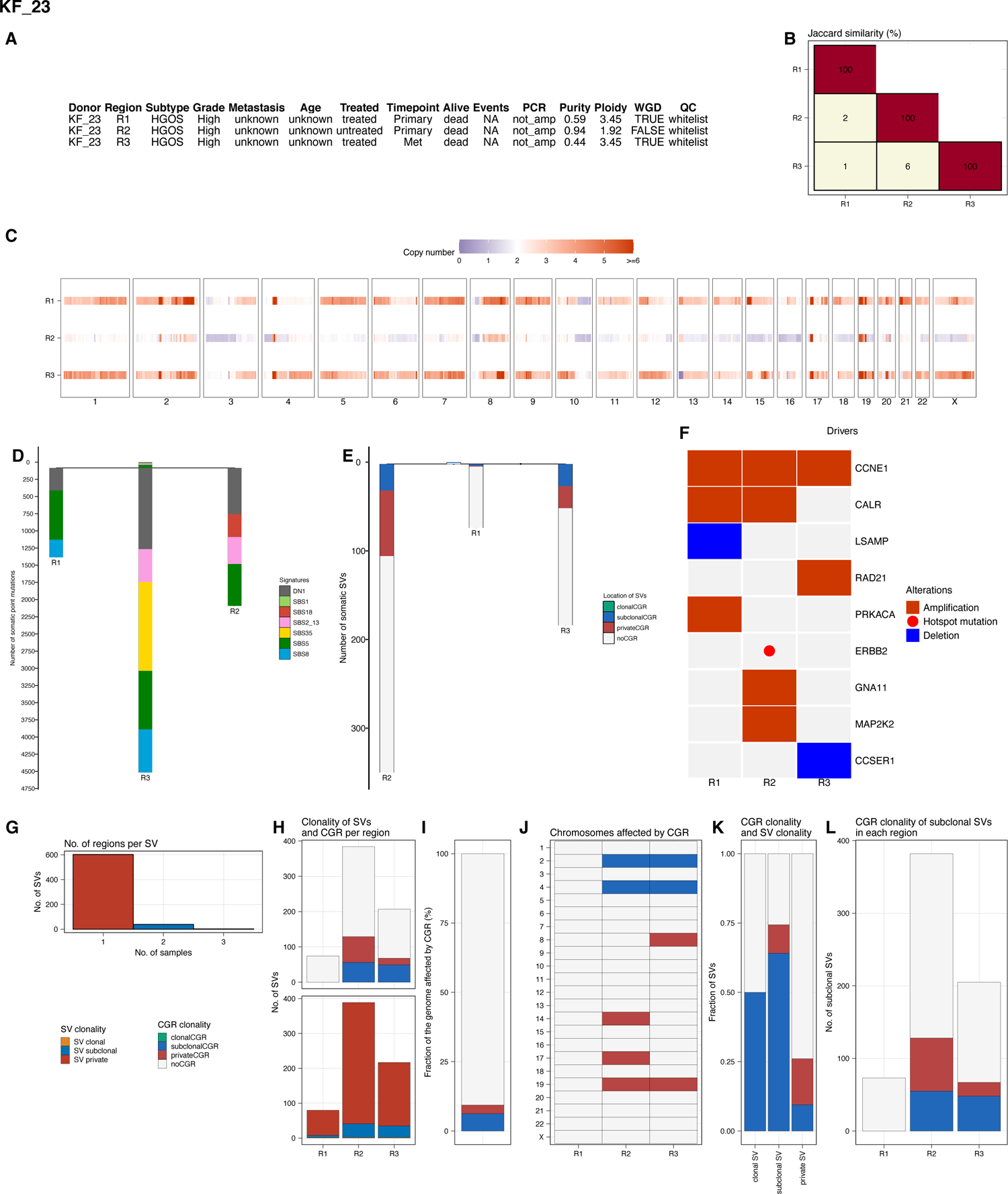

